# Neoantigen Cancer Vaccines and Different Immune Checkpoint Therapies Each Utilize Both Converging and Distinct Mechanisms that in Combination Enable Synergistic Therapeutic Efficacy

**DOI:** 10.1101/2023.12.20.570816

**Authors:** Sunita Keshari, Alexander S. Shavkunov, Qi Miao, Akata Saha, Charmelle D. Williams, Anna M. Highsmith, Josué E. Pineda, Elise Alspach, Kenneth H. Hu, Kristen E. Pauken, Ken Chen, Matthew M. Gubin

## Abstract

The goal of therapeutic cancer vaccines and immune checkpoint therapy (ICT) is to eliminate cancer by expanding and/or sustaining T cells with anti-tumor capabilities. However, whether cancer vaccines and ICT enhance anti-tumor immunity by distinct or overlapping mechanisms remains unclear. Here, we compared effective therapeutic tumor-specific mutant neoantigen (NeoAg) cancer vaccines with anti-CTLA-4 and/or anti-PD-1 ICT in preclinical models. Both NeoAg vaccines and ICT induce expansion of intratumoral NeoAg-specific CD8 T cells, though the degree of expansion and acquisition of effector activity was much more substantial following NeoAg vaccination. Further, we found that NeoAg vaccines are particularly adept at inducing proliferating and stem-like NeoAg-specific CD8 T cells. Single cell T cell receptor (TCR) sequencing revealed that TCR clonotype expansion and diversity of NeoAg-specific CD8 T cells relates to their phenotype and functional state associated with specific immunotherapies employed. Effective NeoAg vaccines and ICT required both CD8 and CD4 T cells. While NeoAg vaccines and anti-PD-1 affected the CD4 T cell compartment, it was to less of an extent than observed with anti-CTLA-4, which notably induced ICOS^+^Bhlhe40^+^ Th1-like CD4 T cells and, when combined with anti-PD-1, a small subset of Th2-like CD4 T cells. Although effective NeoAg vaccines or ICT expanded intratumoral M1-like iNOS^+^ macrophages, NeoAg vaccines expanded rather than suppressed (as observed with ICT) M2-like CX3CR1^+^CD206^+^ macrophages, associated with the vaccine adjuvant. Further, combining NeoAg vaccination with ICT induced superior efficacy compared to either therapy in isolation, highlighting the utility of combining these modalities to eliminate cancer.

**Highlights:**

- NeoAg cancer vaccines utilize distinct mechanisms from αCTLA-4 or αPD-1 ICT
- NeoAg vaccines induce TCF1^+^ stem-like and proliferating NeoAg-specific CD8 T cells
- CD8 TCR clonotype expansion relates to phenotype and functional state associated with immunotherapy
- NeoAg vaccines induce partially distinct macrophage remodeling from ICT
- NeoAg vaccines synergize with ICT, exceeding combination αCTLA-4/αPD-1 ICT efficacy

## INTRODUCTION

For cancer immunotherapies such as immune checkpoint therapy (ICT), T cell recognition of tumor antigens is critical for efficacy^1–4^. In contrast to aberrantly expressed non-mutant tumor antigens, tumor-specific neoantigens (NeoAgs) formed from somatic alterations in cancer cells are largely excluded from immune tolerance and are exclusively expressed in cancer cells, making them favorable cancer vaccine targets^2–4^. Significant progress has been made in the field of NeoAg cancer vaccine development, showing promise in early-phase clinical trials^5–12^. Despite this, many fundamental questions regarding NeoAg vaccines remain unclear, including how to best combine therapeutic vaccines with other T cell-directed therapeutic modalities including ICT to promote optimal outcomes in cancer patients.

We previously used immunogenomic/mass spectrometry approaches to identify NeoAgs and subsequently demonstrated that therapeutic NeoAg cancer vaccines could provoke tumor rejection in methylcholanthrene (MCA)-induced sarcoma models^13^. Others have used similar approaches to identify immunogenic NeoAgs^4,6,7,14–17^. We further showed that NeoAgs are major targets of T cells reactivated by ICT and that anti-CTLA-4 and anti-PD-1 ICT induces changes in both CD4 and CD8 T cells within the tumor microenvironment (TME)^13,18–21^, consistent with findings from others^22,23^. While both conventional CD4 and CD8 T cells drive immunotherapeutic responses to cancer, CD8 T cells are often the most potent direct inducers of tumor cell death^24^. In both cancer patients and preclinical models, intratumoral CD8 T cells that express activation markers including inhibitory receptors such as PD-1, LAG-3, and TIM-3 often exist in a terminally differentiated state and may display a range of functional capabilities from short-lived cytotoxic and cytokine producing CD8 T effector cells to dysfunctional or exhausted CD8 T cells that exist in a state of limited or restrained functional capabilities^25^. These dysfunctional or exhausted CD8 T cells exist on a spectrum and may progress from intermediate dysfunctional/exhausted to terminal dysfunctional/exhausted CD8 T cells characterized by high, sustained expression of inhibitory receptors, reduced function, and unique transcriptional and epigenetic profiles. These features differentiate dysfunctional/exhausted CD8 T cells from memory T cells and T cells displaying stem-like properties (often referred to as progenitor/precursor exhausted CD8 T cells). These distinct states are driven by key transcription factors, including TCF-1, which promotes stemness or memory-like attributes^26,27^, and TOX, which plays a crucial role in establishing terminal dysfunction/exhaustion^28–30^. Chronic antigen exposure and/or signals within the TME promote maintenance of NFAT-independent TOX expression and establishment of a fixed epigenetic landscape in terminal dysfunctional/exhausted CD8 T cells^31^. The increased presence of PD-1^hi^ TOX^+^ TCF-1^−^ CD8 T cells in tumor biopsies correlates with a poorer prognosis in patients treated with ICT and these cells likely lack the ability to gain significant effector function following PD-1/PD-L1 blockade^32,33^. Instead, stem-like PD-1^+^ Tim-3^−^ TCF-1^+^ CD8 T cells within tumors and lymph nodes expand and differentiate into PD-1^+^ Tim-3^+^ CD8 T effector-like cells in response to anti-PD-1/PD-L1 ICT^25,34–37^.

While T cells are the major target of NeoAg vaccines and ICT, myeloid cells are a critical component of the TME^38^. Macrophages are amongst the most abundant intratumoral myeloid cell population and may comprise both embryonically-derived tissue-resident macrophages and monocyte-derived macrophages, with the latter accounting for a majority of macrophages present at diseased sites^39–41^. We previously observed major complexity in the ICT-induced changes occurring in the intratumoral macrophage compartment, despite T cells being the predominant direct target of ICT^19–21^. These changes included remodeling from M2-like CX3CR1^+^CD206^+^ macrophages in progressively growing tumors to M1-like iNOS^+^ macrophages in tumors that go on to reject in response to ICT. Further, blockade of TREM2 expressed on macrophages induced a decline in CX3CR1^+^CD206^+^ macrophages and promoted macrophages expressing immunostimulatory molecules, with anti-TREM2 monoclonal antibody (mAb) dampening tumor growth and augmenting anti-PD-1 efficacy^42^.

Tumor immune cell compositions clearly play a major role in response to immunotherapy^43,44^, but the heterogeneity and dynamics of immune infiltrates in response to immunotherapies such as NeoAg cancer vaccines is not thoroughly characterized. Further, although much progress has been made towards defining the mechanisms behind ICT efficacy, our understanding is still incomplete and direct comparisons between cancer vaccines and different ICTs used alone or in combination are largely lacking. A more refined understanding of how NeoAg vaccines impact the immune TME in comparison to other immunotherapies can inform rational use of NeoAg vaccines and combinatorial immunotherapies.

To address this, we developed preclinical models to interrogate potential synergies between the mechanisms underlying NeoAg cancer vaccines and different ICTs. We systematically compared different immunotherapies that lead to tumor rejection, including NeoAg cancer vaccines, anti-CTLA-4, anti-PD-1, and anti-CTLA-4 + anti-PD-1 ICT using mouse melanoma models expressing defined NeoAgs. NeoAg vaccines induced the most robust expansion of polyfunctional NeoAg-specific CD8 T cells, including proliferating and stem-like CD8 T cells. Further, NeoAg-specific CD8 TCR clonotype expansion and diversity of NeoAg-specific CD8 T cells related to their phenotype and functional state associated with specific immunotherapies used. Anti-CTLA-4 and/or anti-PD-1 ICT increased the frequency and effector function of intratumoral NeoAg-specific CD8 T cells, with anti-CTLA-4 containing treatments also dramatically altering the CD4 T cell compartment. Both NeoAg vaccines and ICT resulted in an expansion of M1-like iNOS^+^ macrophages and while ICT reduced the frequency of intratumoral CX3CR1^+^CD206^+^ M2-like macrophages, CX3CR1^+^CD206^+^ macrophages were largely maintained in NeoAg vaccine treated mice. To investigate whether the unique impacts of NeoAg vaccines and ICT combine for enhanced tumor control, we tested the efficacy of NeoAg vaccination in combination with either anti-CTLA-4 or anti-PD-1 and found that the window of therapeutic efficacy was extended by combination treatments, further supporting the rationale of combining NeoAg vaccines with ICT.

## RESULTS

### NeoAg vaccines and ICT induce T cell-dependent long-term tumor protection

For this study, we modified the genetically engineered mouse model (GEMM)-derived *Braf^V600E^ Pten^-/-^ Cdkn2a^-/-^* YUMM1.7 mouse melanoma line^45^ to express different combinations of MHC-I and MHC-II NeoAgs. GEMM tumors are generally poorly immunogenic; however, they can be engineered to express NeoAgs to study tumor-immune interactions^20,46–49^. We engineered YUMM1.7 to express known tumor antigens via introduction of minigenes encoding the G1254V mutation in Laminin subunit alpha 4 (mLama4^MHC-I^), the A506T mutation in Alpha-1,3-glucosyltransferase (mAlg8^MHC-I^), and the N710Y mutation in Integrin beta 1 (mItgb1^MHC-II^) NeoAgs^13,20^ in various combinations: mLama4^MHC-I^ + mItgb1^MHC-II^ (Y1.7LI line) or mAlg8^MHC-I^ + mItgb1^MHC-II^ (Y1.7AI line) **(Figure S1A)**. Consistent with prior observations^45,50^, the parental YUMM1.7 melanoma line was insensitive to anti-CTLA-4 and/or anti-PD-1 ICT **(Figure S1B)**. In contrast, enforced expression of mLama4^MHC-I^ or mAlg8^MHC-I^ NeoAg along with mItgb1^MHC-II^ NeoAg rendered YUMM1.7 melanoma lines (Y1.7LI and Y1.7AI) sensitive to anti-CTLA-4 ICT **(Figure 1A)**.

**Figure 1.**
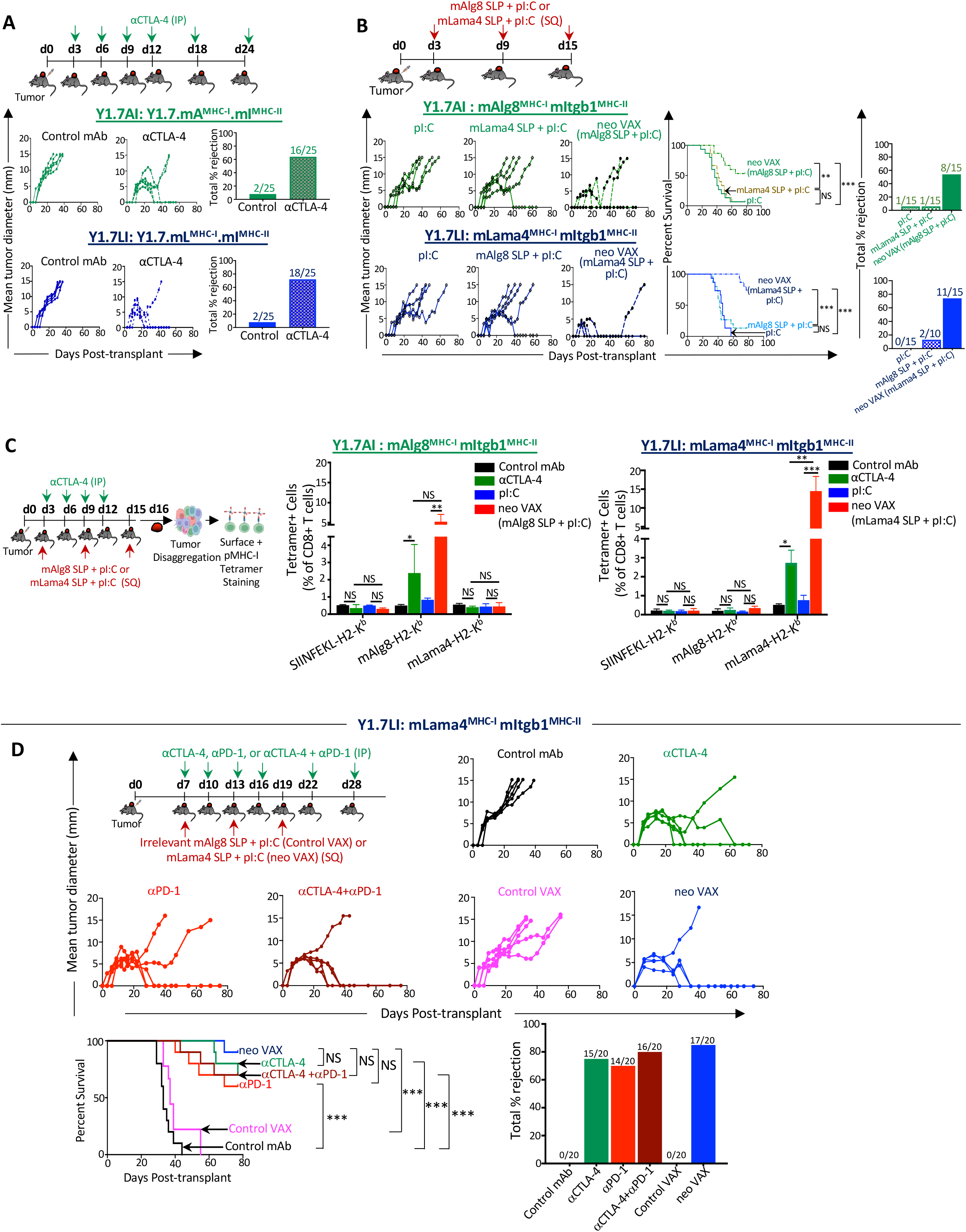
Therapeutic NeoAg vaccines or ICT inhibit NeoAg-expressing *Braf^V600E^ Pten^-/-^ Cdkn2a^-/-^* melanoma growth. **(A)** Tumor growth and percent tumor rejection in wildtype (WT) C57BL/6J mice transplanted with Y1.7 mA^MHC-I^.mI^MHC-II^ (Y1.7AI) and Y1.7 mL^MHC-I^.mI^MHC-II^ (Y1.7LI) melanoma cells and treated with control mAb or anti-CTLA-4 immune checkpoint therapy (ICT) starting on d. 3 post tumor-transplant, and subsequently on d. 6, 9, 12, 18, 24. **(B)** Tumor growth, cumulative mouse survival, and percent tumor rejection in WT C57BL/6J mice transplanted with Y1.7AI and Y1.7LI melanoma cells and treated with mAlg8 or mLama4 NeoAg synthetic long peptide (SLP) plus poly I:C (pI:C) vaccines or pI:C alone starting on d. 3 post tumor-transplant and given every 6 days for 3 total doses. **(C)** Bar graphs displaying mAlg8 or mLama4 tetramer-specific CD8 T cells in Y1.7AI and Y1.7LI tumors treated with control mAb, anti-CTLA-4, pI:C, mAlg8 SLP + pI:C NeoAg vaccine (for Y1.7AI) or mLama4 SLP + pI:C NeoAg vaccine (for Y1.7LI) as in **(A)** and **(B)** and harvested on d. 16 post-tumor transplant. SIINFEKL-H2-K^b^ tetramer served as irrelevant control. **(D)** Tumor growth, cumulative mouse survival, and percent tumor rejection in WT C57BL/6J mice transplanted with Y1.7LI melanoma cells and treated with control mAb, anti-CTLA-4, anti-PD-1, anti-CTLA-4 + anti-PD-1, irrelevant (for Y1.7LI) mAlg8 SLP + pI:C (control VAX), or relevant mLama4 SLP + pI:C (neo VAX) starting on d. 7 post tumor-transplant, and subsequently on d. 10, 13, 16, 22, 28 for ICT and d. 13, 19 for NeoAg vaccines. Tumor growth data in **(A)**, **(B)**, and **(D)** are presented as individual mouse tumor growth as mean tumor diameter and are representative of **(A)** five, **(B)** three, or **(D)** four independent experiments. Tumor rejection graphs display cumulative percentage of mice with complete tumor rejection from independent experiments. Cumulative survival curves and tumor rejection graphs include mice from three independent experiments (\*\**P* < *0.01*, \*\*\**P* < *0.001*, log-rank (Mantel–Cox) test). Bar graphs in **(C)**, display mean ± SEM and are representative of at least three independent experiments (\**P* < 0.05, \*\**P* < 0.01, \*\*\**P* < 0.005, NS, not significant; unpaired, two-tailed Student’s *t* test). See also **Figure S1**.

We next asked whether therapeutic cancer vaccines composed of the synthetic long peptide (SLP) containing the minimal MHC-I NeoAg epitope and the adjuvant poly I:C (pI:C) could induce regression of the Y1.7LI and Y1.7AI NeoAg-expressing lines. Tumor bearing mice treated with pI:C alone displayed outgrowth of Y1.7LI or Y1.7AI melanoma, whereas vaccines comprising relevant NeoAg SLP + pI:C (neo VAX) induced complete rejection or delayed outgrowth of both Y1.7 NeoAg expressing variants **(Figure 1B)**. NeoAg vaccine-induced tumor rejection was dependent upon specific NeoAg expression, as mAlg8 SLP + pI:C did not induce Y1.7LI (mLama4-expressing) tumor rejection and vice versa with Y1.7AI (mAlg8-expressing) **(Figure 1B)**. Mice that rejected Y1.7AI or Y1.7LI tumors upon neo VAX or anti-CTLA-4 were rechallenged in the absence of any additional treatment with the same tumor lines at least 60 days after rejection of primary tumors. Upon secondary challenge, no detectable tumor was observed indicating long-term protection against rechallenge with the same tumor **(Figure S1C)**. In contrast, both Y1.7-NeoAg expressing lines grew out when injected into naïve mice in the absence of treatment, indicating cell line preparations used in rechallenge experiments were capable of tumor formation. When mice that previously rejected Y1.7LI tumors upon were rechallenged with parental YUMM1.7, progressive tumor growth was observed **(Figure S1D)**, indicating immunity was NeoAg-specific.

We next used peptide-MHC (pMHC) tetramers to detect intratumoral CD8 T cells recognizing the mLama4 or mAlg8 NeoAg presented on H-2K^b^. Tumors from anti-CTLA-4 treated mice contained greater frequencies of mAlg8- or mLama4-specific CD8 T cells compared to mice receiving control mAb **(Figures 1C and S1E)**. Whereas pI:C alone had little effect on the frequency of NeoAg-specific CD8 T cells, neo VAX induced an over 5-fold or more increase in mAlg8- or mLama4-specific CD8 T cells **(Figures 1C and S1E)**. Neo VAX also significantly increased the frequency of NeoAg-specific CD8 T cells co-expressing the inhibitory receptors PD-1 and TIM-3 **(Figure S1F)**, although this does not necessarily indicate reduced function and may instead reflect antigen stimulation and T cell activation state^24,51^.

To expand on these observations, we focused on the Y1.7LI line and delayed treatment initiation until day 7 post-transplant. Y1.7LI tumor bearing mice treated with control mAb or control VAX (irrelevant mAlg8 SLP + pI:C) starting on day 7 displayed progressive tumor outgrowth **(Figure 1D)**. In contrast, anti-CTLA-4, anti-PD-1, combination anti-CTLA-4 plus anti-PD-1, or neo VAX induced tumor rejection in a majority of mice. ICT- and neo VAX-induced tumor rejection was dependent on both CD4 and CD8 T cells, as mAb depletion of either T cell subset abolished therapeutic efficacy **(Figure S2A)**. Y1.7LI-rechallenged mice that rejected Y1.7LI tumors upon neo VAX or anti-CTLA-4 and/or anti-PD-1 initiated on day 7, but not untreated naïve mice, showed no detectable tumor upon secondary challenge **(Figure S2B)**.

### Tumor microenvironment remodeling induced by NeoAg vaccines and ICT

We next used an unbiased approach to assess whether effective tumor-specific NeoAg vaccines induced TME alterations that are distinct or overlapping with different forms of ICT. Y1.7LI tumor bearing mice were treated with (1) control mAb, (2) anti-CTLA-4, (3) anti-PD-1, (4) anti-CTLA-4 + anti-PD-1, (5) control VAX (irrelevant SLP + pI:C), or (6) neo VAX (mLama4 SLP + pI:C) beginning on day 7 **(Figure 2A)**. Tumors were harvested on day 15 (a critical timepoint prior to tumor rejection during ICT or neo VAX in this model) and live CD45^+^ cells were sorted for scRNAseq. We used unsupervised graph-based clustering to stratify myeloid cells and lymphocytes (**Figures 2B and 2C)**. scRNAseq and flow cytometry both indicated that immunotherapy altered the proportions of different myeloid and lymphoid subsets **(Figure S3A)**.

**Figure 2.**
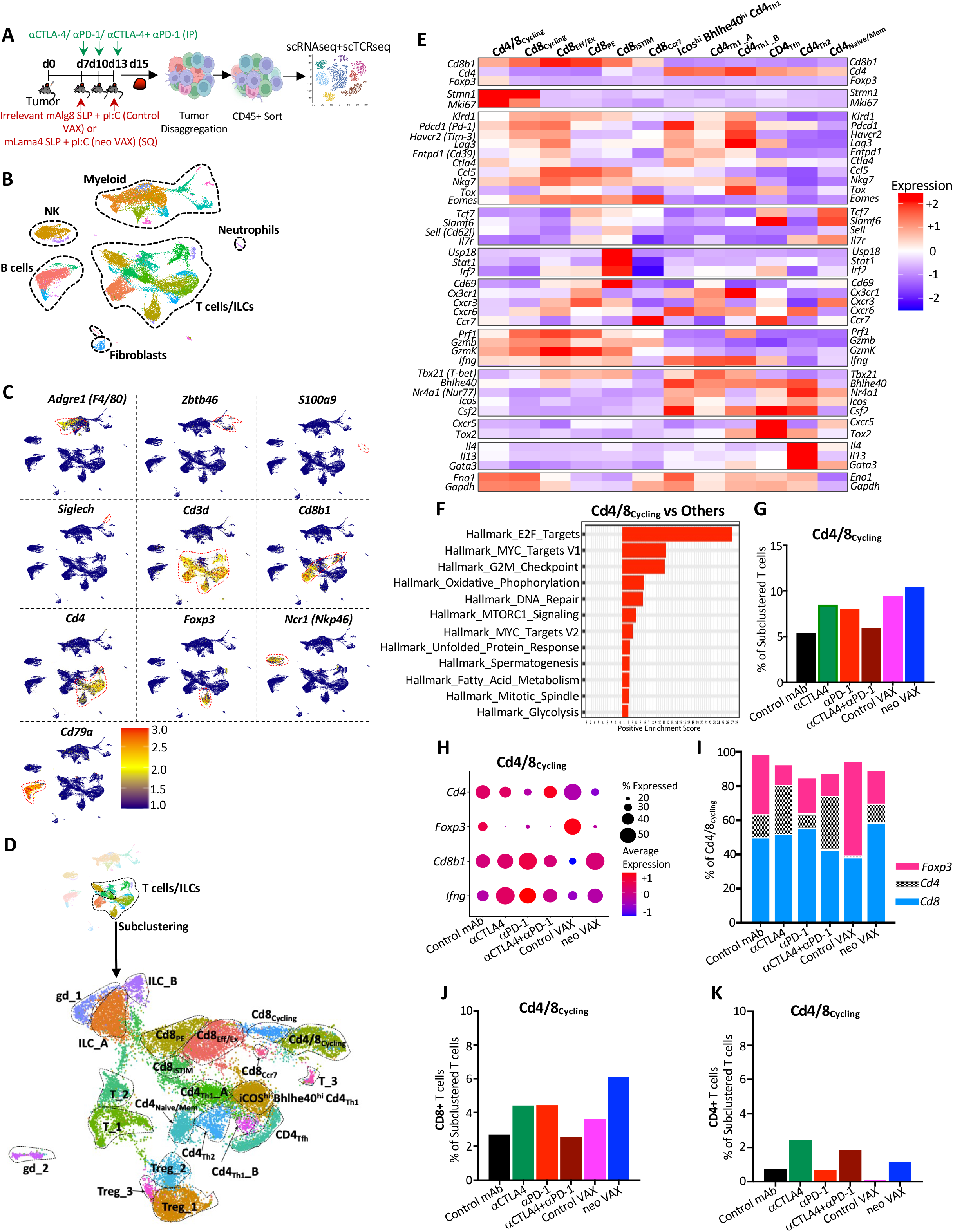
scRNAseq of intratumoral immune cells from Y1.7LI tumor bearing mice treated with NeoAg vaccines or ICT. **(A)** Experimental setup for **(B)-(K)**. WT C57BL/6J mice were injected with Y1.7LI melanoma cells and subsequently treated beginning on d. 7 with control mAb, anti-CTLA-4, anti-PD-1, anti-CTLA-4 + anti-PD-1, irrelevant (for Y1.7LI) mAlg8 SLP + pI:C (control VAX), or relevant mLama4 SLP + pI:C (neo VAX) and harvested on d. 15 post-tumor transplant. Intratumoral live CD45^+^ cells were sorted and analyzed by scRNAseq. **(B)** UMAP plot from scRNAseq of intratumoral CD45^+^ cells with annotated cell types. **(C)** Feature plot showing lineage-specific transcripts defining lymphoid and myeloid cell types. **(D)** Feature plots displaying subclustering of activated T cell-containing clusters, subclustered T cell/ILC cluster annotations (middle plot), and *Cd4* and *Cd8* expression (bottom plot). **(E)** Heat map displaying average expression of select transcripts by cluster. **(F)** Gene set enrichment analysis (GSEA) displaying significantly enriched gene sets in cluster Cd4/8_Cycling_. **(G)** Proliferating T cells in cluster Cd4/8_Cycling_ by treatment condition represented as percentage of subclustered T cells. **(H)** Dot plot depicting expression level and percent of cells expressing *Foxp3*, *Cd4*, *Cd8*, *Ifng* in Cd4/8_Cycling_ by treatment condition. **(I)** Percentage of Foxp3^+^ Tregs, conventional CD4 T cells, or CD8 T cells in Cd4/8_Cycling_ by treatment condition. **(J)** Graph displaying CD8 T cells from cluster Cd4/8_Cycling_ represented as percentage of total subclustered T cells. **(K)** Graph displaying conventional CD4 T cells from cluster Cd4/8_Cycling_ represented as percentage of total subclustered T cells. See also **Figures S4** and **S5**.

To gain more insights into how the different immunotherapies altered T cells in the TME, we chose clusters containing activated T cells for subclustering and identified multiple clusters of conventional CD4 and CD8 T cells, Foxp3^+^ CD4^+^ T regulatory cells (Tregs), gamma delta T cells (γοT), and innate lymphoid cells (ILCs) **(Figures 2D, S3B-S3E, S4, and S5)**.

While most clusters contained either CD4 or CD8 T cells, cluster Cd4/8**_Cycling_** contained a mix of Tregs, CD4 T cells, and CD8 T cells and displayed a cell proliferation transcriptional signature **(Figures 2D-2F, S4 and S5)**. Not only did tumors from neo VAX, anti-CTLA-4, or anti-PD-1 treated mice have a greater frequency of cells within Cd4/8**_Cycling_**, but the ratio of cycling conventional CD4 and CD8 T cells to Tregs was higher as compared to control mAb or control VAX **(Figures 2G-2K)**.

Anti-CTLA-4 (+/- anti-PD-1) reduced proliferating Tregs and expanded CD4 T cells within Cd4/8**_Cycling_**, while the ratio of proliferating CD8 T cells to Tregs or CD4 T cells was higher with anti-PD-1. Interestingly, neo VAX contained the greatest ratio of cycling CD8 T cells to other T cells in this cluster **(Figures 2H-J)**.

Although this analysis did not distinguish their antigen specificity, we identified 5 exclusively CD8 T cell clusters, spanning a range of activation states including proliferating (Cd8**_Cycling_**), CD69^hi^ IFN stimulated [Cd8**_iSTIM_** (interferon <u>STIM</u>ulated)], PD-1^+^ TCF7^+^ plastic/stem-like or progenitor exhausted (Cd8**_PE_**), and PD-1^+^ TCF7^−^ terminal effectors or dysfunctional/exhausted CD8 T cells (Cd8**_Eff/Ex_**) **(Figures 2D, 2E, S4, S5, and S6A-S6F)**. Cd8_Cycling_ exhibited features of proliferation/cycling but was exclusively composed of CD8 T cells which displayed a more activated phenotype compared to Cd4/8**_Cycling_ (Figures S4, S5, S6A, and S6B)**. Whereas the percentage of Cd8**_Cycling_** cells increased modestly with anti-CTLA-4 or anti-PD-1, neo VAX drove ∼2-fold increase in the frequency of cells within this cluster **(Figure S6B)**, thus indicating that neo VAX more robustly expands subsets of proliferating CD8 T cells.

Cluster Cd8**_Eff/Ex_** expressed little detectable *Tcf7* (encoding TCF-1) and displayed elevated transcript expression of multiple inhibitory receptors (e.g., *Pdcd1* (PD-1), *Havcr2* (TIM-3), *Lag3*) and other genes associated with T cell activation, effector function, and also exhaustion/dysfunction including *Tox* **(Figures S5, S6A, and S6C)**. Cd8**_PE_** expressed *Pdcd1*, but to less of an extent than Cd8**_Eff/Ex_**, and additionally expressed *Slamf6* and *Tcf7,* indicating a phenotype consistent with progenitor/precursor exhausted T cells that display plastic/stem-like properties **(Figures S5, S6A, and S6D)**. neo VAX, anti-CTLA-4, or anti-PD-1 reduced the fraction of cells within Cd8**_Eff/Ex_** and Cd8**_PE_** with combination anti-CTLA-4 and anti-PD-1 standing out as the only treatment to not decrease the frequency of Cd8**_Eff/Ex_ (Figures S6C and S6D)**.

Within Cd8**_Cycling_**, Cd8**_PE_**, Cd8**_iSTIM_**, and Cd8**_Ccr7_**, the highest expression of *Lag3*, *Cd39*, and *Gzmb* within each respective cluster was observed with combination anti-CTLA-4 + anti-PD-1 ICT **(Figures S5, S6A, S6B, and S6D-S6F)**. Additionally, *Prf1* was most robustly induced by combination ICT in all CD8 clusters, except for Cd8**_Ccr7_**, where neo VAX induced the highest expression **(Figures S5 and S6A-S6F)**. Further, a pattern emerged within CD8 T cells whereby in each cluster, anti-CTLA-4 (alone or in combination with anti-PD-1), as well as neo VAX to some extent, drove higher expression of *Cd226* encoding the co-activating receptor CD226/DNAM-1. CD226 counteracts the actions of the inhibitory receptor TIGIT by competing for binding to ligands such as CD155^52^. Expression of *Tigit* followed an inverse pattern as *Cd226* with anti-CTLA-4 containing treatments and neo VAX reducing *Tigit* expression within clusters expressing the highest levels of *Tigit* (Cd8**_Eff/Ex_**, Cd8**_Cycling_**, Cd8**_Ccr7_**) **(Figures S5, S6A, S6B, S6C, and S6F)**.

### Anti-PD-1 expands PD-1^+^ TCF7^−^ NeoAg-specific Teff/Tex and robustly expands Bhlhe40^hi^ PD-1^+^ TCF7^−^ NeoAg-specific Teff/Tex when combined with anti-CTLA-4

We and others previously demonstrated that tumor antigen-specific CD8 T cells have unique features as compared to bystander CD8 T cells and that immunotherapy primarily affects tumor antigen-specific versus bulk CD8 T cells^13,18,53–55^. Therefore, we monitored CD8 T cells specific for the mLama4 NeoAg in the setting of neo VAX or ICT **(Figure 3A)**. Anti-CTLA-4 and/or anti-PD-1 increased the overall frequency of intratumoral CD8 T cells with anti-CTLA-4 (+/- anti-PD-1) also driving a significant increase in mLama4-specific CD8 T cells as a percent of CD8 T cells or CD45^+^ cells and anti-PD-1 significantly increasing mLama4-specific CD8 T cells as a percent of CD45^+^ cells **(Figures 3B-3D and S7A)**. Notably, neo VAX drove the greatest increase in mLama4-specific CD8 T cells from less than 2% (control mAb or control VAX) to over 20% of CD8 T cells, which accounted for over 4% of intratumoral CD45^+^ cells **(Figures 3C, 3D, and S7A)**.

**Figure 3.**
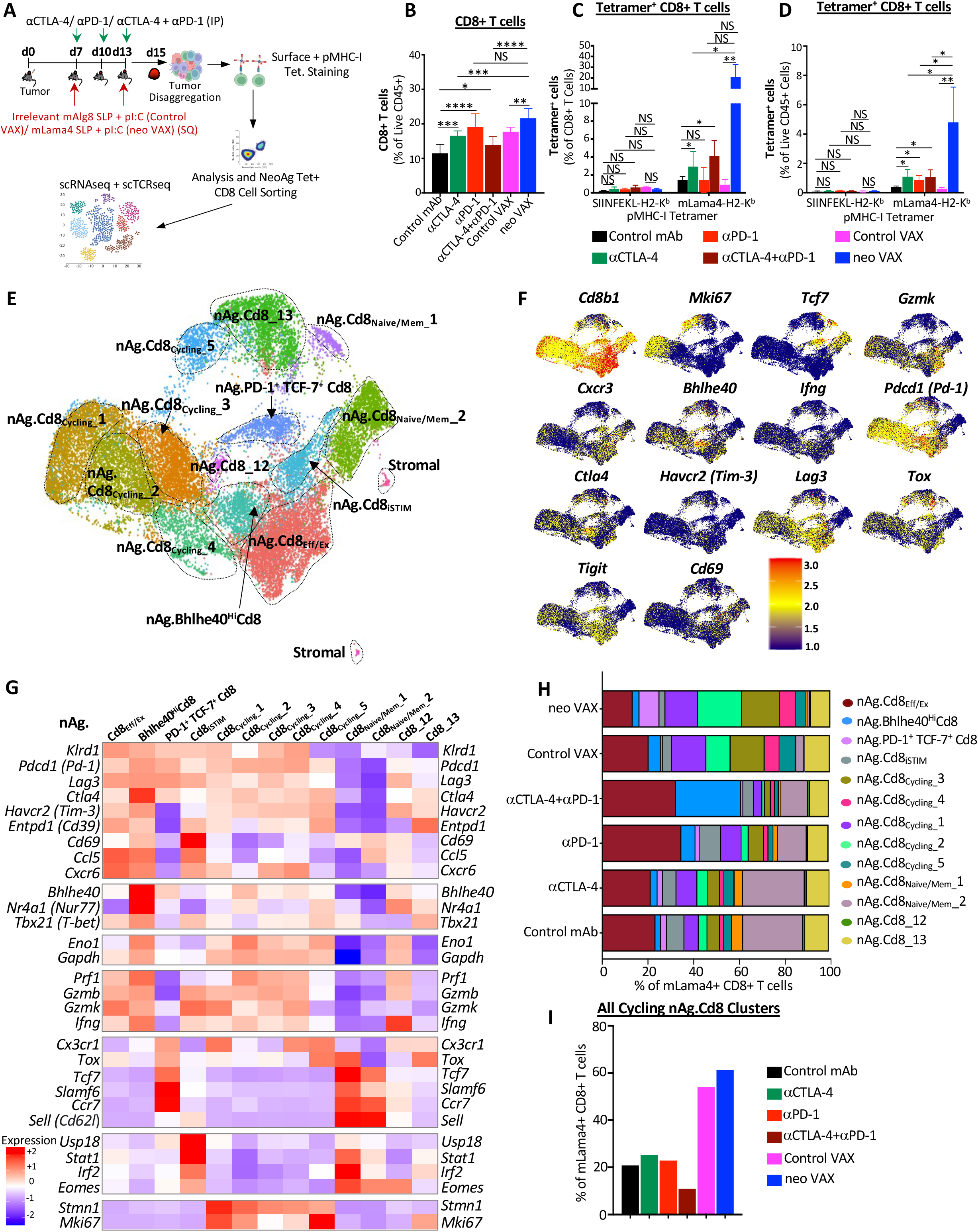
NeoAg vaccines and ICT induce shared and distinct alterations to NeoAg-specific CD8 T cells. **(A)** Experimental setup for **(B)-(I)**. WT C57BL/6J mice were injected with Y1.7LI melanoma cells and subsequently treated beginning on d. 7 with control mAb, anti-CTLA-4, anti-PD-1, anti-CTLA-4 + anti-PD-1, irrelevant (for Y1.7LI) mAlg8 SLP + pI:C (control VAX), or relevant mLama4 SLP + pI:C (neo VAX) and harvested on d. 15 post-tumor transplant. Single cell suspensions of harvested tumors were stained with SIINFEKL- or mLama4-H2-K^b^ PE and APC labelled tetramers and surface stained with flow antibodies for analysis or sorting of mLama4 tetramer positive CD8 T cells for scRNAseq. **(B)** Graph displaying CD8 T cells as a percentage of intratumoral live CD45^+^ cells in Y1.7LI tumors under different treatment conditions. **(C)** and **(D)** Graph displaying mLama4 tetramer-positive CD8 T cells as a percentage of **(C)** CD8 T cells and **(D)** CD45^+^ cells in Y1.7LI tumors under different treatment conditions. **(E)** UMAP plot from scRNAseq of mLama4 NeoAg-specific CD8 T cells. Cell types were annotated based on transcriptional states of NeoAg-specific CD8 T cells. **(F)** Feature plots displaying expression of select phenotype and lineage transcripts. **(G)** Heat map displaying average expression of select transcripts by cluster. **(H)** Bar graph displaying frequency of mLama4 NeoAg-specific CD8 T cells within each cluster by treatment condition. **(I)** Frequency of total mLama4 NeoAg-specific CD8 T cells within the 5 cycling clusters combined by treatment condition. See also **Figures S7** and **S8**.

Since our scRNAseq profiling of CD45^+^ cells did not distinguish NeoAg-specific CD8 T cells, we profiled NeoAg-specific CD8 T cells by sorting intratumoral mLama4 tetramer positive CD8 T cells from mice under different treatment conditions **(Figure 3A)**. We profiled between 937 to 1762 mLama4-specific CD8 T cells for each of the different ICT treatment conditions and 4459, 6723, and 7646 mLama4-specific CD8 T cells for control mAb, control VAX, and neo VAX, respectively. The two smallest clusters contained contaminating stromal cells, with the remaining clusters comprising NeoAg-specific CD8 T cells that we annotated based on expression of select transcripts and gene set enrichment patterns **(Figures 3E-3G, S7B, S7C, S8, and S9)**; this enabled us to distinguish additional features that were not evident from profiling bulk CD8 T cells.

Clusters nAg.Cd8**_Eff/Ex_** and nAg.**Bhlhe40^Hi^**Cd8 expressed *Pdcd1, Havcr2* (TIM-3), *Lag3,* and *Tigit,* as well as effector transcripts (e.g., *Nkg7*, *Ccl5, Gzmb*, *Gzmk*, *Prf1, Cxcr6*). These two clusters also expressed *Tox* and exhibited little to no detectable expression of *Tcf7* **(Figures 3F, 3G, S7B, and S7C)**, consistent with effector and/or dysfunctional/exhausted CD8 T cells. neo VAX most notably reduced the proportion of nAg.Cd8**_Eff/Ex_** cells, whereas the proportion of cells in this cluster increased with anti-PD-1 (+/- anti-CTLA-4) **(Figure 3H)**. In nAg.**Bhlhe40^Hi^**Cd8, the top defining marker of this cluster was *Bhlhe40* **(Figures 3G and S8)**, which we previously demonstrated was upregulated in tumor-specific T cells and required for CD4 and/or CD8 T cell effector function and response to ICT^21^. In addition to *Bhlhe40* (as well as *Pdcd1, Havcr2,* and *Lag3*), this cluster also expressed other transcripts induced by TCR activation, including *Ctla4, Cd69, Nr4a1* (Nur77), and *Nr4a3* and also displayed high expression of *Tbx21* (T-bet) and *Ifng* **(Figures 3G and S7B)**. As compared to control mAb treatment where nAg.**Bhlhe40^Hi^**Cd8 represented ∼2.4% of mLama4-specific CD8 T cells, a small increase in frequency was observed with anti-CTLA-4, control VAX, or neo VAX, and a more substantial ∼2.6-fold increase occurred with anti-PD-1 **(Figure 3H)**. Strikingly, anti-CTLA-4 and anti-PD-1 combination ICT increased this cluster to over 28% of mLama4-specific CD8 T cells.

In addition to increasing the frequency of cells within PD-1^+^ TCF7^−^ Teff/Tex clusters (nAg.Cd8**_Eff/Ex_** and nAg.**Bhlhe40^Hi^**Cd8), combination ICT increased expression of *Bhlhe40, Fasl, Il7r, Icos,* and *Cd28*, while decreasing *Tox*, *Pdcd1, Lag3, Entpd1,* and *Tigit* expression within both clusters **(Figures S7B and S7C)**. Further, combination ICT decreased *Havcr2* and increased *Cd69* expression in cluster nAg.**Bhlhe40^Hi^**Cd8. The decrease in *Tox*, *Pdcd1, Lag3, Entpd1,* and *Tigit* (and *Havcr2* in nAg.**Bhlhe40^Hi^**Cd8) was also observed with anti-CTLA-4 (but not with anti-PD-1) **(Figures S7B and S7C)**, suggesting that these changes induced by combination therapy were primarily driven by anti-CTLA-4. In contrast, increased *Bhlhe40* expression was most prominent in the presence of anti-PD-1. Other features (e.g., increased *Icos*, *Cd28*, and *Fasl*) were unique to anti-CTLA-4 and anti-PD-1 combination ICT treatment **(Figure S7B)**.

### NeoAg vaccination preferentially increases PD-1^+^ TCF7^+^ stem-like as well as proliferating NeoAg-specific CD8 T cells

Amongst the most prominent NeoAg vaccine-driven changes, NeoAg vaccines drove an over 3-fold increase in the frequency of mLama4-specific CD8 T cells within cluster nAg.**PD-1^+^TCF7^+^**Cd8 as compared to control mAb and over 8-fold increase as compared to control VAX **(Figure 3H)**. Cluster nAg.**PD-1^+^TCF7^+^**Cd8 displayed high expression of *Pdcd1*; low to moderate expression of *Ifng*, *Gzmk*, and *Prf1*; and little to no detectable expression of *Havcr2* or *Entpd1* **(Figures 3G and S7B)**. nAg.**PD-1^+^TCF7^+^**Cd8 also expressed transcripts encoding molecules related to T cell homing such as *Ccr7*, as well as *Bach2*^56^, *Slamf6*, and *Tcf7*, consistent with CD8 T cells with plastic or stem-like properties seen in progenitor exhausted CD8 T cells **(Figures 3G, S7B and S8)**. While NeoAg vaccines promoted this population, the proportion of NeoAg-specific CD8 T cells within this cluster was largely unchanged with anti-CTLA-4, reduced slightly with anti-PD-1, and even further reduced with combination anti-CTLA-4 and anti-PD-1 **(Figure 3H)**. Anti-CTLA-4 containing treatments displayed decreased expression of *Pdcd1*, *Lag3*, *Tigit* and increased expression of transcripts encoding molecules related to T cell quiescence and homing such as *S1pr1, Sell* (Cd62l), and *Klf2*, as well as *Il7r* **(Figures S7B and S7C)**.

We annotated 5 clusters of “cycling” NeoAg-specific CD8 T cells displaying a range of activation states and proliferation signatures **(Figures S7B and S8)**. NeoAg vaccination and control VAX increased the frequency of cells each of the 5 cycling NeoAg-specific CD8 T cell clusters, although to differing degrees **(Figure 3I)**. This suggests that although far more NeoAg-specific CD8 T cells are observed within tumors treated with neo VAX as compared to control VAX **(Figures 3C and 3D)**, within NeoAg-specific CD8 T cells, both control VAX and neo VAX promotes cycling tumor-specific CD8 T cells. Together, these 5 cycling clusters represented 20.9% of all mLama4-specific CD8 T cells under control mAb treatment, 54.1% under control VAX treatment, and 61.3% under neo VAX treatment **(Figure 3I)**. The frequency of total cells within cycling clusters was modestly increased by anti-CTLA-4 or anti-PD-1 ICT, whereas anti-CTLA-4 plus anti-PD-1 combination ICT decreased the frequency by almost half. Within nAg.Cd8**_Cycling__1**, nAg.Cd8**_Cycling__3**, and nAg.Cd8**_Cycling__4**, either control VAX or neo VAX increased the frequency of NeoAg-specific CD8 T cells to about the same level **(Figure 3H)**. In contrast, nAg.Cd8**_Cycling__2** represented 10.6% of NeoAg-specific CD8 T cells under control VAX conditions, whereas under neo VAX conditions, the frequency of cells within this cluster increased to 19.2% of NeoAg-specific CD8 T cells **(Figure 3H)**. As compared to the other cycling clusters, nAg.Cd8**_Cycling__2** expressed higher *Xcl1*, *Tnfrsf4* (OX40), *Tnfrsf9* (4-1BB), *Prf1*, and *Ifng* **(Figures S7B and S9)**.

### TCR repertoire clonality is associated with different NeoAg-specific CD8 T cell states

We next assessed the relationship between TCR clonality and phenotype of mLama4 NeoAg-specific CD8 T cells. A total of 15,668 clonotypes expressing both TCR alpha and beta chains (**Figures 4A-4C)** and 17,492 NeoAg-specific CD8 T cells with at least one productive TCR alpha or beta chain or both **(Figures S10A-S10C)** were analyzed separately and primarily focused our analyses on clonotypes expression both TCR alpha and beta. Amongst NeoAg-specific CD8 T cells with both TCR alpha and beta with an activated phenotype, the 5 cycling NeoAg-specific CD8 T cell clusters display highest overlapping TCR clonotypes with each other and nAg.**Bhlhe40^Hi^**Cd8 **(Figures 4A and 4B)**. nAg.Cd8**_Eff/Ex_** also displayed overlap with nAg.**Bhlhe40^Hi^**Cd8 and cycling CD8 T cell clusters. Although nAg.**PD-1^+^TCF7^+^**Cd8 contained far fewer overlapping TCR clonotypes, nAg.**PD-1^+^TCF7^+^**Cd8 with TCR expressing both alpha and beta chain, shared the largest frequency of clonotypes with nAg.Cd8_**13**, followed by nAg.Cd8**_iSTIM_**, nAg.Cd8_**12** and nAg.Cd8**_Eff/Ex_ (Figure 4B)**.

**Figure 4.**
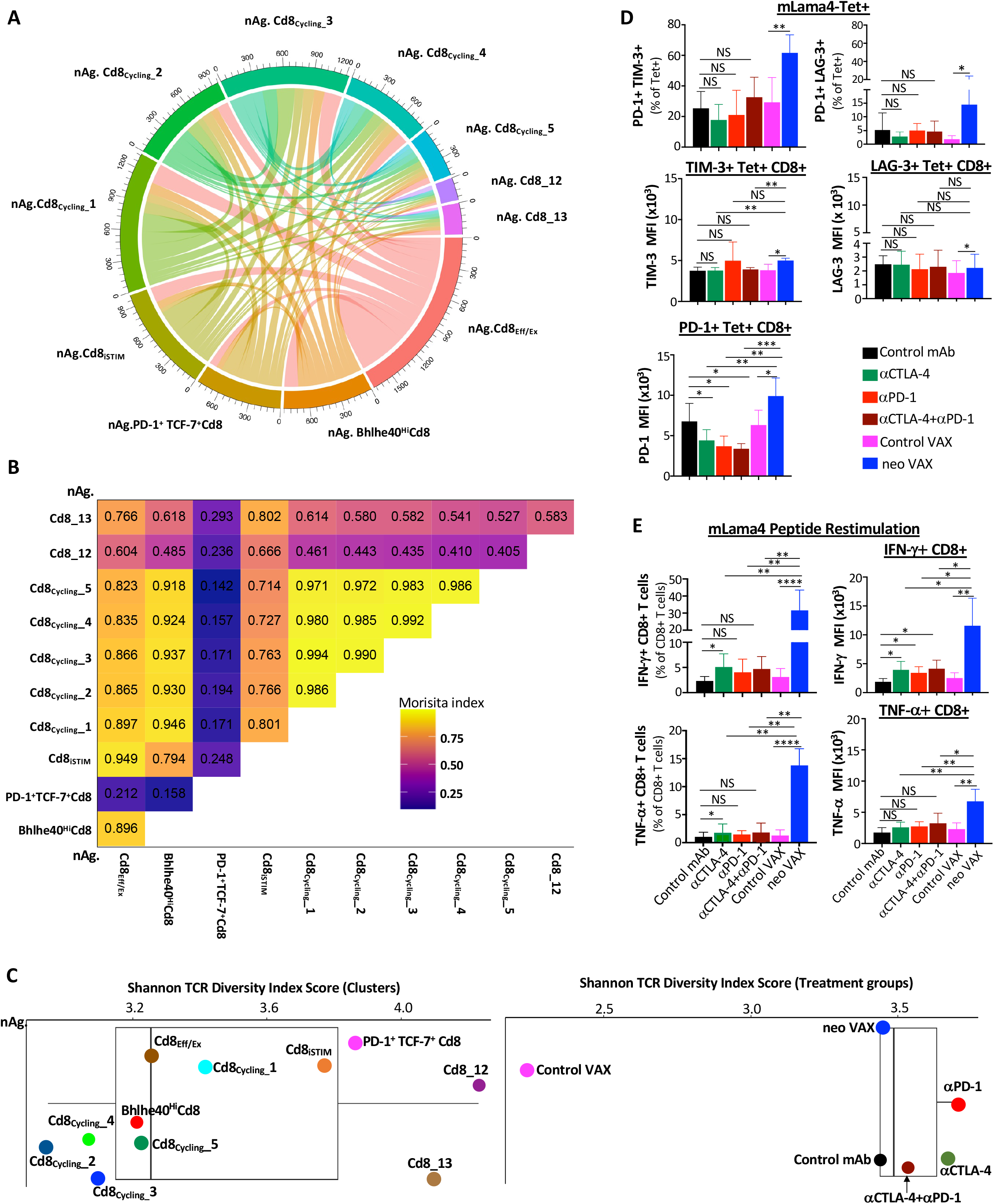
NeoAg-specific alpha-beta TCR clonotype expansion and diversity relates to phenotype and functional state of T cells associated with different immunotherapies. **(A)** Chord diagram displaying overlapping TCR clonotypes of mLama4 NeoAg-specific CD8 T cells by cluster. **(B)** Morisita index values depicting overlapping TCR clonotypes of mLama4 NeoAg-specific CD8 T cells by cluster. **(C)** Shannon TCR diversity index by clusters and treatment groups. **(D)** Graphs displaying percent of PD-1^+^ TIM-3^+^/LAG-3^+^ or MFI of PD-1, TIM-3, or LAG-3 on PD-1^+^, TIM-3^+^, or LAG-3^+^ mLama4-specific CD8 T cells in Y1.7LI tumors under different treatment conditions and harvested on d. 15 post-tumor transplant. **(E)** Graph displaying IFN-γ^+^ or TNF-α^+^ CD8 T cells and IFN-γ or TNF-α MFI as assessed by intracellular cytokine staining of mLama4 peptide restimulated CD8 T cells isolated from Y1.7LI tumors under different treatment conditions and harvested on d. 15 post-tumor transplant. Bar graphs in **(D)** and **(E)** display mean ± SEM and are representative of at least three independent experiments (\**P* < 0.05, \*\**P* < 0.01, \*\*\**P* < 0.005, **** *P* < 0.0001; NS, not significant, unpaired t test). See also **Figure S10.**

Shannon Diversity Index suggested a lower TCR diversity in the cycling clusters, nAg.**Bhlhe40^Hi^**Cd8, and nAg.Cd8**_Eff/Ex_** with nAg.Cd8**_iSTIM_** displaying greater diversity **(Figure 4C)**. nAg.**PD-1^+^TCF7^+^**Cd8 and nAg.Cd8_13 displayed greater diversity, with nAg.Cd8_12 displaying the highest Shannon Diversity Index score. Comparing treatment groups, the largest putative increase in NeoAg-specific CD8 T cell TCR diversity with TCR alpha and beta pair occurred with either anti-PD-1 or anti-CTLA-4 ICT followed by anti-CTLA-4 and anti-PD-1 combination ICT, with all ICT treatment groups displaying a higher Shannon Diversity Index score than control mAb and neo VAX, which had a similar diversity score **(Figure 4C)**. Amongst NeoAg-specific CD8 T cells with one or both TCR alpha or beta chain, anti-CTLA-4 exhibited the highest Shannon TCR Diversity Score, followed by anti-PD-1, anti-CTLA-4 and anti-PD-1 combination ICT, control mAb, and neo VAX **(Figure S10C).** Control VAX displayed by far the least TCR diversity or highest clonality of NeoAg-specific CD8 T cells among all treatment conditions (**Figures 4C and S10C)**.

Thus, while ICT increase TCR diversity amongst mLama4 NeoAg-specific CD8 T cells, NeoAg vaccines induce mLama4 NeoAg-specific CD8 T cells with more expanded clonotypes and less diversity compared to ICT.

### NeoAg vaccines induce robust expansion of NeoAg-specific IFN-γ^+^ CD8 T cells expressing PD-1 and LAG-3 and/or TIM-3

Since we noted that mice treated with neo VAX displayed a greater frequency of PD-1^+^ TIM-3^+^ NeoAg-specific CD8 T cells as compared to other conditions when treatment was initiated on d. 3 post-tumor transplant **(Figure S1F)**, we assessed surface expression of PD-1, TIM-3, and LAG-3 on intratumoral mLama4 NeoAg-specific CD8 T cells from mice when treatment initiation occurred on d. 7 (as in our scRNAseq experiments). As expected, a majority of NeoAg-specific CD8 T cells expressed PD-1, with similar frequencies of PD-1^+^ TIM-3^+^ or PD-1^+^ LAG-3^+^ NeoAg-specific CD8 T cells observed between control mAb, control VAX, and the different ICT treatment conditions **(Figures 4D and S7D)**. However, expression of PD-1 on a per cell basis was lower in ICT treated groups. In contrast, a dramatic increase in the percentage of PD-1^+^ TIM-3^+^ or PD-1^+^ LAG-3^+^ NeoAg-specific CD8 T cells was observed in mice treated with neo VAX and amongst PD-1^+^, TIM-3^+^, or LAG-3^+^ NeoAg-specific CD8 T cells, PD-1, TIM-3, and LAG-3, respectively, was expressed higher in the neo VAX treated group (**Figure 4D**). Intracellular cytokine staining (ICS) on isolated intratumoral CD8 T cells restimulated with the mLama4 NeoAg peptide revealed that anti-CTLA-4 increased the frequency of IFN-γ^+^ or TNFα^+^ CD8 T cells, while neo VAX induced the greatest expansion (> 5-fold) of IFN-γ^+^ or TNFα^+^ CD8 T cells **(Figure 4E)**. Amongst mLama4 NeoAg-stimulated IFN-γ^+^ CD8 T cells, expression of IFN-γ increased significantly with anti-CTLA-4 and/or anti-PD-1, with neo VAX prompting the most robust increase **(Figure 4E)**.

### Anti-CTLA-4 promotes Th1-like CD4 T cells expressing ICOS and Bhlhe40, while combination anti-CTLA-4 and anti-PD-1 ICT induces a small subset of Th2-like CD4 T cells

Since effective neo VAX or anti-CTLA-4/anti-PD-1 ICT require not only CD8 T cells, but also CD4 T cells **(Figure S2A)**, we examined CD4 T cells from our scRNAseq performed on sorted CD45^+^ cells **(Figure 2A)**. Anti-CTLA-4 prominently induced a higher frequency of conventional CD4 T cells and reduced the percentage of Tregs as assessed by both scRNAseq and flow cytometry (**Figures 2G-2I, 2K, S3A, and S3B)**. Notably, anti-CTLA-4 (+/- anti-PD-1) induced subpopulations of Th1-like cells expressing *Ifng* and *Bhlhe40*, including cluster **ICOS^hi^Bhlhe40^hi^** CD4**_Th1_** that also highly expressed *Icos*, *Pdcd1*, *Ctla4*, *Cxcr6*, *Csf2* (GM-CSF), *Fasl*, *Furin* (encoding a TCR/IL-12-STAT4-induced proprotein convertase), and *Tnfaip3* (encoding the A20 protein that regulates TCR/CD28-mediated NF-κB activation and TCR-mediated survival) **(Figures 2E, 5A, 5B, S5 and S11A). ICOS^hi^Bhlhe40^hi^** CD4**_Th1_** displayed enrichment in IL-2 STAT5 and IL-6 JAK STAT3 signaling, TNFa signaling via NF-κB, and IFN-γ response gene sets amongst others **(Figure S11A)**. neo VAX also exhibited a greater frequency of cells within this cluster as compared to control VAX **(Figure 5B)**. Cd4**_Th1__A** also expressed *Icos* and *Bhlhe40*, but to less of an extent than **ICOS^hi^Bhlhe40^hi^** CD4**_Th1_** (**Figures 5A and S5**). Cd4**_Th1_**_**A** was further distinguished from **ICOS^hi^Bhlhe40^hi^** CD4**_Th1_** by lower *Furin, Cxcr6, Runx3, Tnfaip3, Pdcd1, Havcr2*, and *Lag3* expression and higher *Tbx21* (Tbet) and *Il7r* expression. Although both clusters expressed glycolytic enzyme transcripts, greater expression of several of these transcripts was seen in **ICOS^hi^Bhlhe40^hi^** CD4**_Th1_**, while Cd4**_Th1__A** displayed gene set enrichment in Fatty Acid Metabolism **(Figures S5, S11A, and S11B)**. Additionally, both clusters displayed significant enrichment in TGF beta signaling gene sets **(Figures S11A and S11B)**. Anti-CTLA-4 dramatically increased the frequency of Bhlhe40^+^ CD4**_Th1__A**, with anti-PD-1, and to less of an extent neo VAX, also increasing cells within this cluster **(Figure 5B)**. CD4**_Th1_**_**B** was the smallest cluster of Th1-like cells and exhibited high *Ifng*, *Pdcd1*, *Havcr2, and Tigit* expression **(Figures 5A, S5, and S11C)**. This cluster also expressed the highest level of *Lag3* and *Tox* amongst all CD4 clusters **(Figures 2E, 5A and S5)**. Only subtle changes to the frequency of cells within this cluster were seen with treatments apart from control VAX and combination anti-CTLA-4 and anti-PD-1, with the latter displaying the highest frequency of cells within this cluster amongst all conditions **(Figure 5B).**

**Figure 5.**
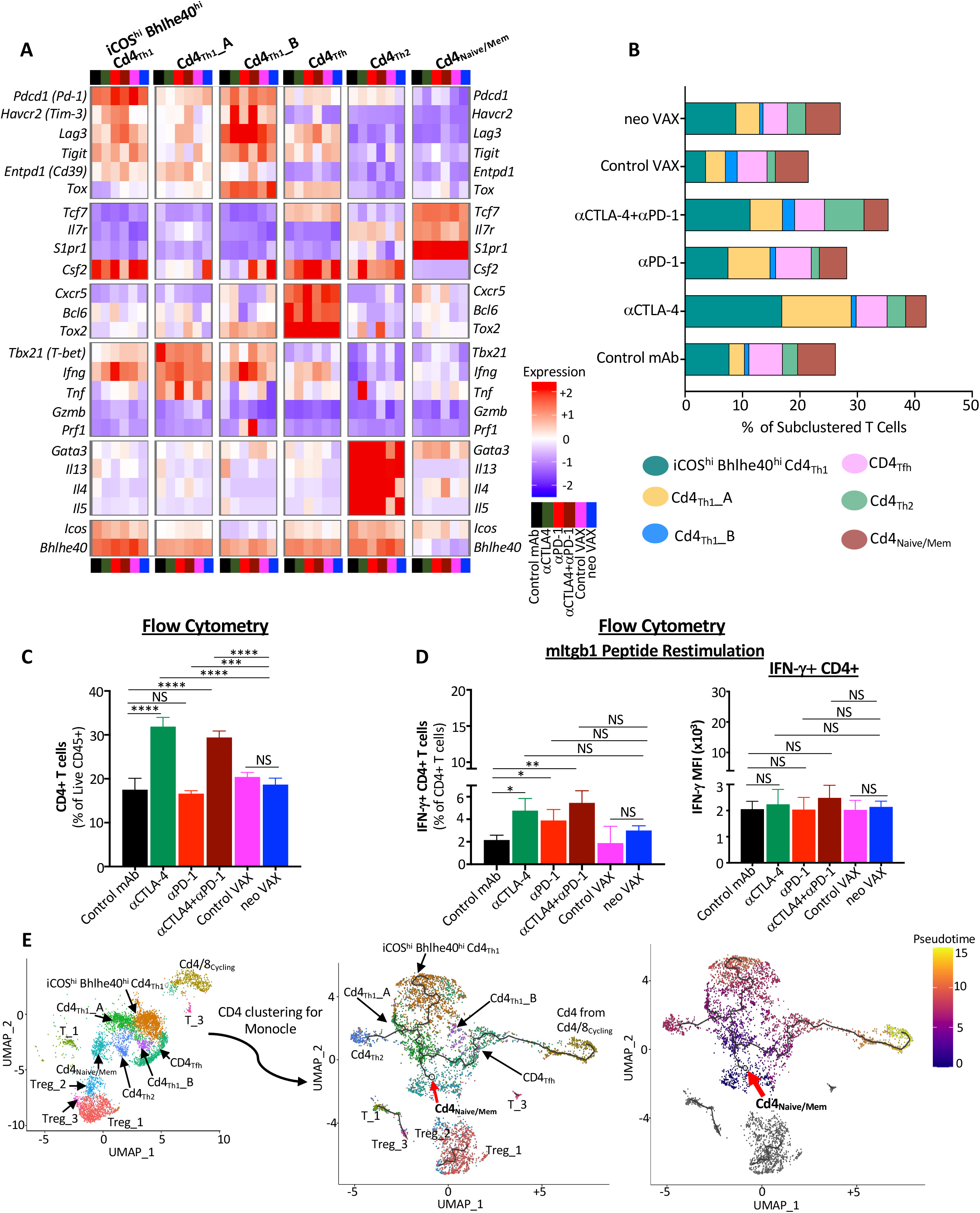
Anti-CTLA-4 induces an ICOS^+^ Bhlhe40^+^ Th1-Like subpopulation of CD4 T cells and a small Th2-Like subpopulation when combined with anti-PD-1. **(A)** Heat map displaying normalized expression of select genes in each CD4 T cell cluster by treatment condition. **(B)** Bar graphs depicting frequency of CD4 T cells within each cluster by treatment condition. **(C)** Graph displaying CD4 T cells as a percentage of intratumoral live CD45^+^ cells as determined by flow cytometry in Y1.7LI tumors under different treatment conditions and harvested on d. 15 post-tumor transplant. **(D)** Graph displaying IFNγ^+^ CD4 T cells as assessed by intracellular cytokine staining on CD4 T cells isolated from Y1.7LI tumors under different treatment conditions and harvested on d. 15 post-tumor transplant. **(E)** Monocle 3-Guided Cell Trajectory of CD4 T Cell Clusters. UMAP plot displaying exclusively CD4 T cell-containing clusters (left) of all experimental conditions, CD4 T cell trajectory graph overlaid on UMAP (middle) where the origin of the inferred pseudotime is indicated by the red arrow and assigned with pseudotime score 0, and geodesic distances and pseudotime score among other CD4 T cells are calculated from there based on transcripts associated cell states. CD4 T cell clusters overlaid on Monocle3 pseudotime plot (right). Bar graphs in **(C)** and **(D)** display mean ± SEM and are representative of at least three independent experiments (\**P* < 0.05, \*\**P* < 0.01, \*\*\**P* < 0.005, \*\*\*\**P* < 0.0001, NS, not significant, unpaired t test). See also **Figure S11**.

The increase in IFN-γ expressing Th1-like cells most prominently induced by anti-CTLA-4 was reflected by ICS on isolated intratumoral CD4 T cells restimulated *ex vivo* with the mItgb1 MHC-II NeoAg peptide. Anti-CTLA-4 +/- anti-PD-1 induced the strongest increase in the overall frequency of conventional CD4 T cells, with anti-CTLA-4 and/or anti-PD-1 increasing the frequency of IFN-γ^+^ CD4 T cells upon restimulation with mItgb1 peptide **(Figures 5C and 5D)**. This is in contrast to neo VAX, where only subtle changes were observed. Altogether, these findings indicate that while mice treated with anti-CTLA-4, alone or in combination with anti-PD-1, display the most dramatic increase in IFN-γ-producing Th1-like CD4 T cells within the tumor, anti-PD-1 also incites IFN-γ^+^ CD4 T cells **(Figure 5D)**. This is also supported by comparing the expression of *Ifng* transcript within *Ifng*^+^ CD4 T cell clusters, where anti-PD-1 induced increased *Ifng* expression, even in clusters whose frequency was unaltered by anti-PD-1 (**Figures 5A, S5, and S11A-S11D**).

Interestingly, combination ICT induced expansion of Cd4**_Th2_**, a small cluster that express *Icos* and *Bhlhe40*, as well as *Furin, Tnfaip3*, *Cd28*, and *Il7r*. Unlike the other ICOS^+^ Bhlhe40^+^ clusters, *Ifng, Havcr2,* and *Lag3* were barely detectable and instead, Cd4**_Th2_** expressed *Gata3*, *Il4*, *Il5*, and *Il13,* indicative of Th2-like CD4 T cells (**Figures 5A, S5, S11E, and S11F**).

To gain insight into the temporal dynamics of the observed changes in CD4 T cells, we used Monocle to analyze scRNAseq data^57^. Monocle suggested that the starting point for conventional CD4 T cells corresponds to cells within either the Cd4**_Naive/Mem_** cluster (expressing *Tcf7, Il7r,* and *S1pr1*) or CD4 T cells within the Cd4/8**_Cycling_** cluster **(Figure 5E)** with Cd4**_Tfh_** (displaying T follicular helper-like transcriptional features) connecting Cd4/8**_Cycling_** CD4 T cells to the main trajectory towards Cd4**_Naive/Mem_** and the branch to more activated, polarized CD4 T cells. Notably, a pseudotime trajectory branch point occurs whereby activated CD4 T cells occupy Th1-like **ICOS^hi^Bhlhe40^hi^**Cd4**_Th1_** driven by anti-CTLA-4 (+/- anti-PD-1) (and to a lesser extent by neo VAX) or encounter another branch whereby they assume one of two fates: they either become Th1-like CD4 T cells within Cd4**_Th1__A** or become Th2-like Cd4**_Th2_**, with Cd4**_Th1__A** being induced by anti-CTLA-4 and/or anti-PD-1 or neo VAX and Cd4**_Th2_** primarily being driven by combination anti-CTLA-4 and anti-PD-1 **(Figure 5E)**.

### Features of intratumoral Treg subpopulations during NeoAg vaccine or ICT treatment

We also identified three CD4 Foxp3^+^ Treg clusters **(Figures S3B)**. Treg_1 and Treg_3 appeared to be the most activated with Treg_3 expressing the highest level of *Ctla4*, *Havcr2*, and *Klrg1* **(Figure S5)**. Mice treated with anti-CTLA-4 alone or in combination with anti-PD-1 experienced a decrease in frequency of Treg_1 and Treg_3 **(Figures S3B)**, which is consistent with previous results that the anti-CTLA-4 mAb we used (mouse IgG2b; clone 9D9) partially depletes Tregs, especially those highly expressing CTLA-4^19,21–23,58–60^. Treg_2 expressed lower amounts of *Ctla4, Havcr2, Tigit*, and *Klrg1* with the frequency of these Tregs not being affected by anti-CTLA-4, whereas anti-PD-1 with or without anti-CTLA-4, control VAX, or neo VAX displaying a greater frequency of cells in this cluster **(Figure S3B)**. As compared to control VAX, the cellular density of Treg_1 and Treg_2 decreased in tumors from mice treated with neo VAX **(Figures S3B)**. Further, transcript expression of *Foxp3* in Treg_2 was lower in the neo VAX group. These alterations to the overall frequency of Tregs most prominently observed in the presence of anti-CTLA-4 were also corroborated by flow cytometry analysis **(Figure S3A)**.

### Intratumoral myeloid cell compartment during NeoAg vaccines or ICT treatment

To characterize intratumoral monocytes/macrophages and DCs, we subclustered myeloid cells excluding the single cluster of neutrophils **(Figures 2B, 2C, S3A, and S12A)**. In addition to a cluster of plasmacytoid DCs (pDCs), four other DC clusters were identified **(Figures S12A-S12E)**. Cluster CD103^+^cDC1 expressed multiple classical DC (cDC) 1 transcripts including *Itgae* (*Cd103*), *Xcr1*, and *Clec9a* **(Figures S12B and S12E)**. CD63^+^Ccr7^+^cDC and Ccr7^+^cDC expressed *Ccr7*, *Cd1d1*, *Cd200*, *Fscn1*, *Cd274* (PD-L1), and *Pdcd1lg2* (PD-L2). As compared to Ccr7^+^cDC, CD63^+^Ccr7^+^cDC expressed higher *Cd63, Cd40, Btla,* and *Cd70* **(Figures S12D and S12E)**. These two migratory cDC clusters are consistent with mregDCs, a term describing a maturation state of cDC1s and cDC2s upon uptake of tumor antigen and although they express immunoregulatory molecules, they are not necessarily immunosuppressive^61,62^.

### Distinct Macrophage Remodeling Induced by NeoAg Vaccines and ICT

Overall, monocytes/macrophages represented a plurality of intratumoral CD45^+^ cells and displayed a range of phenotypic states^63,64^ (**Figures 6A, S3A, and S13**). Ccr2^+^M_**c1** displayed transcripts consistent with monocytes, including *Ccr2* and *Chil3*, and the frequency of cells within this cluster increased slightly with anti-PD-1 or neo VAX **(Figures 6A, 6B, and S13C)**. While *Chil3^+^* monocytes were previously shown to be reduced by a NeoAg vaccine in preclinical models^65^, the NeoAg vaccine and adjuvant used in that setting differed from ours.

**Figure 6.**
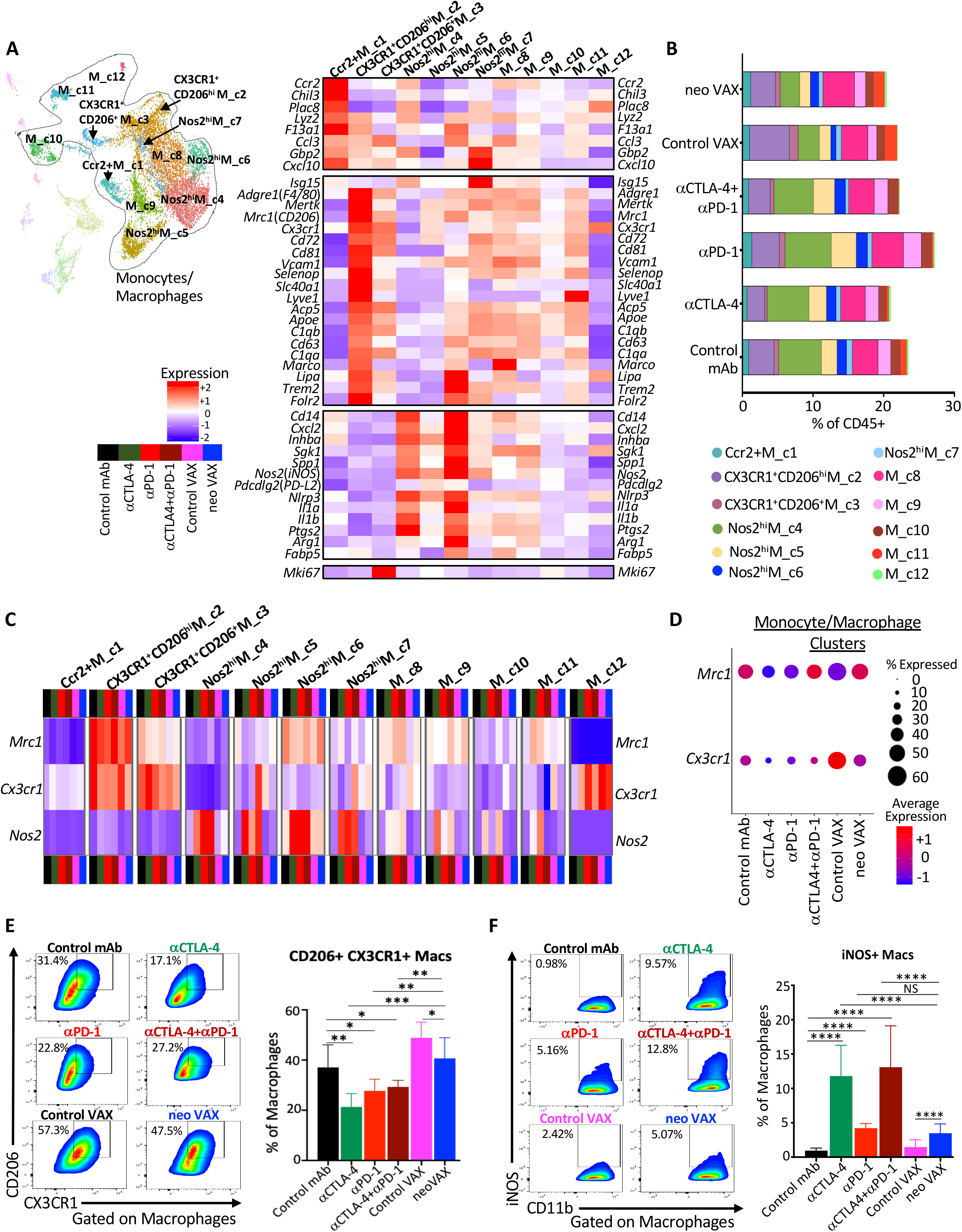
NeoAg vaccines promote partially distinct macrophage remodeling from ICT. **(A)** UMAP displaying sub-clustering of select myeloid clusters from CD45^+^ scRNAseq analysis (See Figure 2A) and heat map displaying normalized expression of select genes in each monocyte/macrophage cluster. **(B)** Percent monocytes/macrophages in each cluster by condition and treatment represented as percent of live CD45^+^ cells. **(C)** Heat map displaying normalized expression of *Mrc1* (CD206), *Cx3cr1*, and *Nos2* (iNOS) in each monocyte/macrophage cluster by treatment condition. **(D)** scRNAseq dot plot depicting expression level/percent of cells expressing *Mrc1* and *Cx3cr1* within all monocytes/macrophages clusters by treatment condition. **(E)** Representative flow cytometry plots and graph displaying CX3CR1^+^CD206^+^ macrophages in Y1.7LI tumors under different treatment conditions and harvested on d. 15 post-tumor transplant. **(F)** Representative flow cytometry plots and graph displaying iNOS^+^ macrophages in Y1.7LI tumors under different treatment conditions and harvested on d. 15 post-tumor transplant. For flow cytometry analysis in **(E)** and **(F)**, dot plot displaying CX3CR1^+^CD206^+^ and iNOS^+^ macrophages are gated on macrophages using a gating strategy previously described_97_. Bar graphs in **(E)** and **(F)** display mean ± SEM and are representative of at least three independent experiments (\*\**P* < 0.01, \*\*\*\**P* < 0.0001, NS, not significant, unpaired *t* test). See also **Figures S12** and **S13**.

We previously demonstrated that anti-CTLA-4 and/or anti-PD-1 induces macrophage TME remodeling characterized by a reduction in M2-like macrophages co-expressing the fractalkine receptor (CX3CR1) and CD206 and an increase in M1-like iNOS^+^ macrophages in mouse MCA sarcoma models^19,21^. We noted a similar ICT-induced remodeling trend in the Y1.7LI melanoma model. Whereas a slight decrease in the frequency of CX3CR1^+^ CD206^hi^ M_**c2** cells expressing high levels of *Cx3cr1*, *Mrc1 (Cd206)*, *Trem2*, *Vcam1*, *Cd63, Cd81,* and *Cd72* was observed with anti-CTLA-4 +/- anti-PD-1 ICT, expression of *Cx3cr1* and the frequency of *Cx3cr1*^+^ macrophages within this cluster was decreased under all ICT treatment conditions or with neo VAX **(Figures 6A-6C, S13A, and S13C)**. CX3CR1^+^CD206^+^ M_**c3** also expressed *Cx3cr1*, as well as *Mrc1*, *Trem2*, *Vcam1*, and *Cd72* with the latter transcripts being expressed less than in CX3CR1^+^ CD206^hi^ M_**c2 (Figure 6A)**. CX3CR1^+^CD206^+^ M_**c3** also displayed high expression of *Mki67* and exhibited lower *Mertk* expression as compared to CX3CR1^+^ CD206^hi^ M_**c2**. Anti-CTLA-4 reduced the frequency of CX3CR1^+^CD206^+^ M_**c3 (Figures 6B and S13A)**. Although the aforementioned two clusters expressed the highest levels of *Cx3cr1* and *Mrc1*, M_**c8** macrophages also expressed *Cx3cr1* and *Mrc1* under control mAb conditions with ICT reducing expression of *Cx3cr1* within these clusters **(Figures 6C and S13A)**. Comparable expression levels of *Cx3cr1* was observed in M_**c8** under control VAX and neo VAX conditions, with neo VAX increasing the frequency of cells within this cluster **(Figures 6B, 6C, and S13A)**. Under control VAX conditions, a proportion of cells in cluster M_**c10** expressed *Cx3cr1* and *Mrc1*, and under either control VAX or neo VAX conditions, macrophages within cluster M_**c11** expressed both *Cx3cr1* and *Mrc1*. The frequency of cells within M_**c11** increased in mice treated with either control VAX or neo VAX, with ICT reducing this population **(Figures 6B and S13A)**. Overall, monocytes/macrophages from mice treated with control VAX and neo VAX displayed higher average expression of *Cx3cr1* as compared to ICT groups, with neo VAX also displaying similar expression of *Mrc1* as control mAb **(Figure 6D)**.

Several monocyte/macrophage clusters expressed high levels of *Nos2* (iNOS); other clusters expressed varying levels of *Nos2*, with expression of *Nos2* being highly correlated with ICT treatment, as well as neo VAX to some extent **(Figures 6C and S13B)**. Further, expression of *Cd274* also correlated with expression of *Nos2* within macrophage clusters, in particular under ICT treatment conditions **(Figure S13C)**. While the overall frequency of these iNOS^+^ M1-like clusters only modestly increased with ICT, the frequency of cells within these clusters expressing *Nos2* and/or *Nos2* expression on a per cell basis dramatically increased under all ICT conditions **(Figures 6B, 6C, and S13B)**. Nos2^hi^M_**c4** and Nos2^hi^ M_**c6** both manifested high expression of *Nos2*, *Il1a*, *Il1b*, *Cxcl2*, *Inhba,* and *Nfkb1,* signatures of inflammatory macrophages **(Figures 6A and S13C)**. While Nos2^hi^M_**c4** displayed classic features of M1-like macrophages including low *Mrc1* expression, Nos2^hi^ M_**c6** moderately expressed *Mrc1* and exhibited higher *F13a1, Trem2,* and *Il1a*, along with lower *Il1r2* expression compared to Nos2^hi^M_**c4 (Figures 6A and S13C)**. Nos2^hi^M_**c4** displayed high expression of *Cxcl9* and *Spp1*, with expression of the latter diminished with ICT or neo VAX **(Figure S13C)**. Higher *CXCL9* and lower *SPP1* expression was recently found to be correlated with a macrophage prognostic score in cancer patients^66^. Nos2^hi^M_**c5** highly expressed *Nos2* in the presence of ICT, with ICT also increasing the frequency of macrophages within this cluster (**Figures 6B, 6C, and S13B**). This cluster also expressed moderate levels of *Mki67* and other cell cycle related transcripts, indicative of iNOS^+^ macrophages with proliferative capabilities **(Figure 6A)**. Nos2^hi^ M_**c7** was the smallest iNOS^+^ macrophage cluster and in addition to *Nos2* expression under ICT conditions, Nos2^hi^ M_**c7** highly expressed interferon-stimulated genes (ISGs) **(Figures 6A, S13B and S13C)**.

These same overall patterns were manifested at the protein level where in anti-CTLA-4 and/or anti-PD-1 treated mice, the frequency of intratumoral CX3CR1^+^CD206^+^ macrophages decreased with a concomitant increase in iNOS^+^ macrophages **(Figures 6E and 6F)**. In contrast, while neo VAX treated mice also displayed a greater frequency of iNOS^+^ macrophages, CX3CR1^+^CD206^+^ macrophages were only slightly reduced by neo VAX as compared to control VAX, but were maintained at a similar frequency as seen in control mAb treated mice **(Figures 6E and 6F)**. These results reveal that despite a relatively a similar abundance of CX3CR1^+^CD206^+^ macrophages that were previously associated with progressively growing tumors in untreated or control mAb treated mice^19,21^, neo VAX induces tumor regression equivalent to ICT.

### ICT Broadens Therapeutic Window for Neoantigen Vaccines

We noted changes that were not only shared between treatment conditions, but also distinct depending upon which treatment strategy was employed, which was further illustrated by Principle Component Analysis (PCA) **(Figure S14)**. This, together with our findings that neo VAX induces robust expansion of IFN-γ-producing NeoAg-specific CD8 T cells that highly express PD-1 **(Figures 3C, 3D, 4D, 4E, and S7A)**, prompted us to asked whether neo VAX could synergize with ICT. While neo VAX or ICT led to robust rejection of Y1.7LI when initiated on d. 7 post-transplant, a majority of tumor bearing mice displayed tumor outgrowth when treatment with anti-CTLA-4, anti-PD-1, or neo VAX was initiated on d. 12 post-transplant. We therefore used a d. 12 treatment start timepoint to assess whether combining neo VAX with anti-CTLA-4 or anti-PD-1 improved efficacy **(Figure 7A)**. Mice treated with neo VAX in combination with anti-CTLA-4 or anti-PD-1 displayed enhanced tumor control as compared to control VAX (irrelevant SLP + pI:C) + anti-PD-1 or control VAX + anti-CTLA-4 **(Figure 7A)**. Further, neo VAX used in combination with anti-CTLA-4 or anti-PD-1 provided superior tumor growth inhibition compared to combination anti-CTLA-4 and anti-PD-1 ICT. To extend our findings to a distinct tumor model, we assessed our vaccine protocol and combination treatment using the MC38 tumor model, which has several known endogenous MHC-I tumor NeoAgs^17,67,68^. We previously confirmed in our MC38 line the presence of point mutations that form NeoAgs (mAdpgk, mRpl18, and mDpagt1)^17,67^. We assessed combinatorial treatments in MC38 tumor bearing mice by choosing an injection dose of cells and treatment schedule where monotherapy with anti-CTLA-4, anti-PD-1, or neo VAX alone is largely ineffective **(Figure 7B)**. PBS, control VAX, or neo VAX was administered to MC38 tumor bearing mice on d. 12 and 19 post-transplant with or without anti-CTLA-4 or anti-PD-1 given on d. 12, 15, 18, and 22. Similar to results in the Y1.7LI model, neo VAX in combination with anti-CTLA-4 or anti-PD-1 provided superior protection versus monotherapy **(Figure 7B)**. These findings in two distinct models complement ongoing NeoAg vaccine clinical trials and further support the rationale for combination NeoAg-based therapies.

**Figure 7.**
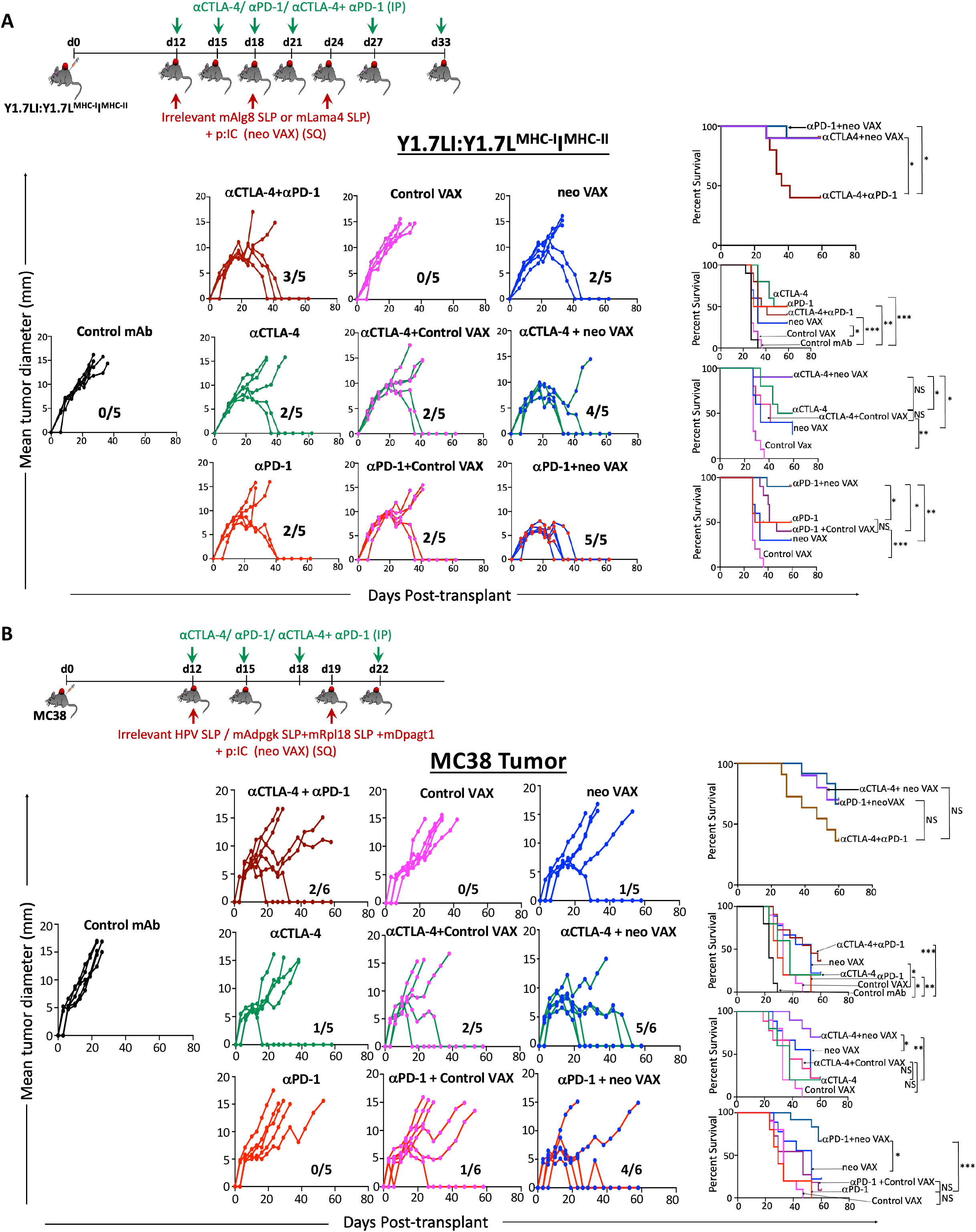
NeoAg vaccines broaden the therapeutic window for anti-CTLA-4 or anti-PD-1 ICT when used in combination. **(A)** Tumor growth and cumulative survival of WT C57BL/6J mice transplanted with Y1.7LI melanoma cells on d. 0 and treated beginning on d. 12 with different monotherapies: control mAb, anti-CTLA-4, anti-PD-1, irrelevant SLP + pI:C (Control VAX), or relevant mLama4 SLP + pI:C (neo VAX); or combination therapies: anti-CTLA-4 + anti-PD-1 combination ICT, anti-CTLA-4 + control VAX, anti-CTLA-4 + neo VAX, anti-PD-1 + control VAX, or anti-PD-1 + neo VAX. **(B)** Tumor growth and cumulative survival of WT C57BL/6J mice transplanted with MC38 cells on d. 0 and treated beginning on d. 12 with different monotherapies: control mAb, anti-CTLA-4, anti-PD-1, irrelevant HPV SLP + pI:C (Control VAX), or relevant mAdpgk SLP + mRpl18 SLP + mDpagt1 SLP + pI:C (neo VAX); or combination therapies: anti-CTLA-4 + anti-PD-1 combination ICT, anti-CTLA-4 + control VAX, anti-CTLA-4 + neo VAX, anti-PD-1 + control VAX, or anti-PD-1 + neo VAX. Tumor growth data in **(A)** and **(B)** are presented as individual mouse tumor growth as mean tumor diameter with fraction indicating (# of mice rejecting tumor)/(# of mice used in experiment) and are representative of three independent experiments. Cumulative survival curves in **(A)** and **(B)** include mice from three independent experiments (\**P* < *0.01*, \*\**P* < *0.05*, \*\*\**P* < *0.001*, log-rank (Mantel–Cox) test).

## Discussion

In this study, we compared different immunotherapies that lead to tumor rejection and pertinent control treatments where tumor progression occurs using mouse melanoma models with relevant gain- and loss-of-function genetic perturbations^45^ and defined NeoAgs. Although prior studies have examined NeoAg vaccines^13,15,17,65,69–71^, few (if any) studies have performed extensive comparisons between NeoAg vaccines, anti-CTLA-4, anti-PD-1, and combination ICT in the same robust experimental system. While most prior studies involving ICT or NeoAg vaccines focused on either lymphoid or myeloid cells^22,69,70,72^, our work has provided insights into both categories of cells and how different immunotherapies differentially affect these cells. Our treatment schedule and analyses were initially performed so that the NeoAg cancer vaccines or ICT we used lead to complete tumor rejection in a majority of mice; thus, we could compare and contrast the molecular and cellular changes that occur as a consequence of NeoAg vaccines or different forms of ICT and link them to outcomes. We specifically chose to study an SLP NeoAg vaccine to complement ongoing clinical trials employing SLPs usually in combination with the adjuvant polyIC:LC^7,10,73^.

The current study makes several key observations. First, NeoAg vaccines and ICT work by several overlapping mechanisms related to the CD8 T cell response, with key differences in the overall magnitude of the response and phenotype of NeoAg-specific CD8 T cells observed. NeoAg vaccines induce the greatest expansion of functional intratumoral NeoAg-specific CD8 T cells including proliferating T cells and PD-1^+^ TCF-1^+^ stem-like CD8 T cells^69,74^. However, anti-CTLA-4 and/or anti-PD-1 also increased the frequency of intratumoral CD8 T cells, including NeoAg-specific CD8 T cells with enhanced production of IFN-γ. Anti-PD-1 alone, or most dramatically when administered in combination with anti-CTLA-4 ICT, induced a subset of Bhlhe40^hi^ NeoAg-specific CD8 T cells also display high expression of *Tbx21* and *Ifng.* We previously documented that ICT promotes Bhlhe40 upregulation in NeoAg tumor-specific T cells and that expression of Bhlhe40 in CD4 and/or CD8 T cells is paramount for effective ICT^21^. A more recent study identified Bhlhe40 as modulating a key differentiation point between progenitor and intermediate subsets of exhausted T cells in an in vitro exhaustion model and chronic LCMV infection^75^. Additionally, Bhlhe40^hi^ NeoAg-specific CD8 T cells expressed *Ctla4, Cd69,* as well as *Nr4a1* and *Nr4a3,* which suggest recent activation and/or TCR stimulation due to their known pattern of rapid and transient expression following T cell stimulation. While some of the alterations in cellular subpopulations and gene/protein expression observed with combination ICT were distinct from either anti-CTLA-4 or anti-PD-1, certain features were also observed with anti-CTLA-4 ICT, whereas other changes were more akin to those observed with anti-PD-1. These findings add to the accumulating evidence that the enhanced anti-tumor activity of combination anti-CTLA-4 and anti-PD-1 ICT is likely mediated by not only additive effects, but also through mechanisms distinct from the monotherapies^19,23^.

Amongst mLama4 NeoAg-specific CD8 T cells with an activated phenotype, cycling/proliferating CD8 T cells displayed a high degree of overlapping TCR clonotypes with each of the cycling clusters, as well as with nAg.**Bhlhe40^Hi^**Cd8. nAg.Cd8**_Eff/Ex_** also displayed overlap with nAg.**Bhlhe40^Hi^**Cd8 and cycling CD8 T cell clusters. A similar observation was made in human non-small cell lung cancers (NSCLC) patients, where the TCRs in CD8 T cells recognizing NeoAgs or non-mutant tumor antigens that expressed markers of exhaustion overlapped to large extent with proliferating CD8 T cells^76^. Shannon Diversity Index suggested a lower TCR diversity in the cycling clusters, nAg.Cd8**_Eff/Ex_**, and nAg.**Bhlhe40^Hi^**Cd8. The lower diversity and high clonotype expansion seen in nAg.Cd8**_Eff/Ex_** and nAg.**Bhlhe40^Hi^**Cd8 are consistent with observations made in human melanoma patients, where it was shown that highly expanded clonotype families were predominantly comprising CD8 T cells expressing markers of exhaustion^53^. Cluster nAg.**PD-1^+^TCF7^+^**Cd8 with a stem-like/progenitor exhausted phenotype displayed greater TCR diversity than cycling clusters, nAg.Cd8**_Eff/Ex_**, nAg.**Bhlhe40^Hi^**Cd8, and nAg.Cd8**_iSTIM_**. We also found that compared to control mAb, a higher Shannon Diversity Index score was observed with any of the ICT treatment conditions assessed, with anti-CTLA-4 promoting the largest putative increase in NeoAg-specific CD8 TCR diversity. NeoAg-specific CD8 TCR from NeoAg vaccine treated mice displayed less diversity, suggesting that NeoAg vaccines promote expansion of NeoAg-specific CD8 T cells with a more restricted TCR repertoire while under control VAX treatment conditions, NeoAg-specific CD8 T cells are highly clonal.

In addition to modulating the CD8 T cell compartment, ICT notably impacted the CD4 T cell compartment as well. Anti-CTLA-4 reduced the frequency of Tregs as expected^19,21,22,58–60^ and induced ICOS^+^ Th1-like conventional CD4 T cells displaying high expression of Bhlhe40^21^. Interestingly, subsets of Th1-like CD4 T cells with high expression of Bhlhe40 were previously found to be enriched in patients with microsatellite instability colorectal cancer, who display favorable outcomes in response to anti-CTLA-4^77^. Further, studies in both preclinical models and human melanoma patients have revealed that anti-CTLA-4 induces ICOS^+^ CD4 T cells expressing IFN-γ^78,79^. Anti-PD-1 also increased the frequency of overall IFN-γ^+^ Th1-like CD4 T cells, but to less of an extent as compared to anti-CTLA-4. Combination anti-CTLA-4 and anti-PD-1 ICT induced a small, but significant subpopulation of Th2-like CD4 T cells (Cd4**_Th2_**).

While vaccines targeting MHC-I NeoAgs predominately altered CD8 T cells, we found that these MHC-I NeoAg vaccines require CD4 T cells for efficacy. The detailed mechanisms regarding the contribution of CD4 T cells in NeoAg vaccines targeting MHC-I NeoAgs remains to be fully elucidated. Although CD4 T cells and MHC-II NeoAgs are critical components of anti-tumor immunity^20,48,80–86^, we specifically chose to utilize an SLP vaccine against a single MHC-I NeoAg to definitively link the MHC-I NeoAg vaccine response to a specific defined NeoAg. Further, since MHC-II NeoAgs are more difficult to predict than MHC-I NeoAgs, we wanted to study the effects of an MHC-I NeoAg vaccine and whether this NeoAg vaccine approach in combination with anti-CTLA-4 or anti-PD-1 ICT could provoke rejection of larger, established tumors. While SLPs offer several advantages over short peptides including the potential to provoke both CD4 and CD8 T cells responses^87,88^; the NeoAg SLPs we used (mAlg8 or mLama4) provoke only NeoAg-specific CD8 T cell responses^13^. Nevertheless, determining whether incorporating an MHC-II NeoAg such as mItgb1 or even a shared, non-mutant antigen will enhance the efficacy of NeoAg vaccines in our models is of future interest.

Beyond the T cell compartment, we noted a divergent impact of NeoAg vaccines on the myeloid compartment compared to ICT. Both ICT and neo VAX increased M1-like iNOS^+^ macrophages to levels higher than with control mAb or control VAX. Since both control VAX and neo VAX contained pI:C, the induction of NeoAg-specific CD8 T cells by the neo VAX, and not just pI:C by itself, likely contributes to some of the changes observed in the macrophage compartment, consistent with observations that peptide vaccine-induced CD8 T cells modify the intratumoral macrophage compartment^89^. ICT reduced the frequency of intratumoral M2-like CX3CR1^+^CD206^+^ macrophages whereas neo VAX (NeoAg SLP + pI:C) treated mice displayed a greater frequency of CX3CR1^+^CD206^+^ macrophages, albeit less than with control VAX (irrelevant SLP + pI:C), as compared to control mAb or ICT treated mice. Therefore, NeoAg vaccines to provoke tumor regression in a TME that is partially distinct from that of ICT. In MCA sarcoma models, we found that ICT-driven induction of iNOS^+^ macrophages was dependent upon IFN-γ, whereas ICT-driven depletion of CX3CR1^+^CD206^+^ macrophages was partially independent of IFN-γ^19^. In our vaccine setting, we hypothesize that favors the induction of T cell-derived IFN-γ and other signals that drives monocyte polarization to iNOS^+^ macrophages upon entering the tumor, but other signals promote maintenance, expansion, or induction of CX3CR1^+^CD206^+^ macrophages as well. These signals are yet unknown but are likely induced by the pI:C (contained in both the control VAX and neo VAX), which acts as a TLR3 agonist in the endosome to potently induce a type I IFN response and can also activate RIG-I/MDA-5 in the cytosol to promote IL-12 production^90,91^. Although we use “M1-like” and “M2-like”, our current study further supports the concept that intratumoral macrophages display a spectrum of activation states and do not fit exclusively into “M1” or “M2” states^63^. While CX3CR1^+^CD206^+^ macrophages display expression patterns consistent with immunosuppressive macrophages, transcriptional profiling and select phenotype marker expression may not distinguish macrophages as immunosuppressive. Nevertheless, it is tempting to speculate that combining NeoAg vaccines that maintain or promote CX3CR1^+^CD206^+^ macrophages expressing high levels of *Trem2* with treatments targeting this macrophage population^42,92^ might enhance the efficacy of NeoAg vaccines.

The unique features induced by each immunotherapy condition prompted us to assess combining NeoAg vaccines with anti-CTLA-4 or anti-PD-1 ICT. In both the Y1.7LI melanoma model and MC38 model, NeoAg vaccines combined with either anti-CTLA-4 or anti-PD-1 leads to equal or even better anti-tumor immune responses than even combination anti-CTLA-4 and anti-PD-1. While up to 20-30% of patients treated with anti-CTLA-4 or anti-PD-1 may experience durable cancer control, ∼50% of metastatic melanoma patients treated with the combination of anti-CTLA-4 plus anti-PD-1 experience durable cancer control; however, immune related adverse events remain a problem^93,94^. As NeoAg vaccines have demonstrated favorable safety profiles^6,7^, combining NeoAg vaccines with single agent ICT may yield robust anti-tumor immunity with less toxicity than anti-CTLA-4 and anti-PD-1 combination ICT^69–72^. While we find that anti-CTLA-4 or anti-PD-1 can synergize with neo VAX in different tumor models when we give the first NeoAg vaccine and ICT mAb at the same time, the timing of treatment may impact the response in certain situations, as observed in other models and vaccine settings^95^. Although our approach targeting a single NeoAg in the Y1.7 model and three NeoAgs in the MC38 model was efficacious, it is likely that targeting multiple NeoAgs and possibly even shared, non-mutant antigens will be required in patients due to tumor heterogeneity and therapy induced-immunoediting, with at least some of the antigens targeted by the vaccine needing to be clonal NeoAgs^96,97^.

This study provides key insights into the transcriptional, molecular, and functional changes that occur within major immune cell populations within the TME following different forms of cancer immunotherapy and compliments ongoing human clinical studies of NeoAg vaccines. Although we did not fully elaborate on every specific immune cell population we profiled, our analyses were designed to interrogate the entire immune TME, and thus our study should additionally provide an important resource. The myeloid and lymphoid cell subsets and potential biomarkers we have described herein should inform the development of improved personalized NeoAg vaccines and combinatorial therapies in human patients.

## STAR★Methods

### Key resources Table S1

#### Mice

All mice used were on a C57BL/6 background. Wildtype (WT) C57BL/6J mice were purchased from Jackson Labs. All *in vivo* experiments used 8- to 12-week-old male or female mice (to match the sex and strain of the tumors). All mice were housed in a specific pathogen-free animal facility. All animal studies were performed in accordance with, and with the approval of the Institutional Animal Care and Use Committee (IACUC) of The University of Texas MD Anderson Cancer Center (Houston, TX).

#### Plasmids

Gene blocks for mAlg8, mItgb1, or mLama4 were purchased from Integrated DNA Technologies. Minigene constructs were cloned into the BglII site of pMSCV-IRES GFP (mAlg8 and mItgb1) or pMSCV (mLama4 and mItgb1) using the Gibson Assembly method (New England Biolabs). To generate neoantigen-expressing Y1.7 melanoma cell lines, constructs were transiently transfected into Phoenix Eco cells using Fugene (Promega). After 48 hours, viral supernatants were filtered and subsequently used for transfection of Y1.7 melanoma cell line. Y1.7 mLama4 ^MHC-I^.mItgb1 ^MHC-II^ (Y1.7LI) and Y1.7 mAlg8 ^MHC-I^.mItgb1 ^MHC-II^ (Y1.7AI) were sorted based on GFP positivity and clones were verified for neoantigen expression.

#### Tumor cell lines

The *Braf^V600E^ Cdkn2a^-/-^ Pten^-/-^* YUMM1.7 parental line was originally generated in a male GEMM on the C57BL/6 background as described^45^. Parental YUMM1.7 was purchased from ATCC (CRL-3362) and was modified to generate NeoAg-expressing Y1.7 lines. The MC38 line was obtained from B. Schreiber (Washington University in St. Louis School of Medicine). All tumor cell lines were found to be free of common mouse pathogens and Mycoplasma as assessed by IDEXX IMPACT I mouse pathogen testing [PCR evaluation for: Corynebacterium bovis, Corynebacterium sp. (HAC2), Ectromelia, EDIM, Hantaan, K virus, LCMV, LDEV, MAV1, MAV2, mCMV, MHV, MNV, MPV, MTV, MVM, Mycoplasma pulmonis, Mycoplasma sp., Polyoma, PVM, REO3, Sendai, TMEV] in December 2023. Tumor cell lines from the same cryopreserved stocks that were used in this study tested negative for Mycoplasma and were authenticated and found to be free of non-mouse cells as assessed by mouse cell STR profiling (IDEXX CellCheck mouse 19 plus Mycoplasma spp. testing) in December 2023.

#### Tumor transplantation

The *Braf^V600E^ Cdkn2a^-/-^ Pten^-/-^* YUMM1.7 parental melanoma line, Y1.7LI or Y1.7AI melanoma line, and the MC38 colorectal cancer line cells were propagated in R-10 plus BME media [RPMI media (HyClone) supplemented with 1% l-glutamine, 1% penicillin–streptomycin, 1% sodium pyruvate, 0.5% sodium bicarbonate, 0.1% 2-mercaptoethanol, and 10% heat-inactivated fetal calf serum (FCS) (HyClone) upon thawing, tumor lines were passaged 3 to 6 times before experimental use. Prior to injection, cells were washed extensively, resuspended at a concentration of 0.5 × 10^6^ cells (for YUMM1.7, Y1.7LI, and Y1.7AI) or 1.5 x 10^6^ cells (for MC38) in 150 µL of endotoxin-free PBS and 150 µL was injected subcutaneously into the flanks of recipient mice. Tumor cells were >90% viable at the time of injection as assessed by Trypan blue exclusion. Tumor growth was quantified by caliper measurements and expressed as the average of two perpendicular diameters. Lack of survival was defined as mouse death or mean tumor diameter size of 15 mm.

#### Tumor rechallenge

For tumor rechallenge, mice that rejected primary tumors after treatment with anti-CTLA-4, anti-PD-1, anti-CTLA-4 + anti-PD-1, or NeoAg vaccines were then rechallenged with same number of cells used in primary challenge with either the same tumor line used in the primary tumor challenge or a different tumor line as indicated at least 60 days after complete rejection of the primary tumor.

#### *In vivo* antibody treatments

For ICT treatment, YUMM1.7 parental, Y1.7LI, or Y1.7AI tumor-bearing mice were treated intraperitoneally with 200 μg of anti-CTLA-4 and/or anti-PD-1 on d. 3, 6, 9, 12, 18, and 22 or d. 7, 10, 13, 16, 22, and 28; or d. 12, 15, 18, 21, 27 and 33 post-tumor transplant. For controls, mice were injected with 200 μg of IgG2a isotype control antibodies. MC38 tumor-bearing mice were treated intraperitoneally with 200 μg of anti-CTLA-4 and/or anti-PD-1 on d. 12, 15, 18, and 22 post-transplant. For antibody depletion studies, 250 μg of control mAb, anti-CD4, or anti-CD8α was injected intraperitoneally into mice at d. −1 and every 7 days thereafter until day 20. CD4 and CD8 depletion was verified by flow cytometry analysis of surface-stained peripheral blood monocytes (PBMC) and intratumoral immune cells. For *in vivo* experiments, “*In vivo* Platinum”-grade antibodies that were verified to be free of mouse pathogens (IDEXX IMPACT I mouse pathogen testing) were purchased from Leinco Technologies: anti-PD-1 (rat IgG2a clone RMP1–14), anti-CTLA-4 (murine IgG2b clone 9D9), anti-CD4 (rat IgG2b clone GK1.5), anti-CD8α (rat IgG2b clone YTS169.4), and isotype controls (rat IgG2a clone 1–1, mouse IgG2a clone OKT3, or rat IgG2b clone 1–2).

#### Peptides

Mutant Lama4 8-mer (<u>V</u>GFNFRTL), mutant Lama4 SLP (QKISFFDGFE<u>V</u>GFNFRTLQPNGLLFYYT), mutant Adpgk SLP (HLELASMTN<u>M</u>ELMSSIVHQ), mutant Rpl18 SLP (KAGGKILTFD<u>R</u>LALESPK), mutant Dpagt1 SLP (EAGQSLVISASIIVFNL<u>L</u>ELEGDYR), mutant Alg8 8-mer (ITY<u>T</u>WTRL), OVA-I_257–264_ (SIINFEKL), mutant Itgb1 SLP (DDCWFYFTYSVNGY<u>N</u>EAIVHVVETPDCP), and OVA-II_323–339_ (ISQAVHAAHAEINEAGR) peptides were custom ordered from Peptide 2.0. All peptides were HPLC purified to >95% purity.

#### Vaccination

Y1.7LI or Y1.7AI tumor bearing male mice were vaccinated subcutaneously with 10 μg mLama4 or mAlg8 synthetic long peptide (SLP) in combination with 50 μg of VacciGrade™ high molecular weight Polyinosinic-polycytidylic acid (pI:C) (InvivoGen) in a total volume of 150 µL diluted in endotoxin-free PBS on d. 3, 9, and 15 or d. 7, 13, and 19 or on d. 12, 18, and 24 post tumor transplant. MC38 tumor bearing female mice were vaccinated subcutaneously with 20 μg of mAdpgk SLP plus 20 μg of mRpl18 SLP plus 20 μg of mDpagt1 plus 50 μg pI:C adjuvant or control vaccine composed of 40 μg of irrelevant HPV SLP + 50 μg of pI:C on d. 12 and 19 post-tumor transplant. For SLP, peptide sequence used for mLama4; QKISFFDGFE<u>VGFNFRTL</u>QPNGLLFYYT (epitope underlined), for mAlg8; AVG<u>ITYTWTRL</u>YASVLTGSLV (epitope underlined), for mAdpgk; HLELASMTNMELMSSIVHQ, for mRpl18; KAGGKILTFD***R***LALESPK and for mDpagt1; EAGQSLVISASIIVFNLLELEGDYR. mLama4 SLP served as a relevant SLP for the Y1.7LI line and an irrelevant SLP for the Y1.7AI line. mAlg8 served as a relevant SLP for the Y1.7AI line and an irrelevant SLP for the Y1.7LI tumor.

#### Tetramers

OVA-I (SIINFEKL)-H-2K^b^ (irrelevant control tetramer), mutant Alg8-H-2K^b^, and mutant Lama4-H-2K^b^ tetramers conjugated to PE or APC fluorophores, were obtained from the Baylor College of Medicine MHC Tetramer Production Facility.

#### Tumor and spleen harvest

Established tumors were excised from mice, minced, and treated with 1 mg/mL type IA collagenase (Sigma-Aldrich) in HBSS (Hyclone) for 45 minutes at 37°C. Cells were washed thrice. Red blood cells were lysed using ACK lysis buffer (Gibco). To remove aggregates and clumps, cells were passed through a 40-μm strainer. Spleens were harvested, crushed, and vigorously resuspended to make single-cell suspensions. To remove aggregates and clumps, cells were passed through a 70-μm strainer and subsequently through a 40-μm strainer.

#### TIL peptide restimulation

For peptide and PMA/ionomycin T-cell stimulation, cells from tumors, isolated as described above (see tumor and spleen harvest section), stained, and CD4 and CD8 T cells were sorted. For sorting CD4 and CD8 T cells, tumor cells were stained for 5 min at room temperature with 500 ng of Fc block (anti-CD16/32) and then stained with antibodies to CD45, CD3ε, CD4 or CD8α and Zombie NIR Viability dye in 100 µl of staining buffer. Cells were incubated for 30 minutes at 4°C. Live CD45^+^Cd3ε^+^CD4^+^ and live CD45^+^Cd3ε^+^CD8α^+^ were then sorted on a BD FACSAria II (BD Biosciences). Splenocytes harvested from naive mice and 100,000 splenocytes were then pulsed with 1 μM of various 8- or 9- or 17- or 28-mer peptides or simulated with 10 ng/mL of PMA (MilliporeSigma) and 1 μg/mL of ionomycin (Fisher) and 100,000 CD4 or CD8 TIL were subsequently added and incubated at 37 °C. Naive splenocytes added with or without CD4 or CD8 TIL, was included as control. After 1 h, BD GolgiPlug (BD Bioscience) was added in, and cells were incubated for an additional 5 h at 37 °C.

#### Tetramer staining

For tetramer staining, cells were stained for 5 min at room temperature with 500 ng of Fc block (anti-CD16/32). H-2K^b^ tetramers conjugated to PE (1:50) or APC (1:100) for mutated Alg8, mutated Lama4, or SIINFEKL were added to cells and incubated for 20 min at 37°C. Tetramer-stained cells were further stained with surface antibody for anti-CD45, anti-Thy1.2, anti-CD8α, anti-CD4, anti-PD-1, anti-TIM-3, and anti-LAG-3 antibody for 20 min at 4 °C.

#### Flow cytometry

For flow cytometry, cells were stained for 5 minutes at room temperature with rat anti-mouse CD16/32 (mouse BD Fc Block; clone 2.4G2, BD Biosciences) at 1 μg/million cells and then surface stained with flow antibodies for 20 minutes at 4°C. Surface antibodies were diluted in FACS staining buffer (PBS with 2% FCS, 2 mmol/L EDTA, and 0.05% NaN3; Sigma). Anti-mouse CD45-BV605, CD90.2/Thy1.2-PE-Cy7, anti-mouse CD8α-BV786, anti-mouse CD4-BV711, anti-mouse CD19-BV650, anti-mouse CD20-BV421, anti-mouse CD45R/B220-BUV395, anti-mouse Nkp46/CD335-FITC, anti-mouse γδ TCR-PE-Cy7, anti-mouse PD-1-BV421, anti-mouse TIM-3, anti-mouse LAG-3-PerCP-Cy5.5, anti-mouse CD3ε-APC , anti-mouse CD64-BV421, anti-mouse Ly6G-Alexa Fluor 700, anti-mouse CX3CR1-FITC, anti-mouse I-A/I-E-BV650, anti-mouse CD103-BV421, anti-mouse CD24-BV711, anti-mouse CD11c-BV786, anti-mouse CD11b-APC, anti-mouse F4/80-BUV395, anti-mouse CD64-APC, CD117-FITC, anti-mouse CD11b-PerCP-Cy5.5, anti-mouse PDCA-1/BST-2 BV650, anti-mouse CD172a APC, anti-mouse PDL1-PE, anti-mouse FcεRI-PE-Cy7 were used for surface staining at the indicated dilutions. Zombie NIR Viability dye was added at 1:500 during surface staining.

For intracellular staining, surface-stained cells were fixed and permeabilized with Fixation/Permeabilization Solution Kit (BD Bioscience). Fixed and permeabilized cells were then stained with anti-mouse CD206-PE-Cy7 and anti-mouse iNOS/NOS2-PE for 30 minutes at 4°C.

For FOXP3 staining, surface-stained cells were fixed and permeabilized using the eBioscience FOXP3/Transcription Factor Staining Buffer Set. Fixed and permeabilized cells were then stained with anti-mouse FOXP3-FITC for 30 minutes at 4°C.

For intracellular cytokine staining of lymphocytes, tumor cells were isolated and CD4 and CD8 T cells were sorted and added to peptide pulsed or PMA+Ionomycin stimulated splenocytes and incubated at 37°C for 6 hours with GolgiStop (BD Bioscience). Cells were then washed and stained for 5 minutes at room temperature with Fc block at 1 μg/million cells and then surface stained for 30 minutes at 4°C, and then fixed and permeabilized with BD Fixation and Permeabilization Kit. Fixed and permeabilized cells were then stained with anti-mouse IFN-γ-APC and anti-mouse TNF-PE-Cy7 for 30 minutes at 4°C. All flow cytometry was performed on an BD Fortessa X-20, BD LSR, BD Fortessa, and analyzed using FlowJo software. Gating strategy used is depicted in **Figure S15**.

### scRNAseq

#### Antibody hashing for multiplexing

Antibody hashing and multiplexing was utilized for scRNAseq/scTCRseq of NeoAg-specific CD8 T cells. For CD45^+^ scRNAseq experiments, antibody hashing and multiplexing was not performed. For analysis of NeoAg-specific CD8 T cells, cell and nuclei labeling were performed according to an adapted BioLegend cell hashing protocol (TotalSeq™-C Antibodies and Cell Hashing with 10x Single Cell 5’ Reagent Kit v1.1 Protocol, BioLegend). Single cell suspensions of harvested tumors from treated mice were resuspended in BioLegend Cell Staining Buffer containing Fc receptor block and stained with mLama4 PE and APC labelled tetramers for 20 min at 37°C. Tetramer-stained cells from control mAb, control VAX, and neo VAX treatment conditions were immediately surface stained by adding anti-CD90.2/Thy1.2-PE-Cy7 and anti-CD8α-BV786 antibodies and incubated for 20 min at 4°C. Tetramer-stained samples from anti-CTLA-4, anti-PD-1, and anti-CTLA-4 plus anti-PD-1 treated groups were incubated with mixture of surface stain (anti-CD90.2/Thy1.2-PE-Cy7 and anti-CD8α-BV786 antibodies) and barcoded antibodies with unique hashtags for each treatment condition [anti-CTLA-4: Hashtag 1 Total Seq™-C0301 anti-mouse Hashtag 1 Antibody; anti-PD-1: Hashtag 2 (Total Seq™-C0302 anti-mouse Hashtag 2 Antibody); anti-CTLA-4 + anti-PD-1 combination: Hashtag 3 (Total Seq™-C0303 anti-mouse Hashtag 3 Antibody)]. Hashtag antibodies were used at a concentration of 1 μg per 2 million cells. Staining with surface antibodies and hashtag antibodies was done for 30 min at 4°C. Cells were then washed 3X with BioLegend Cell Staining Buffer. Sorted mLama4 tetramer-specific CD8 T cells with unique hashtags (anti-CTLA-4, anti-PD-1, and anti-CTLA-4 + anti-PD-1 samples) were pooled for single-cell library generation and CITE-seq (cellular indexing of transcriptomes and epitopes by sequencing) through multiplexing. Separate libraries were generated for control mAb, control VAX, and neo VAX samples and, thus, these were not multiplexed.

#### scRNAseq with TCR and FBC sample Processing

For TCRseq of NeoAg-specific CD8 T cells, samples were hash tagged and processed as described in “antibody hashing” section above. Cells were counted on a Countess 3 FL automated cell counter (Life Technologies) and viabilities were determined using trypan blue exclusion. Cell capture processing and gene expression, TCR, and feature barcode library preparations were performed following 10X Genomics’ guidelines for 5’ scRNAseq which included TCR and cell surface marker detection [CG000330_Chromium Next GEM Single Cell 5’ v2 (Dual Index) with Feature Barcode technology-Rev F]. QC steps after cDNA amplification and library preparation steps were carried out by running ThermoFisher Qubit HS dsDNA Assay along with Agilent (Santa Clara, CA) HS DNA Bioanalyzer for concentration and quality assessments, respectively. Library sample concentrations were verified using qPCR using a KAPA Biosystems KAPA Library Quantification Kit prior to pooling. Libraries were normalized to 5 nM for pooling. The gene expression, TCR, and FBC libraries were pooled in a ratio 5:1:1 (where applicable-one sample out of four). The pool was sequenced using a NovaSeq6000 S4-XP,200-cycle flow cell lane. The run parameters used were 26 cycles for read 1, 90 cycles for read2, 10 cycles for index1, and 10 cycles for index2 as stipulated in the protocol mentioned above. Raw sequencing data (fastq file) was demultiplexed and analyzed using 10X Genomics Cell Ranger v.7.1.0 software utilizing standard default settings and the cellranger count command to generate html QC metrics and coupé/vloupe files for each sample.

#### CD45^+^ scRNAseq library generation

Droplet-based 5ʹ end massively parallel scRNAseq was performed by encapsulating sorted live CD45^+^ tumor-infiltrating cells into droplets and libraries were prepared using Chromium Next GEM Single-cell 5ʹ Reagent Kit v2 (10x Genomics) according to manufacturer’s protocol. The generated scRNAseq libraries were sequenced using an Illumina NovaSeq6000 S2 flow cell.

#### scRNAseq alignment, barcode assignment, and unique molecular identifier counting

The Cell Ranger Single-Cell Software Suite available at https://support.10xgenomics.com/single-cell-gene-expression/software/overview/welcome was used to perform sample demultiplexing, barcode processing, and single-cell 5ʹ counting. Cellranger mkfastq was used to demultiplex raw base call files from the NovaSeq6000 sequencer, into sample-specific fastq files. Files were demultiplexed with 81.9% to 97.1% perfect barcode match, and 90%+ q30 reads. Afterward, fastq files for each sample were processed with Cellranger count, which was used to align samples to mm10 genome, filtered, and quantified. For each sample, the recovered cells’ parameter was specified as 10,000 cells that we expected to recover for each individual library.

#### Preprocessing analysis with Seurat package

The Seurat pipeline was applied to each dataset following tutorial specifications from https://satijalab.org/seurat/articles/archive; version 4.3 and https://hbctraining.github.io/scRNA-seq_online/. Data from all groups were merged into a single Seurat object, and integration was performed using the reciprocal principal component analysis (PCA) workflow to identify integration anchors. After integration, genes that were expressed in fewer than 3 cells and cells that contained fewer than 500 transcripts (unique molecular identifiers; UMI) were excluded. Cells with more than 10% of mitochondrial transcripts were also excluded from analysis. The cutoffs used were set based on the characteristics of the cell population in each dataset. Data were normalized using LogNormalize method (counts for each cell divided by the total counts for that cell, multiplied by the scale factor of 10^4^ and natural-log transformed using log1p). PCA was performed on about 2,000 genes with PCA function. A uniform manifold approximation and projection (UMAP) dimensional reduction was performed on the scaled matrix (with most variable genes only) using the first 40 or 50 principal components (PCA) for mLama4 neoAg-specific CD8 T cells and CD45^+^ cells, respectively, to obtain a two-dimensional representation of the cell states. For clustering, we used the function FindClusters that implements SNN (shared nearest neighbor) modularity optimization–based clustering algorithm on 30 PCA components, leading to 33 clusters.

#### Identification of cluster-specific genes and marker-based classification

To identify marker genes, the FindAllMarkers function was used with likelihood-ratio test for single-cell gene expression. To characterize clusters, we used ImmGen database. For heat map representation, mean expression of markers inside each cluster was used. To compare gene expression for the clusters inside cohorts (e.g., T cells, macrophages) we used FindMarkers function to calculate average log2 fold change and identify differentially expressed genes between each pair of experimental conditions using a Wilcoxon rank-sum test for calculating P values and Bonferroni correction for Padj values.

#### T cell population analysis

To gain more insights into different immunotherapies-induced T cells remodeling in the TME, we subclustered activated T cells (excluding quiescent T cell clusters 10 and 12). Identification of most variable genes, PCA, UMAP, clustering, and marker selection analysis were performed as described above.

#### Gene set enrichment analysis (GSEA)

To identify if MSigDB hallmark gene sets are up-regulated or down-regulated between clusters and treatments, we performed gene set enrichment analysis. Fold-changes of gene expression between comparisons were calculated using Seurat R package v.4.3.0.1, and normalized enrichment scores as well as p-values of given gene sets were then estimated using the gage R package v.2.46.1.

#### Pseudo time trajectory analysis

To determine the potential lineage differentiation within CD4 T cell subpopulations, we used the Monocle3 R package to construct CD4 differentiation trajectories after specifying the corresponding cells as root nodes. Subsequently, graph test was used to find the pseudo time trajectory difference genes, and the obtained genes were used to plot the heat map.

#### scTCRseq Analysis

scTCRseq data for mLama4 NeoAg-specific CD8 T cells for each sample were processed by CellRanger. For TCR selection a meta data .csv was exported after initial QC and imported into R and TCR clones were further analyzed in combination with the corresponding scRNAseq data using the R packages scRepertoire v.2.0.0 and Seurat v.4.3.0.1. mLama4 NeoAg-specific CD8 T cells with at least one productive TCR alpha or beta chain or both and separately, with paired TCR alpha beta chains were considered for precise identification of TCRs. The total number of this NeoAg-specific CD8 T cells with TCR alpha and beta pair set was 15,668 from 17,492 total TCR (TCR with single alpha or beta or both alpha and beta pair) [control mAb: 3,118 from 3,539 total TCR; anti-CTLA-4: 1,208 from 1,394; anti-PD-1: 1,162 from 1,283; anti-CTLA-4 plus anti-PD-1: 657 from 790; control VAX: 4,986 from 5,622; neo VAX: 4,537 from 4,864]. The Shannon Index of diversity was calculated with the R package scRepertoire (V.2.0.0) (https://www.borch.dev/uploads/screpertoire/articles/clonal_diversity). Downsampling to the smallest repertoire size and bootstrapping to return the mean diversity estimates was performed with the number of calculations set to the default of 100.

#### Statistical analysis

Samples were compared using an unpaired, two-tailed Student t test, two-way ANOVA, or log-rank (Mantel–Cox) test unless specified otherwise.

## Supporting information

Key Sources Table

## Data and software availability

Data files for the sequencing data reported in this article will be deposited in the Gene Expression Omnibus (GEO) database and made publicly available at the time of publication. Software used in this study is available online: current version of Cell Ranger: https://support.10xgenomics.com/single-cell-gene-expression/software/downloads/latest; Seurat 4. 3.0.1: https://satijalab.org/seurat/; ggplot2 3.3.3: https://ggplot2.tidyverse.org/index.html; scRepertoire 2.0.0: https://www.borch.dev/uploads/screpertoire/; and ImmGen: https://www.immgen.org. All other data generated in this study are available within the article and its Supplementary Data files, will be provided upon request at the time of publication, and/or will made publicly available at the time of publication via deposition in appropriate databases.

## Acknowledgements

S. Keshari was a Balzan Postdoctoral Research Fellow supported by The International Balzan Prize Foundation. M.M. Gubin is a Cancer Prevention and Research Institute of Texas (CPRIT) Scholar in Cancer Research and an Andrew Sabin Family Fellow. This work was supported by CPRIT (Recruitment of First-Time Tenure-Track Faculty Members; RR190017), an Andrew Sabin Family Foundation Fellowship, Parker Institute for Cancer Immunotherapy (PICI) Bridge Scholar Award, University of Texas (UT) Rising Stars Award, and the University of Texas MD Anderson Cancer Center (MDACC) Support Grant (CCSG) New Faculty Award supported by the National Institutes of Health (NIH)/National Cancer Institute (NCI) (P30CA016672) to M.M. Gubin; and NIH/NCI U01CA247760 to K. Chen. K.H. Hu is a CPRIT Scholar in Cancer Research and a PICI and V Foundation Bridge Scholar. K.E. Pauken is supported by an Andrew Sabin Family Foundation Fellowship, a Melanoma SPORE Developmental Research Program Grant, and a UT Rising STARs Award. The Flow Cytometry and Cellular Imaging Core Facility was supported in part by MDACC and NIH/NCI Core grant P30CA016672. scRNAseq was performed by the MDACC Advanced Technology Genomics Core (ATGC) Facility supported by an NCI Core grant [CA016672 (ATGC)]. We would like to thank David Pollock at MDACC ATGC Facility for assistance with scRNAseq. We would like to thank the Baylor College of Medicine MHC Tetramer Core and thank the core director, X. Lily Wang for production of MHC tetramers used in this study. We would like to thank Prachi Sao (MDACC) for assistance with deconvolution of multiplexed hashtagged scRNAseq samples. We would like to thank Mehdi Chaib, (MDACC) for providing feedback to the manuscript. The authors thank all members of the Gubin lab for helpful discussions and technical support.

## Authors’ Contributions

S. Keshari: Conceptualization, data curation, investigation, visualization, methodology, data analysis, writing–original draft, writing–review and editing. A.S. Shavkunov: Conceptualization, data curation, investigation, data analysis, writing–review and editing. Q. Miao: Conceptualization, data curation, investigation, visualization, data analysis, writing–review and editing. A. Saha: Data curation, investigation, visualization, writing–review and editing. C.D. Williams: data curation, investigation, visualization, writing–review and editing. A.M. Highsmith: data curation, investigation, visualization, writing–review and editing. J.E. Pineda: data curation, investigation, visualization, writing–review and editing. E. Alspach: Resources, formal analysis, investigation, visualization, writing–review and editing. K. Hu: Formal analysis, investigation, visualization, writing–review and editing. K.E. Pauken: Formal analysis, investigation, visualization, writing– review and editing. K. Chen: Resources, formal analysis, investigation, visualization, writing–review and editing. M.M. Gubin: Conceptualization, resources, data curation, formal analysis, supervision, validation, investigation, methodology, writing–original draft, writing–review and editing.

