## Supplemental File for "Neoantigen Cancer Vaccines and Different Immune Checkpoint Therapies Each Utilize Both Converging and Distinct Mechanisms that in Combination Enable Synergistic Therapeutic Efficacy"

### Figure S1

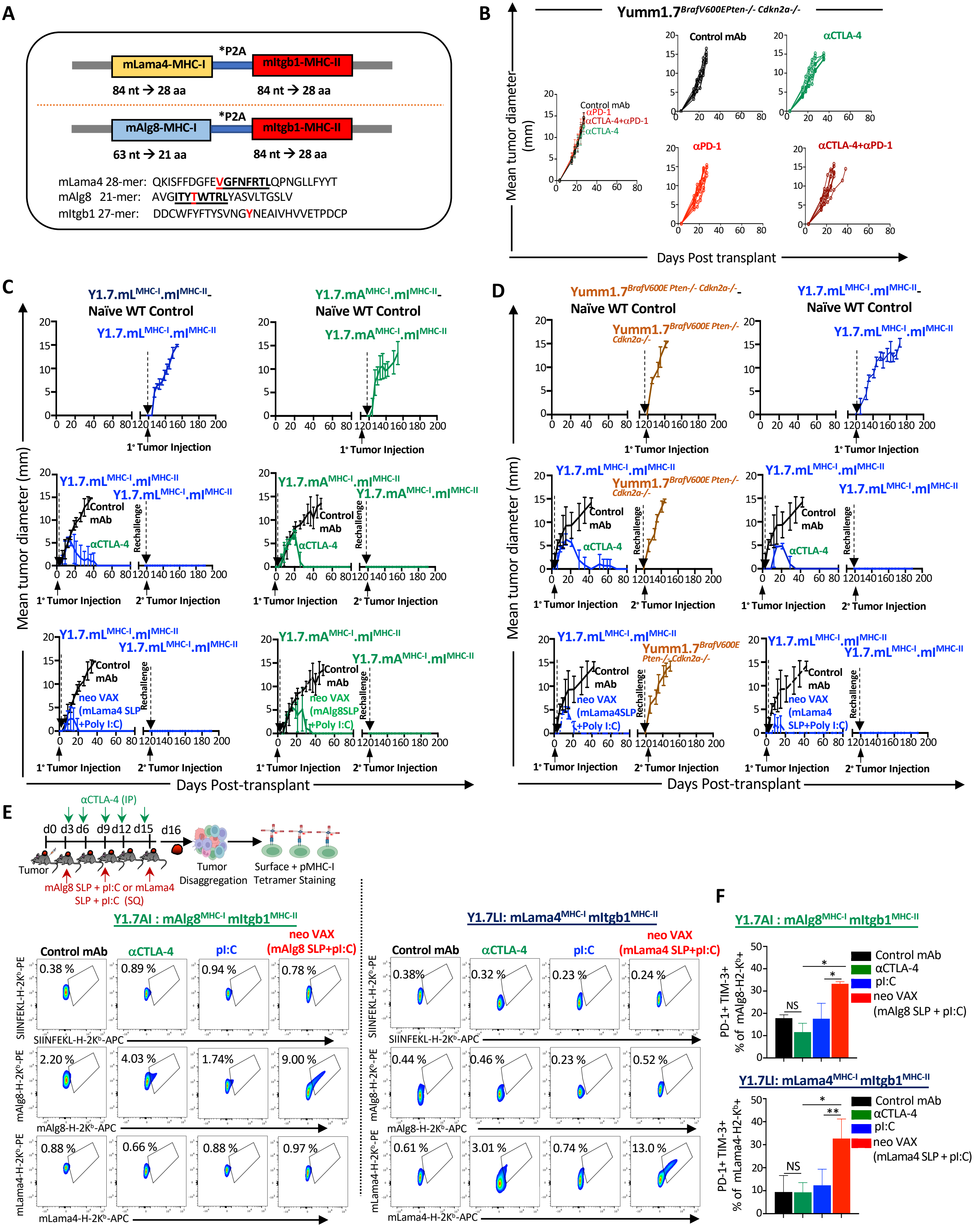

A
mLama4-MHC-I
*P2A
mItgb1-MHC-II
84 nt → 28 aa
84 nt → 28 aa
mAlg8-MHC-I
*P2A
mItgb1-MHC-II
63 nt → 21 aa
84 nt → 28 aa
mLama4 28-mer: QKISFFDGFEVGFNFRTLQPNGLLFYYT
mAlg8 21-mer: AVGITYTWTRLYASVLTGSLV
mItgb1 27-mer: DDCWFYFTYSVNGYNEAIVHVVETPDCP
B
Yumm1.7BrafV600EPten-/- Cdkn2a-/-
Mean tumor diameter (mm)
Control mAb
αPD-1
αCTLA-4+αPD-1
αCTLA-4
Days Post transplant
C
Y1.7.mLMHC-I.mIMHC-II- Naïve WT Control
Y1.7.mAMHC-I.mIMHC-II- Naïve WT Control
Y1.7.mLMHC-I.mIMHC-II
Y1.7.mAMHC-I.mIMHC-II
1° Tumor Injection
2° Tumor Injection
Rechallenge
Control mAb
αCTLA-4
neo VAX (mLama4 SLP +Poly I:C)
neo VAX (mAlg8SLP +Poly I:C)
Mean tumor diameter (mm)
Days Post-transplant
D
Yumm1.7BrafV600E Pten-/- Cdkn2a-/- Naïve WT Control
Y1.7.mLMHC-I.mIMHC-II- Naïve WT Control
Yumm1.7BrafV600E Pten-/- Cdkn2a-/-
Y1.7.mLMHC-I.mIMHC-II
1° Tumor Injection
2° Tumor Injection
Rechallenge
Control mAb
αCTLA-4
neo VAX (mLama4SLP +Poly I:C)
neo VAX (mLama4 SLP+Poly I:C)
Mean tumor diameter (mm)
Days Post-transplant
E
αCTLA-4 (IP)
d0 d3 d6 d9 d12 d15 d16
Tumor
mAlg8 SLP + pI:C or mLama4 SLP + pI:C (SQ)
Tumor Disaggregation
Surface + pMHC-I Tetramer Staining
Y1.7AI : mAlg8MHC-I mItgb1MHC-II
Y1.7LI: mLama4MHC-I mItgb1MHC-II
Control mAb
αCTLA-4
pI:C
neo VAX (mAlg8 SLP+pI:C)
neo VAX (mLama4 SLP+pI:C)
SIINFEKL-H-2Kb-PE
SIINFEKL-H-2Kb-APC
mAlg8-H-2Kb-PE
mAlg8-H-2Kb-APC
mLama4-H-2Kb-PE
mLama4-H-2Kb-APC
0.38 %
0.89 %
0.94 %
0.78 %
2.20 %
4.03 %
1.74%
9.00 %
0.88 %
0.66 %
0.88 %
0.97 %
0.38%
0.32 %
0.23 %
0.24 %
0.44 %
0.46 %
0.23 %
0.52 %
0.61 %
3.01 %
0.74 %
13.0 %
F
Y1.7AI : mAlg8MHC-I mItgb1MHC-II
PD-1+ TIM-3+ % of mAlg8-H2-Kb+
Control mAb
αCTLA-4
pI:C
neo VAX (mAlg8 SLP + pI:C)
NS
*
Y1.7LI: mLama4MHC-I mItgb1MHC-II
PD-1+ TIM-3+ % of mLama4-H2-Kb+
neo VAX (mLama4 SLP + pI:C)
**

**Figure S1. NeoAg vaccines and ICT induces long-term tumor protection in Y1.7AI and Y1.7LI melanoma models. Related to Figure 1.**

(A) Depiction of NeoAg minigenes used to express NeoAgs in the parental *Braf*$^{V600E}$ *Pten*$^{-/-}$ *Cdkn2a*$^{-/-}$ YUMM1.7 melanoma line, along with peptide sequences encoded by minigenes. mLama4 or mAlg8 and mItgb1 NeoAgs were separated by 2A peptides that induce ribosomal skipping during translation. (B) Tumor growth in WT C57BL/6J mice transplanted with parental *Braf*$^{V600E}$ *Pten*$^{-/-}$ *Cdkn2a*$^{-/-}$ YUMM1.7 melanoma cells and treated with control mAb, anti-CTLA-4, anti-PD-1 or anti-CTLA4 + anti-PD-1 combination immune checkpoint therapy (ICT) on d. 3, 6, 9, 12, 18, 24 post tumor-transplant. (C) WT C57BL/6J mice were transplanted with Y1.7 mA$^{MHC-I}$.mI$^{MHC-II}$ (Y1.7AI) and Y1.7 mL$^{MHC-I}$.mI$^{MHC-II}$ (Y1.7LI) melanoma cells and treated with control mAb or anti-CTLA-4 on d. 3, 6, 9, 12, 18, 24 or mAlg8 NeoAg (relevant for Y1.7AI) synthetic long peptide (SLP) + poly I:C (pI:C) or mLama4 NeoAg (relevant for Y1.7LI) synthetic long peptide (SLP) + pI:C on d. 3, 9, 15. Mice were rechallenged with same tumor used for initial tumor challenge at least 60 days post-rejection of primary tumor. Naïve WT C57BL/6J mice transplanted with Y1.7AI or Y1.7LI tumor without any treatment were included as controls, indicating cell line preps used in rechallenge experiments were capable of tumor formation. (D) WT C57BL/6J mice were transplanted with Y1.7LI melanoma cells and treated with anti-CTLA-4 ICT on d. 3, 6, 9, 12, 18, 24 or mLama4 NeoAg SLP + pI:C on d. 3, 9, 15. Mice were rechallenged with either with same tumor used for initial tumor challenge (Y1.7LI) or parental *Braf*$^{V600E}$ *Pten*$^{-/-}$ *Cdkn2a*$^{-/-}$ YUMM1.7 at least 60 days post-rejection of primary tumor. Naïve WT C57BL/6J mice transplanted with either Y1.7LI or parental YUMM1.7 without any treatment were included as controls. (E**)** Representative flow cytometry plots displaying mAlg8 or mLama4 tetramer-specific CD8 T cells in Y1.7AI and Y1.7LI tumors treated with control mAb, anti-CTLA-4, pI:C, mAlg8 SLP + pI:C NeoAg vaccine (for Y1.7AI) or mLama4 SLP + pI:C NeoAg vaccine (for Y1.7LI) and harvested on d. 16 post-tumor transplant. mAlg8-H2-K$^{b}$, mLama4-H2-K$^{b}$, or SIINFEKL-H2-K$^{b}$ (irrelevant control) tetramers were labeled with PE and APC. Dot plots are gated on live CD45$^{+}$ Thy1.2$^{+}$ CD8 T cells. (F) Co-expression of PD-1 and TIM-3 on mAlg8- or mLama4-specific CD8 T cells in Y1.7AI and Y1.7LI tumors treated with control mAb, anti-CTLA-4, pI:C alone, mAlg8 SLP + pI:C NeoAg vaccine (for Y1.7AI), or mLama4 SLP + pI:C NeoAg vaccine (for Y1.7LI). Tumor growth data in (B), (C) and (D) are presented as individual mouse tumor growth as mean tumor diameter and are representative of three independent experiments.

Figure S2

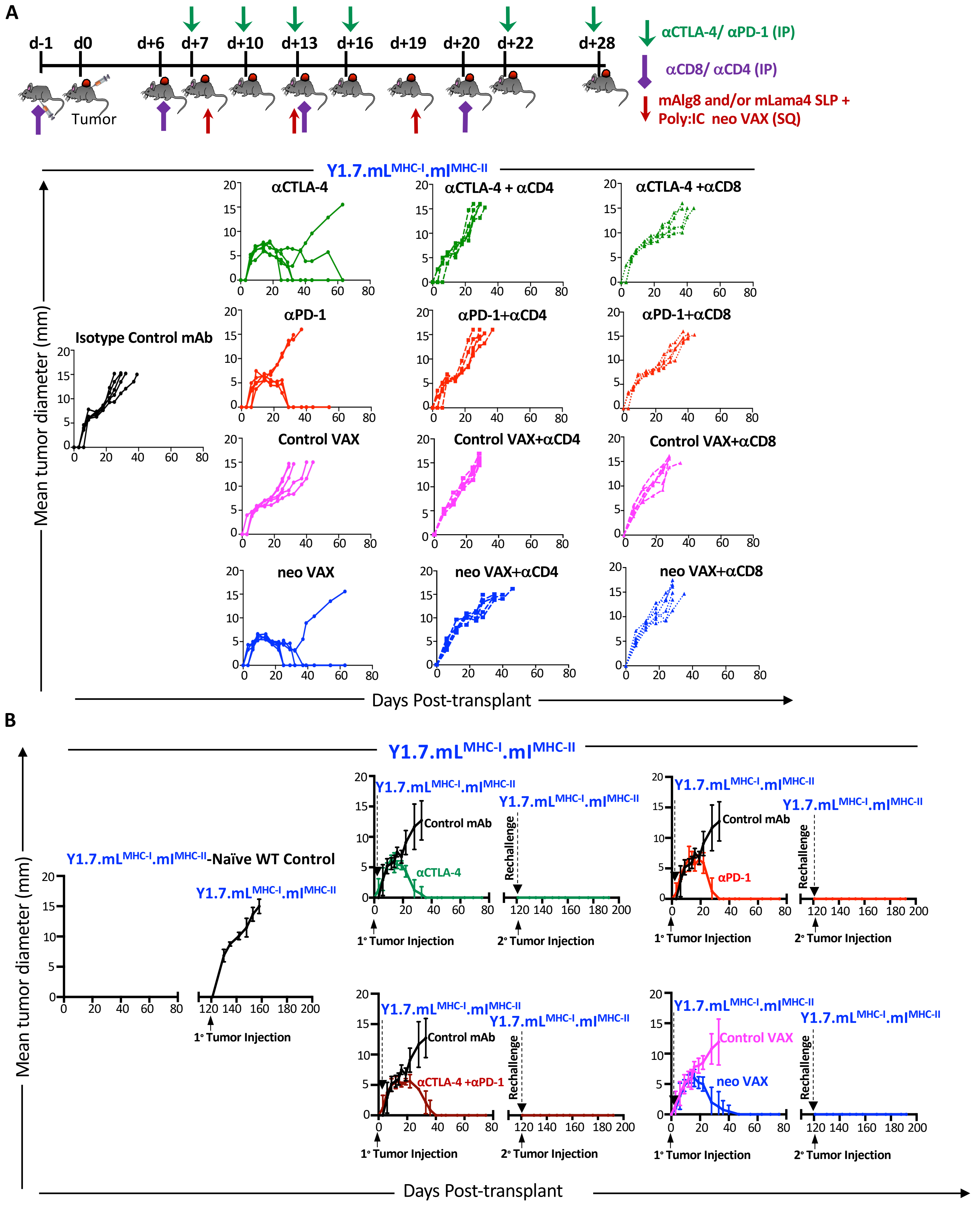
A
d-1
d0
d+6
d+7
d+10
d+13
d+16
d+19
d+20
d+22
d+28
Tumor
αCTLA-4/ αPD-1 (IP)
αCD8/ αCD4 (IP)
mAlg8 and/or mLama4 SLP +
Poly:IC neo VAX (SQ)
Y1.7.mL^MHC-I.mI^MHC-II
Mean tumor diameter (mm)
Isotype Control mAb
αCTLA-4
αCTLA-4 + αCD4
αCTLA-4 +αCD8
αPD-1
αPD-1+αCD4
αPD-1+αCD8
Control VAX
Control VAX+αCD4
Control VAX+αCD8
neo VAX
neo VAX+αCD4
neo VAX+αCD8
Days Post-transplant
B
Y1.7.mL^MHC-I.mI^MHC-II
Y1.7.mL^MHC-I.mI^MHC-II-Naïve WT Control
Control mAb
αCTLA-4
αPD-1
αCTLA-4 +αPD-1
Control VAX
neo VAX
Rechallenge
1° Tumor Injection
2° Tumor Injection
Mean tumor diameter (mm)
Days Post-transplant

**Figure S2. NeoAg vaccines and ICT induce long-term tumor protection in Y1.7LI melanoma models in a T cell-dependent manner. Related to Figure 1.**

(A) Y1.7LI tumor growth in WT C57BL/6J mice treated with isotype control mAb, anti-CD4, or anti-CD8α mAbs on d. -1, 6, 13, 20 and anti-CTLA-4 or anti-PD-1 on d. 7, 10, 13, 16, 22, 28 or irrelevant mAlg8 SLP + pI:C (Control VAX) or relevant mLama4 SLP + pI:C (neo VAX) on d. 7, 13, 19. (B) WT C57BL/6J mice transplanted with Y1.7LI melanoma cells were treated with control mAb, anti-CTLA-4, anti-PD-1, anti-CTLA-4 + anti-PD-1, irrelevant (for Y1.7LI) mAlg8 SLP + pI:C (Control VAX), or relevant mLama4 SLP + pI:C (neo VAX) starting on d. 7 post tumor-transplant, and subsequently on d. 10, 13, 16, 22, 28 for ICT and d. 13, 19 for NeoAg vaccines. Mice were rechallenged with the same tumor line used for initial tumor challenge (Y1.7LI) at least 60 days post-rejection of primary tumor. Naïve WT C57BL/6J mice transplanted with Y1.7LI tumor without any treatment was included as a control. Tumor growth data in (A) and (B) are presented as individual mouse tumor growth as mean tumor diameter and are representative of three independent experiments.

### Figure S3

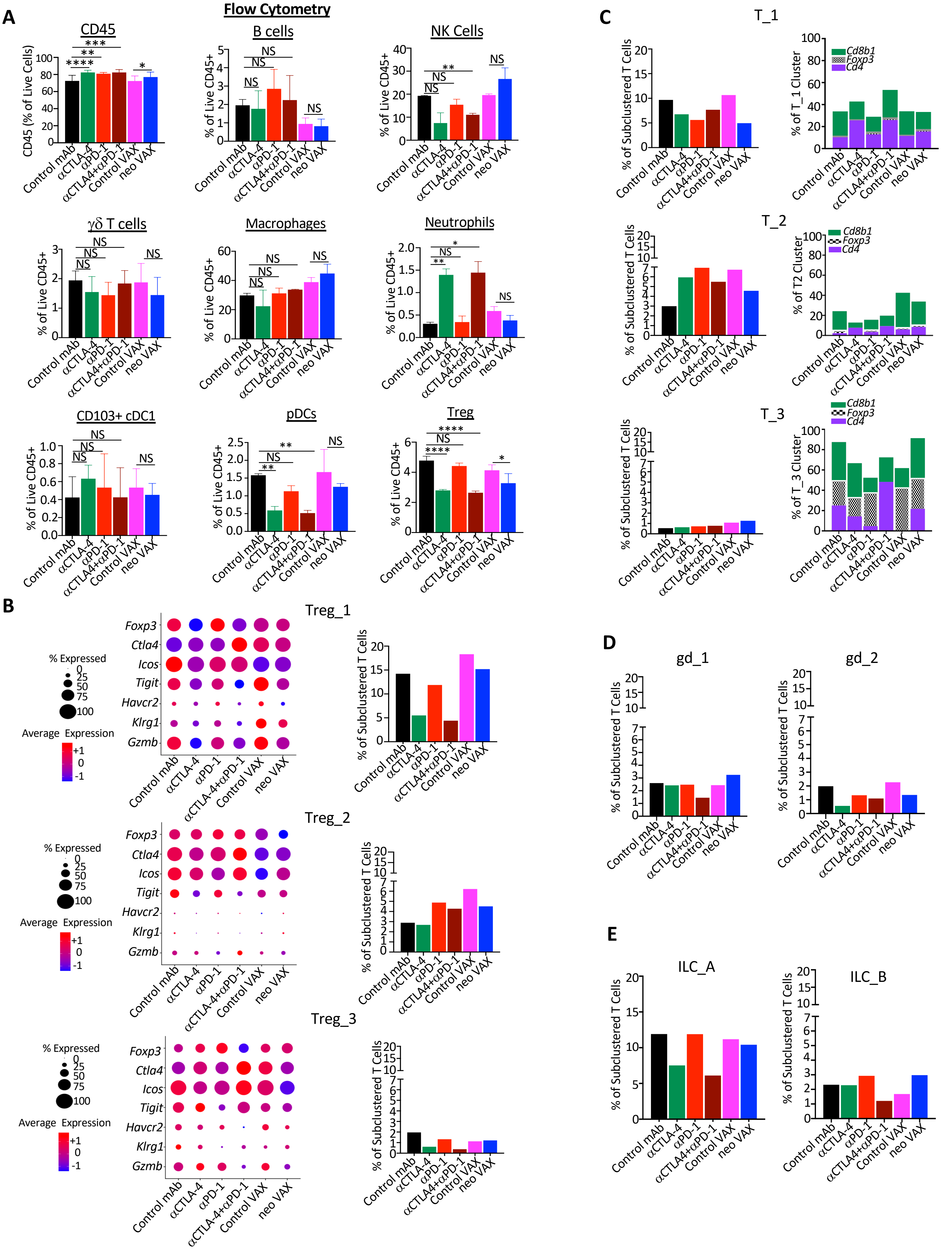
A
Flow Cytometry
CD45
B cells
NK Cells
γδ T cells
Macrophages
Neutrophils
CD103+ cDC1
pDCs
Treg
CD45 (% of Live Cells)
% of Live CD45+
Control mAb
αCTLA-4
αPD-1
αCTLA4+αPD-1
Control VAX
neo VAX
B
Treg_1
Treg_2
Treg_3
% Expressed
Average Expression
Foxp3
Ctla4
Icos
Tigit
Havcr2
Klrg1
Gzmb
% of Subclustered T Cells
C
T_1
T_2
T_3
Cd8b1
Foxp3
Cd4
% of T_1 Cluster
% of T2 Cluster
% of T_3 Cluster
D
gd_1
gd_2
E
ILC_A
ILC_B

**Figure S3. Flow cytometry and scRNAseq analysis of Y1.7LI intratumoral lymphoid and myeloid populations. Related to Figure 2.**

(A) Graph of flow cytometry data displaying intratumoral lymphoid and myeloid cells as a percentage of intratumoral live or live $CD45^+$ cells in Y1.7LI tumors treated with control mAb, anti-CTLA-4, anti-PD-1, anti-CTLA-4 + anti-PD-1, irrelevant (for Y1.7LI) mAlg8 SLP + pI:C (control VAX), or relevant mLama4 SLP + pI:C (neo VAX) beginning on d. 7 post-tumor transplant and harvested on d. 15. (B) Dot plot depicting expression level and percent of cells expressing *Foxp3*, *Ctla4*, *Icos, Tigit, Havcr2* (TIM-3)*, Klrg1, Gzmb* and graph displaying frequency of regulatory T cell (Treg) scRNAseq clusters by treatment condition. (C) Graph displaying mixed T cell clusters represented as percentage of total subclustered T cells and percentage of $Foxp3^+$ CD4 Tregs, conventional CD4 T cells, or CD8 T cells in clusters T_1, T_2, and T_3 by treatment condition. (D) Graph displaying $\gamma\delta$ T cell clusters represented as percentage of total subclustered T cells by treatment condition. (E) Graph displaying ILC clusters represented as percentage of total subclustered T cells by treatment condition. Bar graphs in (A) display mean ± SEM and are representative of at least three independent experiments (**P* < 0.05, ***P* < 0.01, ****P* < 0.005, *****P* < 0.0001, NS, not significant, unpaired t test).

**Figure S4**

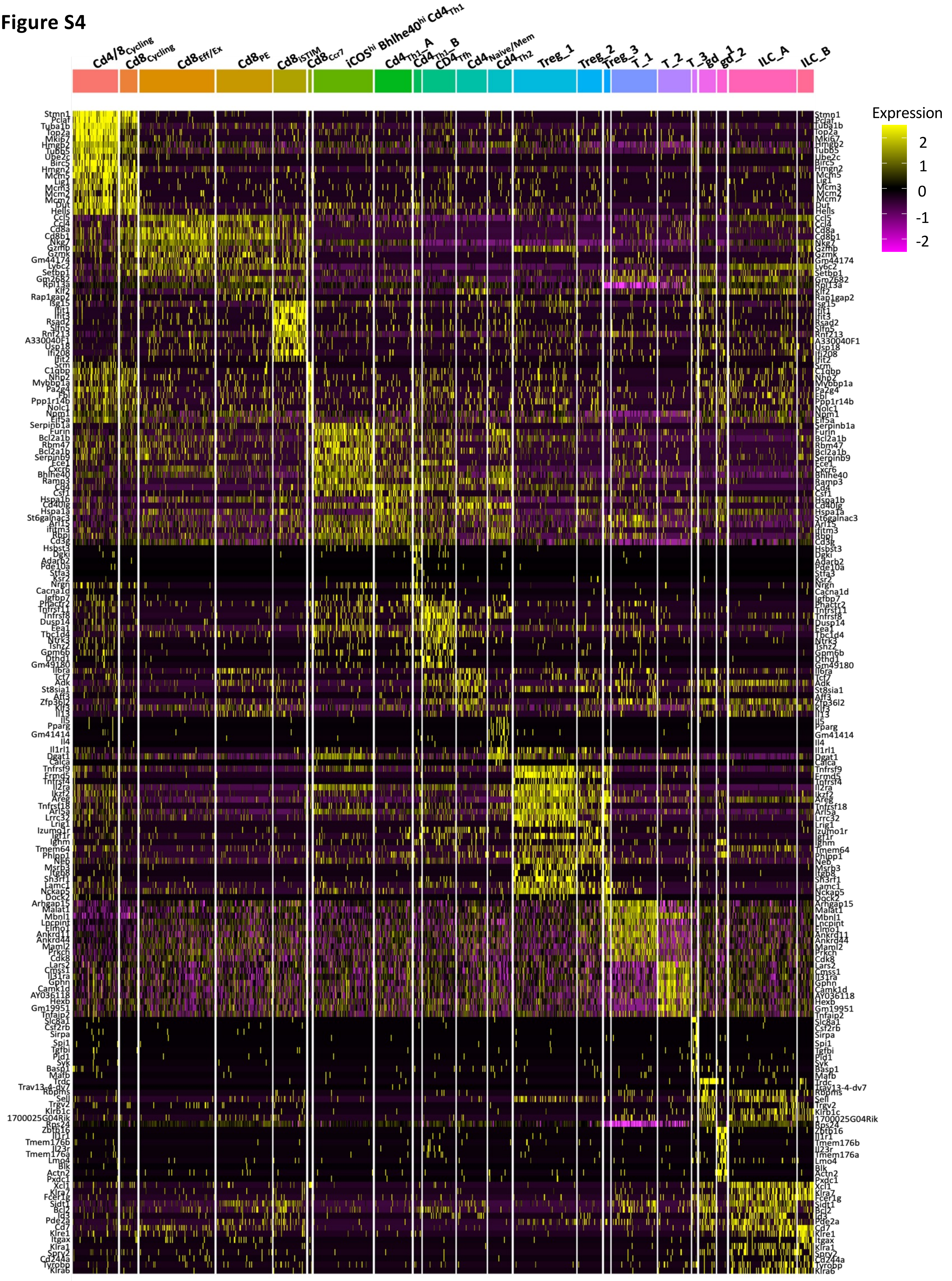

Cd4/8Cycling
Cd8Cycling
Cd8Eff/Ex
Cd8PE
Cd8ISTIM
Cd8Ccr7
iCOShi Bhlhe40hi Cd4Th1
Cd4Th1_A
Cd4Th1_B
CD4Tfh
Cd4Naive/Mem
Cd4Th2
Treg_1
Treg_2
Treg_3
T_1
T_2
T_3
gd_1
gd_2
ILC_A
ILC_B
Expression
2
1
0
-1
-2
Stmn1
Pclaf
Tuba1b
Top2a
Mki67
Hmgb2
Tubb5
Ube2c
Birc5
Hmgn2
Mcm5
Lig1
Mcm3
Mcm2
Mcm7
Dut
Hells
Ccl5
Ccl4
Cd8a
Cd8b1
Nkg7
Gzmb
Gzmk
Gm44174
Ly6c2
Setbp1
Gm2682
Rpl13a
Klf2
Rap1gap2
Isg15
Ifit1
Ifit3
Rsad2
Slfn5
Rnf213
A330040F1
Usp18
Ifi208
Ifit2
Srm
C1qbp
Nhp2
Mybbp1a
Pa2g4
Fbl
Ppp1r14b
Nolc1
Npm1
Eif5a
Serpinb1a
Furin
Bcl2a1b
Rbm47
Bcl2a1b
Serpinb9
Ece1
Cxcr6
Bhlhe40
Ramp3
Cd4
Csf1
Hspa1b
Cd40lg
Hspa1a
St6galnac3
Arl15
Ifitm3
Rbpj
Cd3g
Hsbst3
Dgki
Adarb2
Pde10a
Stfa3
Ksr2
Nrgn
Cacna1d
Igfbp7
Phactr2
Tnfrsf11
Tnfrsf8
Dusp14
Eea1
Tbc1d4
Ntrk3
Tshz2
Gpm6b
Dthd1
Gm49180
Il6ra
Tcf7
Adk
St8sia1
Aff3
Zfp36l2
Klf3
Il13
Il5
Pparg
Gm41414
Il4
Il1rl1
Dgat1
Calca
Tnfrsf9
Frmd5
Tnfrsf4
Il2ra
Ikzf2
Areg
Tnfrsf18
Arl5a
Lrrc32
Lrig1
Izumo1r
Igf1r
Ighm
Tmem64
Phlpp1
Neb
Msrb3
Itgb8
Sh3rf1
Lamc1
Nckap5
Dock2
Arhgap15
Malat1
Mbnl1
Lncpint
Elmo1
Ankrd11
Ankrd44
Maml2
Prkch
Cdk8
Lars2
Cmss1
Il31ra
Gphn
Camk1d
AY036118
Hexb
Gm19951
Tnfaip2
Slc8a1
Csf2rb
Sirpa
Spi1
Tgfbi
Pld1
Syk
Basp1
Mafb
Trdc
Trav13-4-dv7
Rbpms
Sell
Trgv2
Klrb1c
1700025G04Rik
Rps24
Zbtb16
Il1r1
Tmem176b
Il23r
Tmem176a
Lmo4
Blk
Actn2
Pxdc1
Xcl1
Klra7
Fcer1g
Sidt1
Bcl2
Id3
Pde2a
Cd7
Klre1
Itgax
Klra1
Spry2
Cd244a
Tyrobp
Klra6

**Figure S4. Heat map of the top 10 most DEG across T cell/ILC clusters (see also Figure 2D). Related to Figure 2.**

### Figure S5

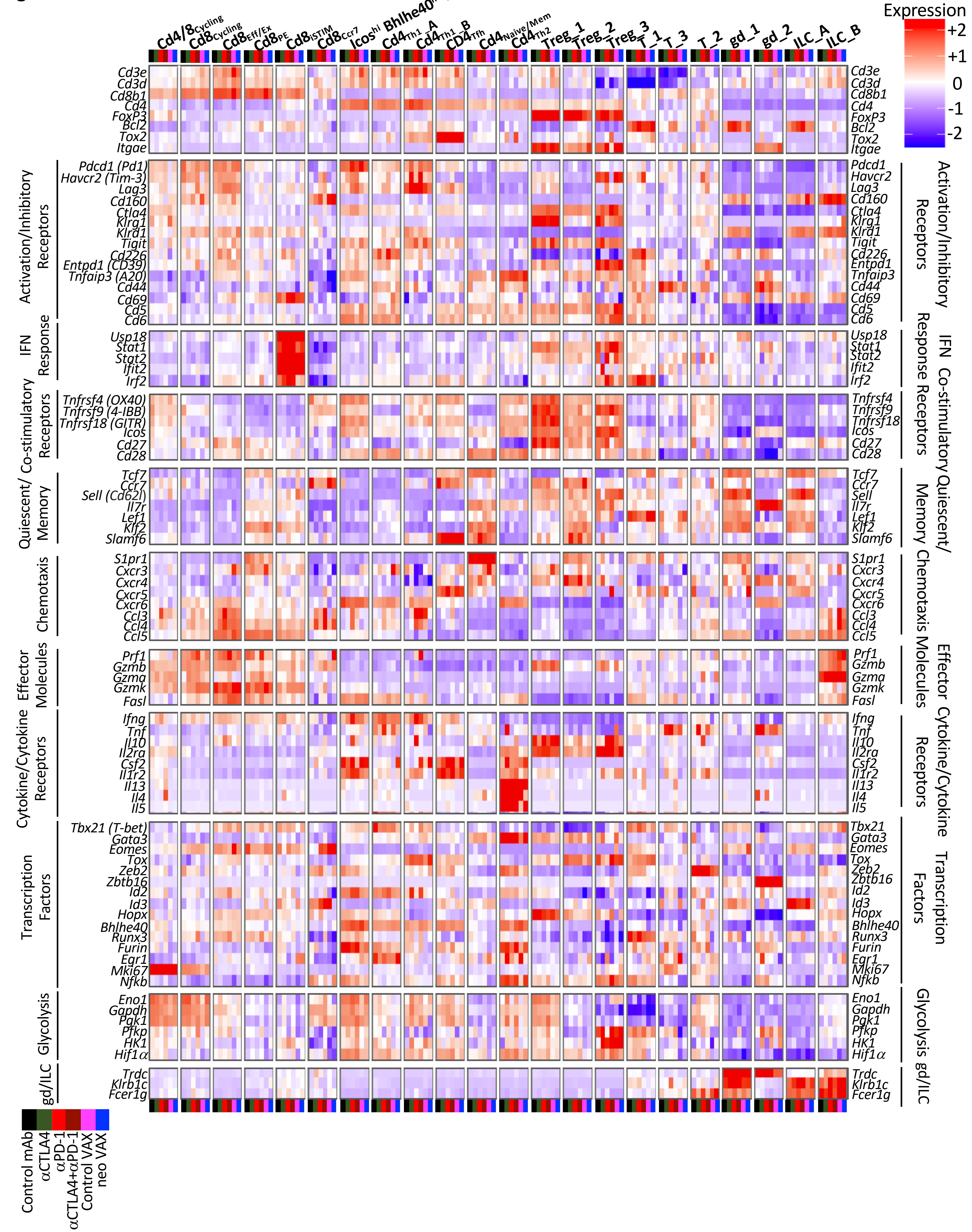
Expression
+2
+1
0
-1
-2
Cd4/8Cycling
Cd8Cycling
Cd8Eff/Ex
Cd8PE
Cd8ISTIM
Cd8Ccr7
Icoshi Bhlhe40hi Cd4Th1
Cd4Th1_A
Cd4Th1_B
CD4Tfh
Cd4Naive/Mem
Cd4Th2
Treg_1
Treg_2
Treg_3
T_1
T_3
T_2
gd_1
gd_2
ILC_A
ILC_B
Cd3e
Cd3d
Cd8b1
Cd4
FoxP3
Bcl2
Tox2
Itgae
Activation/Inhibitory Receptors
Pdcd1 (Pd1)
Havcr2 (Tim-3)
Lag3
Cd160
Ctla4
Klrg1
Klrd1
Tigit
Cd226
Entpd1 (CD39)
Tnfaip3 (A20)
Cd44
Cd69
Cd5
Cd6
IFN Response
Usp18
Stat1
Stat2
Ifit2
Irf2
Co-stimulatory Receptors
Tnfrsf4 (OX40)
Tnfrsf9 (4-IBB)
Tnfrsf18 (GITR)
Icos
Cd27
Cd28
Quiescent/ Memory
Tcf7
Ccr7
Sell (Cd62l)
Il7r
Lef1
Klf2
Slamf6
Chemotaxis
S1pr1
Cxcr3
Cxcr4
Cxcr5
Cxcr6
Ccl3
Ccl4
Ccl5
Effector Molecules
Prf1
Gzmb
Gzma
Gzmk
Fasl
Cytokine/Cytokine Receptors
Ifng
Tnf
Il10
Il2ra
Csf2
Il1r2
Il13
Il4
Il5
Transcription Factors
Tbx21 (T-bet)
Gata3
Eomes
Tox
Zeb2
Zbtb16
Id2
Id3
Hopx
Bhlhe40
Runx3
Furin
Egr1
Mki67
Nfkb
Glycolysis
Eno1
Gapdh
Pgk1
Pfkp
HK1
Hif1α
gd/ILC
Trdc
Klrb1c
Fcer1g
Control mAb
αCTLA4
αPD-1
αCTLA4+αPD-1
Control VAX
neo VAX

**Figure S5. Heat map displaying normalized expression of select genes in each T cell/ILC cluster by treatment condition. Related to Figure 2.**

**Figure S6**

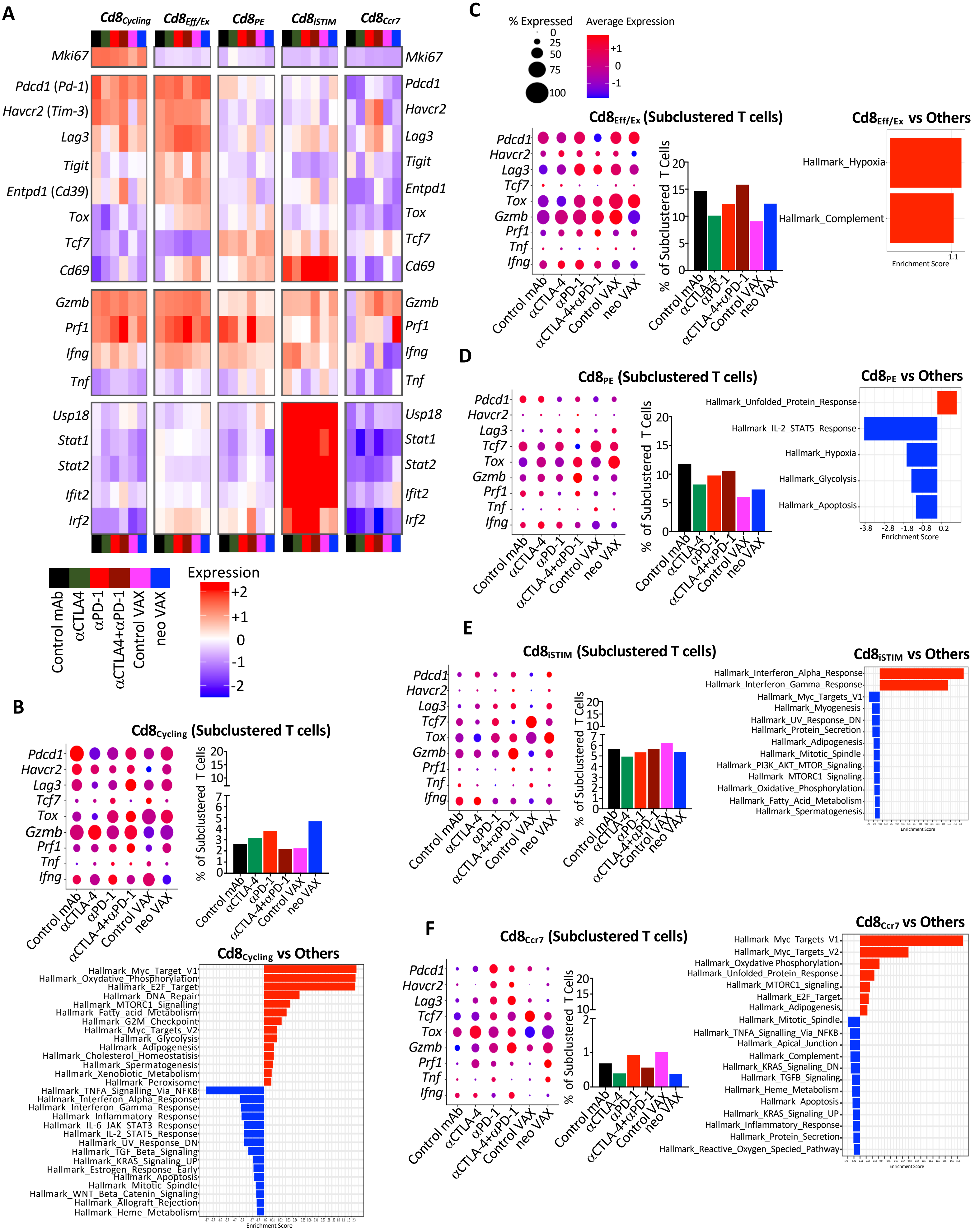
A
Cd8Cycling
Cd8Eff/Ex
Cd8PE
Cd8iSTIM
Cd8Ccr7
Mki67
Pdcd1 (Pd-1)
Havcr2 (Tim-3)
Lag3
Tigit
Entpd1 (Cd39)
Tox
Tcf7
Cd69
Gzmb
Prf1
Ifng
Tnf
Usp18
Stat1
Stat2
Ifit2
Irf2
Pdcd1
Havcr2
Entpd1
Control mAb
αCTLA4
αPD-1
αCTLA4+αPD-1
Control VAX
neo VAX
Expression
+2
+1
0
-1
-2
B
Cd8Cycling (Subclustered T cells)
% of Subclustered T Cells
αCTLA-4
αCTLA-4+αPD-1
Cd8Cycling vs Others
Hallmark_Myc_Target_V1
Hallmark_Oxydative_Phosphorylation
Hallmark_E2F_Target
Hallmark_DNA_Repair
Hallmark_MTORC1_Signalling
Hallmark_Fatty_acid_Metabolism
Hallmark_G2M_Checkpoint
Hallmark_Myc_Targets_V2
Hallmark_Glycolysis
Hallmark_Adipogenesis
Hallmark_Cholesterol_Homeostatisis
Hallmark_Spermatogenesis
Hallmark_Xenobiotic_Metabolism
Hallmark_Peroxisome
Hallmark_TNFA_Signalling_Via_NFKB
Hallmark_Interferon_Alpha_Response
Hallmark_Interferon_Gamma_Response
Hallmark_Inflammatory_Response
Hallmark_IL-6_JAK_STAT3_Response
Hallmark_IL-2_STAT5_Response
Hallmark_UV_Response_DN
Hallmark_TGF_Beta_Signaling
Hallmark_KRAS_Signaling_UP
Hallmark_Estrogen_Response_Early
Hallmark_Apoptosis
Hallmark_Mitotic_Spindle
Hallmark_WNT_Beta_Catenin_Signaling
Hallmark_Allograft_Rejection
Hallmark_Heme_Metabolism
Enrichment Score
C
% Expressed
Average Expression
Cd8Eff/Ex (Subclustered T cells)
Cd8Eff/Ex vs Others
Hallmark_Hypoxia
Hallmark_Complement
Enrichment Score
D
Cd8PE (Subclustered T cells)
Cd8PE vs Others
Hallmark_Unfolded_Protein_Response
Hallmark_IL-2_STAT5_Response
Hallmark_Hypoxia
Hallmark_Glycolysis
Hallmark_Apoptosis
Enrichment Score
E
Cd8iSTIM (Subclustered T cells)
Cd8iSTIM vs Others
Hallmark_Interferon_Alpha_Response
Hallmark_Interferon_Gamma_Response
Hallmark_Myc_Targets_V1
Hallmark_Myogenesis
Hallmark_UV_Response_DN
Hallmark_Protein_Secretion
Hallmark_Adipogenesis
Hallmark_Mitotic_Spindle
Hallmark_PI3K_AKT_MTOR_Signaling
Hallmark_MTORC1_Signaling
Hallmark_Oxidative_Phosphorylation
Hallmark_Fatty_Acid_Metabolism
Hallmark_Spermatogenesis
F
Cd8Ccr7 (Subclustered T cells)
Cd8Ccr7 vs Others
Hallmark_Myc_Targets_V1
Hallmark_Myc_Targets_V2
Hallmark_Oxydative Phosphorylation
Hallmark_Unfolded_Protein_Response
Hallmark_MTORC1_signaling
Hallmark_E2F_Target
Hallmark_Adipogenesis
Hallmark_Mitotic_Spindle
Hallmark_TNFA_Signalling_Via_NFKB
Hallmark_Apical_Junction
Hallmark_Complement
Hallmark_KRAS_Signaling_DN
Hallmark_TGFB_Signaling
Hallmark_Heme_Metabolism
Hallmark_Apoptosis
Hallmark_KRAS_Signaling_UP
Hallmark_Inflammatory_Response
Hallmark_Protein_Secretion
Hallmark_Reactive_Oxygen_Specied_Pathway

**Figure S6. scRNAseq analysis of bulk CD8 T cells from Y1.7LI tumor bearing mice treated with NeoAg vaccines or ICT. Related to Figure 2.**
(A) Heat map displaying normalized expression of select genes in each bulk CD8 T cell clusters (see also Figures 2A and 2D). (B-F) scRNAseq dot plot depicting expression level/percent of cells expressing select transcripts, bar graphs depicting frequency of each CD8 T cell cluster by treatment condition, and GSEA displaying significantly enriched gene sets in each CD8 T cell cluster.

### Figure S7

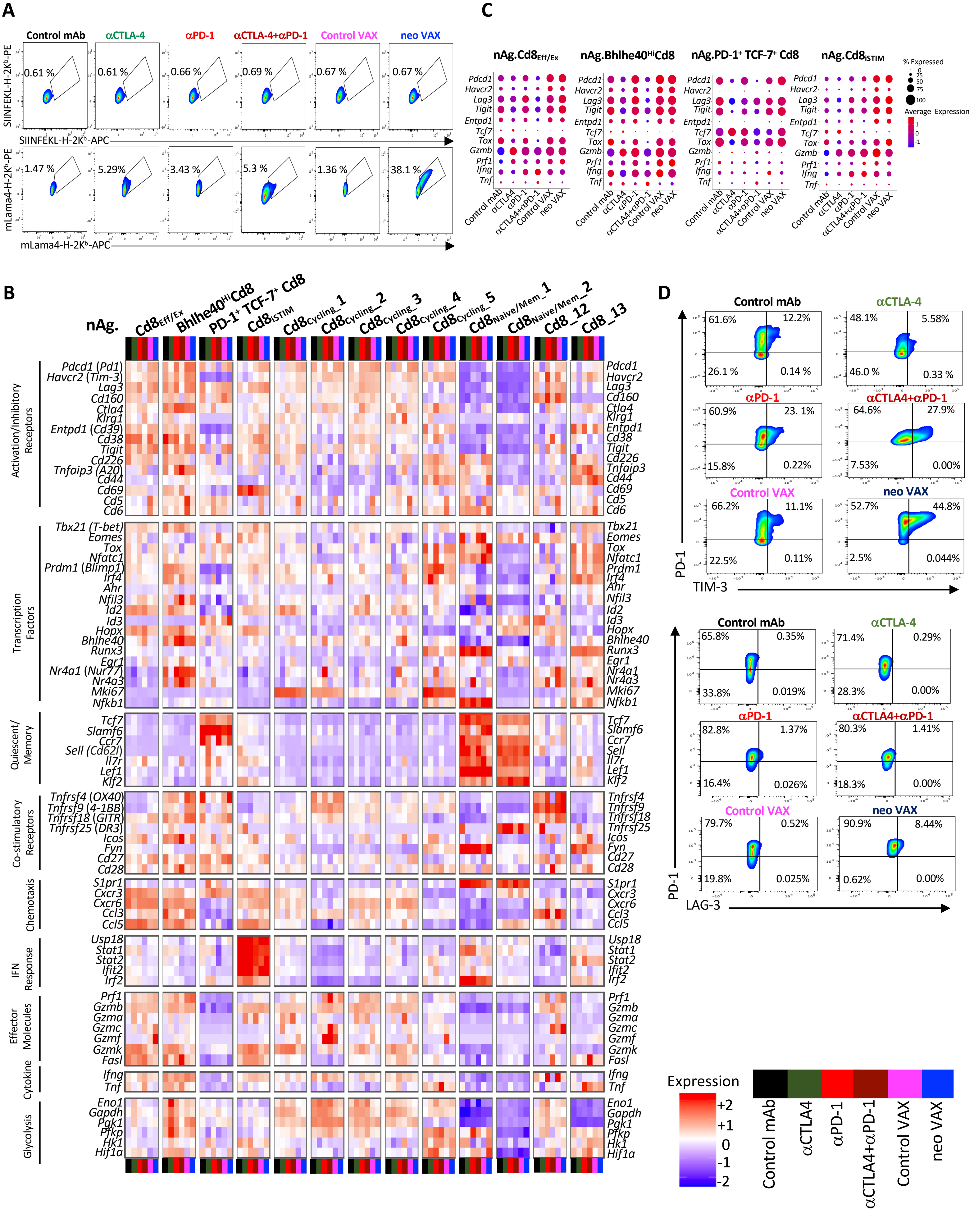
A
Control mAb
αCTLA-4
αPD-1
αCTLA-4+αPD-1
Control VAX
neo VAX
SIINFEKL-H-2Kᵇ-PE
SIINFEKL-H-2Kᵇ-APC
0.61 %
0.61 %
0.66 %
0.69 %
0.67 %
0.67 %
mLama4-H-2Kᵇ-PE
mLama4-H-2Kᵇ-APC
1.47 %
5.29%
3.43 %
5.3 %
1.36 %
38.1 %
C
nAg.Cd8Eff/Ex
nAg.Bhlhe40HiCd8
nAg.PD-1⁺ TCF-7⁺ Cd8
nAg.Cd8iSTIM
Pdcd1
Havcr2
Lag3
Tigit
Entpd1
Tcf7
Tox
Gzmb
Prf1
Ifng
Tnf
Control mAb
αCTLA4
αPD-1
αCTLA4+αPD-1
Control VAX
neo VAX
% Expressed
0
25
50
75
100
Average Expression
1
0
-1
B
nAg.
Cd8Eff/Ex
Bhlhe40HiCd8
PD-1⁺ TCF-7⁺ Cd8
Cd8iSTIM
Cd8cycling_1
Cd8cycling_2
Cd8cycling_3
Cd8cycling_4
Cd8cycling_5
Cd8Naive/Mem_1
Cd8Naive/Mem_2
Cd8_12
Cd8_13
Activation/Inhibitory Receptors
Pdcd1 (Pd1)
Havcr2 (Tim-3)
Lag3
Cd160
Ctla4
Klrg1
Entpd1 (Cd39)
Cd38
Tigit
Cd226
Tnfaip3 (A20)
Cd44
Cd69
Cd5
Cd6
Transcription Factors
Tbx21 (T-bet)
Eomes
Tox
Nfatc1
Prdm1 (Blimp1)
Irf4
Ahr
Nfil3
Id2
Id3
Hopx
Bhlhe40
Runx3
Egr1
Nr4a1 (Nur77)
Nr4a3
Mki67
Nfkb1
Quiescent/ Memory
Tcf7
Slamf6
Ccr7
Sell (Cd62l)
Il7r
Lef1
Klf2
Co-stimulatory Receptors
Tnfrsf4 (OX40)
Tnfrsf9 (4-1BB)
Tnfrsf18 (GITR)
Tnfrsf25 (DR3)
Icos
Fyn
Cd27
Cd28
Chemotaxis
S1pr1
Cxcr3
Cxcr6
Ccl3
Ccl5
IFN Response
Usp18
Stat1
Stat2
Ifit2
Irf2
Effector Molecules
Prf1
Gzmb
Gzma
Gzmc
Gzmf
Gzmk
Fasl
Cytokine
Ifng
Tnf
Glycolysis
Eno1
Gapdh
Pgk1
Pfkp
Hk1
Hif1a
Expression
+2
+1
0
-1
-2
Control mAb
αCTLA4
αPD-1
αCTLA4+αPD-1
Control VAX
neo VAX
D
Control mAb
61.6%
12.2%
26.1 %
0.14 %
αCTLA-4
48.1%
5.58%
46.0 %
0.33 %
αPD-1
60.9%
23. 1%
15.8%
0.22%
αCTLA4+αPD-1
64.6%
27.9%
7.53%
0.00%
Control VAX
66.2%
11.1%
22.5%
0.11%
neo VAX
52.7%
44.8%
2.5%
0.044%
PD-1
TIM-3
Control mAb
65.8%
0.35%
33.8%
0.019%
αCTLA-4
71.4%
0.29%
28.3%
0.00%
αPD-1
82.8%
1.37%
16.4%
0.026%
αCTLA4+αPD-1
80.3%
1.41%
18.3%
0.00%
Control VAX
79.7%
0.52%
19.8%
0.025%
neo VAX
90.9%
8.44%
0.62%
0.00%
PD-1
LAG-3

**Figure S7. scRNAseq and flow cytometry profiling of mLama4 NeoAg-specific CD8 T cells from Y1.7LI tumor bearing mice treated with NeoAg vaccines or ICT. Related to Figure 3.**
(A) Representative flow cytometry plots displaying mLama4 tetramer-specific CD8 T cells in Y1.7LI tumors treated with control mAb, anti-CTLA-4, anti-PD-1, anti-CTLA-4 + anti-PD-1, irrelevant (for Y1.7LI) mAlg8 SLP + pI:C (Control VAX), or relevant mLama4 SLP + pI:C (neo VAX) beginning on d. 7 and harvested on d. 15 post-tumor transplant. mLama4-H2-$K^b$ or SIINFEKL-H2-$K^b$ (irrelevant control) tetramers were labeled with PE and APC. Dot plots are gated on live $CD45^+$ $Thy1.2^+$ CD8 T cells (See also Figure S15). (B) Heat map displaying normalized expression of select genes in each mLama4 NeoAg-specific CD8 T cell clusters by treatment condition (see also Figure 3E). (C) scRNAseq dot plot depicting expression level/percent of cells expressing select transcripts within select mLama4 NeoAg-specific CD8 T cell clusters by treatment condition. (D) Representative flow cytometry plots displaying PD-$1^+$ and/or TIM-$3^+$/LAG-$3^+$ after gating on mLama4 tetramer positive CD8 T cells (See also Figure S15).

**Figure S8**

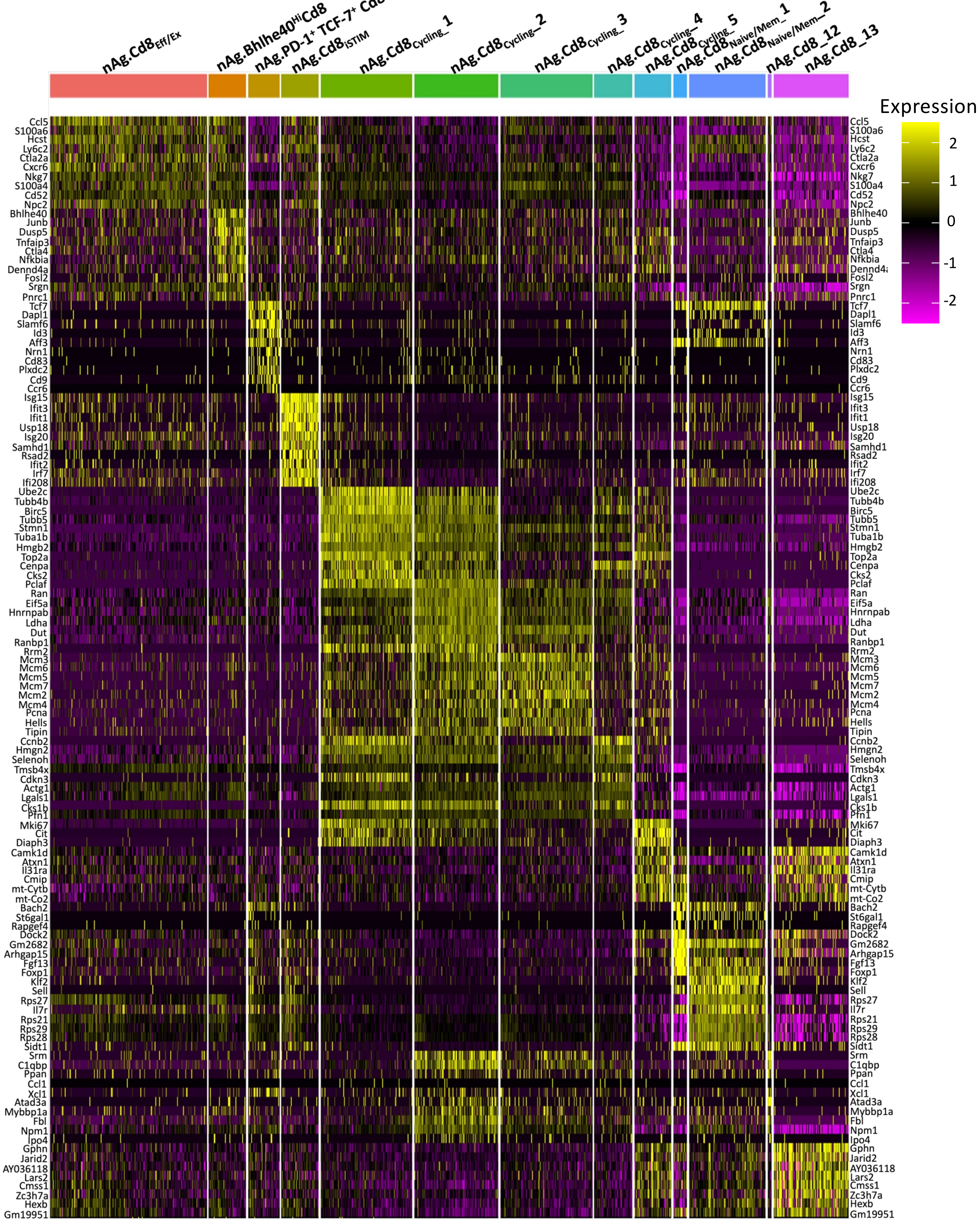
nAg.Cd8Eff/Ex
nAg.Bhlhe40HiCd8
nAg.PD-1+ TCF-7+ Cd8
nAg.Cd8iSTIM
nAg.Cd8Cycling_1
nAg.Cd8Cycling_2
nAg.Cd8Cycling_3
nAg.Cd8Cycling_4
nAg.Cd8Cycling_5
nAg.Cd8Naive/Mem_1
nAg.Cd8Naive/Mem_2
nAg.Cd8_12
nAg.Cd8_13
Expression
2
1
0
-1
-2
Ccl5
S100a6
Hcst
Ly6c2
Ctla2a
Cxcr6
Nkg7
S100a4
Cd52
Npc2
Bhlhe40
Junb
Dusp5
Tnfaip3
Ctla4
Nfkbia
Dennd4a
Fosl2
Srgn
Pnrc1
Tcf7
Dapl1
Slamf6
Id3
Aff3
Nrn1
Cd83
Plxdc2
Cd9
Ccr6
Isg15
Ifit3
Ifit1
Usp18
Isg20
Samhd1
Rsad2
Ifit2
Irf7
Ifi208
Ube2c
Tubb4b
Birc5
Tubb5
Stmn1
Tuba1b
Hmgb2
Top2a
Cenpa
Cks2
Pclaf
Ran
Eif5a
Hnrnpab
Ldha
Dut
Ranbp1
Rrm2
Mcm3
Mcm6
Mcm5
Mcm7
Mcm2
Mcm4
Pcna
Hells
Tipin
Ccnb2
Hmgn2
Selenoh
Tmsb4x
Cdkn3
Actg1
Lgals1
Cks1b
Pfn1
Mki67
Cit
Diaph3
Camk1d
Atxn1
Il31ra
Cmip
mt-Cytb
mt-Co2
Bach2
St6gal1
Rapgef4
Dock2
Gm2682
Arhgap15
Fgf13
Foxp1
Klf2
Sell
Rps27
Il7r
Rps21
Rps29
Rps28
Sidt1
Srm
C1qbp
Ppan
Ccl1
Xcl1
Atad3a
Mybbp1a
Fbl
Npm1
Ipo4
Gphn
Jarid2
AY036118
Lars2
Cmss1
Zc3h7a
Hexb
Gm19951

**Figure S8. Heat map of the top 10 most DEG across NeoAg-specific CD8 T cells clusters. Related to Figure 3.**

### Figure S9

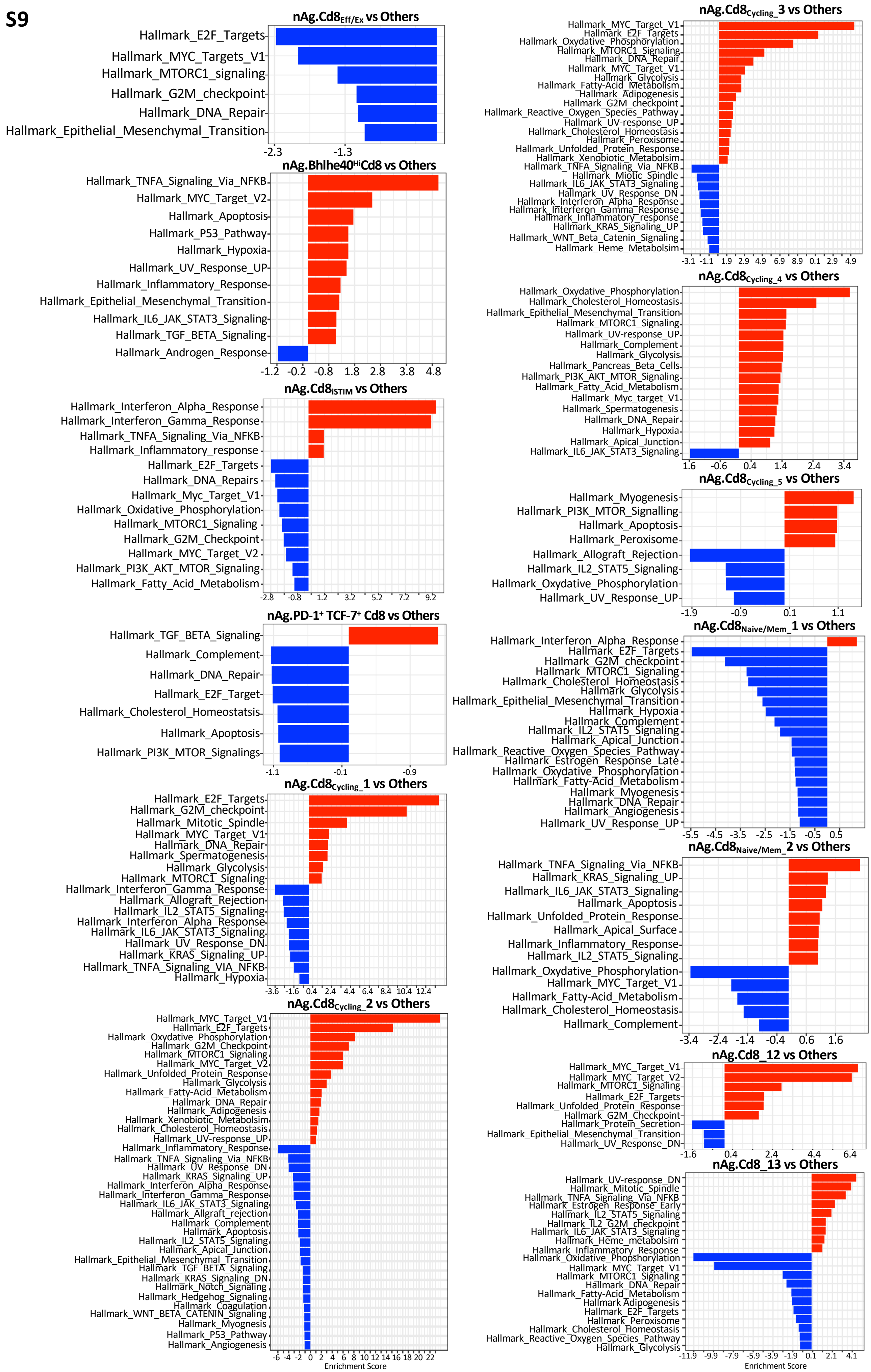

nAg.Cd8Eff/Ex vs Others
Hallmark_E2F_Targets
Hallmark_MYC_Targets_V1
Hallmark_MTORC1_signaling
Hallmark_G2M_checkpoint
Hallmark_DNA_Repair
Hallmark_Epithelial_Mesenchymal_Transition
-2.3
-1.3
nAg.Bhlhe40HiCd8 vs Others
Hallmark_TNFA_Signaling_Via_NFKB
Hallmark_MYC_Target_V2
Hallmark_Apoptosis
Hallmark_P53_Pathway
Hallmark_Hypoxia
Hallmark_UV_Response_UP
Hallmark_Inflammatory_Response
Hallmark_Epithelial_Mesenchymal_Transition
Hallmark_IL6_JAK_STAT3_Signaling
Hallmark_TGF_BETA_Signaling
Hallmark_Androgen_Response
-1.2
-0.2
-0.8
1.8
2.8
3.8
4.8
nAg.Cd8ISTIM vs Others
Hallmark_Interferon_Alpha_Response
Hallmark_Interferon_Gamma_Response
Hallmark_TNFA_Signaling_Via_NFKB
Hallmark_Inflammatory_response
Hallmark_E2F_Targets
Hallmark_DNA_Repairs
Hallmark_Myc_Target_V1
Hallmark_Oxidative_Phosphorylation
Hallmark_MTORC1_Signaling
Hallmark_G2M_Checkpoint
Hallmark_MYC_Target_V2
Hallmark_PI3K_AKT_MTOR_Signaling
Hallmark_Fatty_Acid_Metabolism
-2.8
-0.8
1.2
3.2
5.2
7.2
9.2
nAg.PD-1+ TCF-7+ Cd8 vs Others
Hallmark_TGF_BETA_Signaling
Hallmark_Complement
Hallmark_DNA_Repair
Hallmark_E2F_Target
Hallmark_Cholesterol_Homeostatsis
Hallmark_Apoptosis
Hallmark_PI3K_MTOR_Signalings
-1.1
-0.1
-0.9
nAg.Cd8Cycling_1 vs Others
Hallmark_E2F_Targets
Hallmark_G2M_checkpoint
Hallmark_Mitotic_Spindle
Hallmark_MYC_Target_V1
Hallmark_DNA_Repair
Hallmark_Spermatogenesis
Hallmark_Glycolysis
Hallmark_MTORC1_Signaling
Hallmark_Interferon_Gamma_Response
Hallmark_Allograft_Rejection
Hallmark_IL2_STAT5_Signaling
Hallmark_Interferon_Alpha_Response
Hallmark_IL6_JAK_STAT3_Signaling
Hallmark_UV_Response_DN
Hallmark_KRAS_Signaling_UP
Hallmark_TNFA_Signaling_VIA_NFKB
Hallmark_Hypoxia
-3.6
-1.6
0.4
2.4
4.4
6.4
8.4
10.4
12.4
nAg.Cd8Cycling_2 vs Others
Hallmark_MYC_Target_V1
Hallmark_E2F_Targets
Hallmark_Oxydative_Phosphorylation
Hallmark_G2M_Checkpoint
Hallmark_MTORC1_Signaling
Hallmark_MYC_Target_V2
Hallmark_Unfolded_Protein_Response
Hallmark_Glycolysis
Hallmark_Fatty-Acid_Metabolism
Hallmark_DNA_Repair
Hallmark_Adipogenesis
Hallmark_Xenobiotic_Metabolsim
Hallmark_Cholesterol_Homeostasis
Hallmark_UV-response_UP
Hallmark_Inflammatory_Response
Hallmark_TNFA_Signaling_Via_NFKB
Hallmark_UV_Response_DN
Hallmark_KRAS_Signaling_UP
Hallmark_Interferon_Alpha_Response
Hallmark_Interferon_Gamma_Response
Hallmark_IL6_JAK_STAT3_Signaling
Hallmark_Allgraft_rejection
Hallmark_Complement
Hallmark_Apoptosis
Hallmark_IL2_STAT5_Signaling
Hallmark_Apical_Junction
Hallmark_Epithelial_Mesenchymal_Transition
Hallmark_TGF_BETA_Signaling
Hallmark_KRAS_Signaling_DN
Hallmark_Notch_Signaling
Hallmark_Hedgehog_Signaling
Hallmark_Coagulation
Hallmark_WNT_BETA_CATENIN_Signaling
Hallmark_Myognesis
Hallmark_P53_Pathway
Hallmark_Angiogenesis
-6
-4
-2
0
2
4
6
8
10
12
14
16
18
20
22
Enrichment Score
nAg.Cd8Cycling_3 vs Others
Hallmark_MYC_Target_V1
Hallmark_E2F_Targets
Hallmark_Oxydative_Phosphorylation
Hallmark_MTORC1_Signaling
Hallmark_DNA_Repair
Hallmark_MYC_Target_V1
Hallmark_Glycolysis
Hallmark_Fatty-Acid_Metabolism
Hallmark_Adipogenesis
Hallmark_G2M_checkpoint
Hallmark_Reactive_Oxygen_Species_Pathway
Hallmark_UV-response_UP
Hallmark_Cholesterol_Homeostasis
Hallmark_Peroxisome
Hallmark_Unfolded_Protein_Response
Hallmark_Xenobiotic_Metabolsim
Hallmark_TNFA_Signaling_Via_NFKB
Hallmark_Miotic_Spindle
Hallmark_IL6_JAK_STAT3_Signaling
Hallmark_UV_Response_DN
Hallmark_Interferon_Alpha_Response
Hallmark_Interferon_Gamma_Response
Hallmark_Inflammatory_response
Hallmark_KRAS_Signaling_UP
Hallmark_WNT_Beta_Catenin_Signaling
Hallmark_Heme_Metabolsim
-3.1
-1.1
1.9
2.9
4.9
6.9
8.9
0.1
2.9
4.9
nAg.Cd8Cycling_4 vs Others
Hallmark_Oxydative_Phosphorylation
Hallmark_Cholesterol_Homeostasis
Hallmark_Epithelial_Mesenchymal_Transition
Hallmark_MTORC1_Signaling
Hallmark_UV-response_UP
Hallmark_Complement
Hallmark_Glycolysis
Hallmark_Pancreas_Beta_Cells
Hallmark_PI3K_AKT_MTOR_Signaling
Hallmark_Fatty_Acid_Metabolism
Hallmark_Myc_target_V1
Hallmark_Spermatogenesis
Hallmark_DNA_Repair
Hallmark_Hypoxia
Hallmark_Apical_Junction
Hallmark_IL6_JAK_STAT3_Signaling
1.6
-0.6
0.4
1.4
2.4
3.4
nAg.Cd8Cycling_5 vs Others
Hallmark_Myogenesis
Hallmark_PI3K_MTOR_Signalling
Hallmark_Apoptosis
Hallmark_Peroxisome
Hallmark_Allograft_Rejection
Hallmark_IL2_STAT5_Signaling
Hallmark_Oxydative_Phosphorylation
Hallmark_UV_Response_UP
-1.9
-0.9
0.1
1.1
nAg.Cd8Naive/Mem_1 vs Others
Hallmark_Interferon_Alpha_Response
Hallmark_E2F_Targets
Hallmark_G2M_checkpoint
Hallmark_MTORC1_Signaling
Hallmark_Cholesterol_Homeostasis
Hallmark_Glycolysis
Hallmark_Epithelial_Mesenchymal_Transition
Hallmark_Hypoxia
Hallmark_Complement
Hallmark_IL2_STAT5_Signaling
Hallmark_Apical_Junction
Hallmark_Reactive_Oxygen_Species_Pathway
Hallmark_Estrogen_Response_Late
Hallmark_Oxydative_Phosphorylation
Hallmark_Fatty-Acid_Metabolism
Hallmark_Myogenesis
Hallmark_DNA_Repair
Hallmark_Angiogenesis
Hallmark_UV_Response_UP
-5.5
-4.5
-3.5
-2.5
-1.5
-0.5
0.5
nAg.Cd8Naive/Mem_2 vs Others
Hallmark_TNFA_Signaling_Via_NFKB
Hallmark_KRAS_Signaling_UP
Hallmark_IL6_JAK_STAT3_Signaling
Hallmark_Apoptosis
Hallmark_Unfolded_Protein_Response
Hallmark_Apical_Surface
Hallmark_Inflammatory_Response
Hallmark_IL2_STAT5_Signaling
Hallmark_Oxydative_Phosphorylation
Hallmark_MYC_Target_V1
Hallmark_Fatty-Acid_Metabolism
Hallmark_Cholesterol_Homeostasis
Hallmark_Complement
-3.4
-2.4
-1.4
-0.4
0.6
1.6
nAg.Cd8_12 vs Others
Hallmark_MYC_Target_V1
Hallmark_MYC_Target_V2
Hallmark_MTORC1_Signaling
Hallmark_E2F_Targets
Hallmark_Unfolded_Protein_Response
Hallmark_G2M_Checkpoint
Hallmark_Protein_Secretion
Hallmark_Epithelial_Mesenchymal_Transition
Hallmark_UV_Response_DN
-1.6
0.4
2.4
4.4
6.4
nAg.Cd8_13 vs Others
Hallmark_UV-response_DN
Hallmark_Mitotic_Spindle
Hallmark_TNFA_Signaling_Via_NFKB
Hallmark_Estrogen_Response_Early
Hallmark_IL2_STAT5_Signaling
Hallmark_IL2_G2M_checkpoint
Hallmark_IL6_JAK_STAT3_Signaling
Hallmark_Heme_metabolsim
Hallmark_Inflammatory_Response
Hallmark_Oxidative_Phopshorylation
Hallmark_MYC_Target_V1
Hallmark_MTORC1_Signaling
Hallmark_DNA_Repair
Hallmark_Fatty-Acid_Metabolism
Hallmark Adipogenesis
Hallmark_E2F_Targets
Hallmark_Peroxisome
Hallmark_Cholesterol_Homeostasis
Hallmark_Reactive_Oxygen_Species_Pathway
Hallmark_Glycolysis
-11.9
-9.9
-7.9
-5.9
-3.9
-1.9
0.1
2.1
4.1
Enrichment Score

**Figure S9. GSEA displaying significantly enriched gene sets for each NeoAg-specific CD8 T cells cluster. Related to Figure 3.**

#### Figure S10

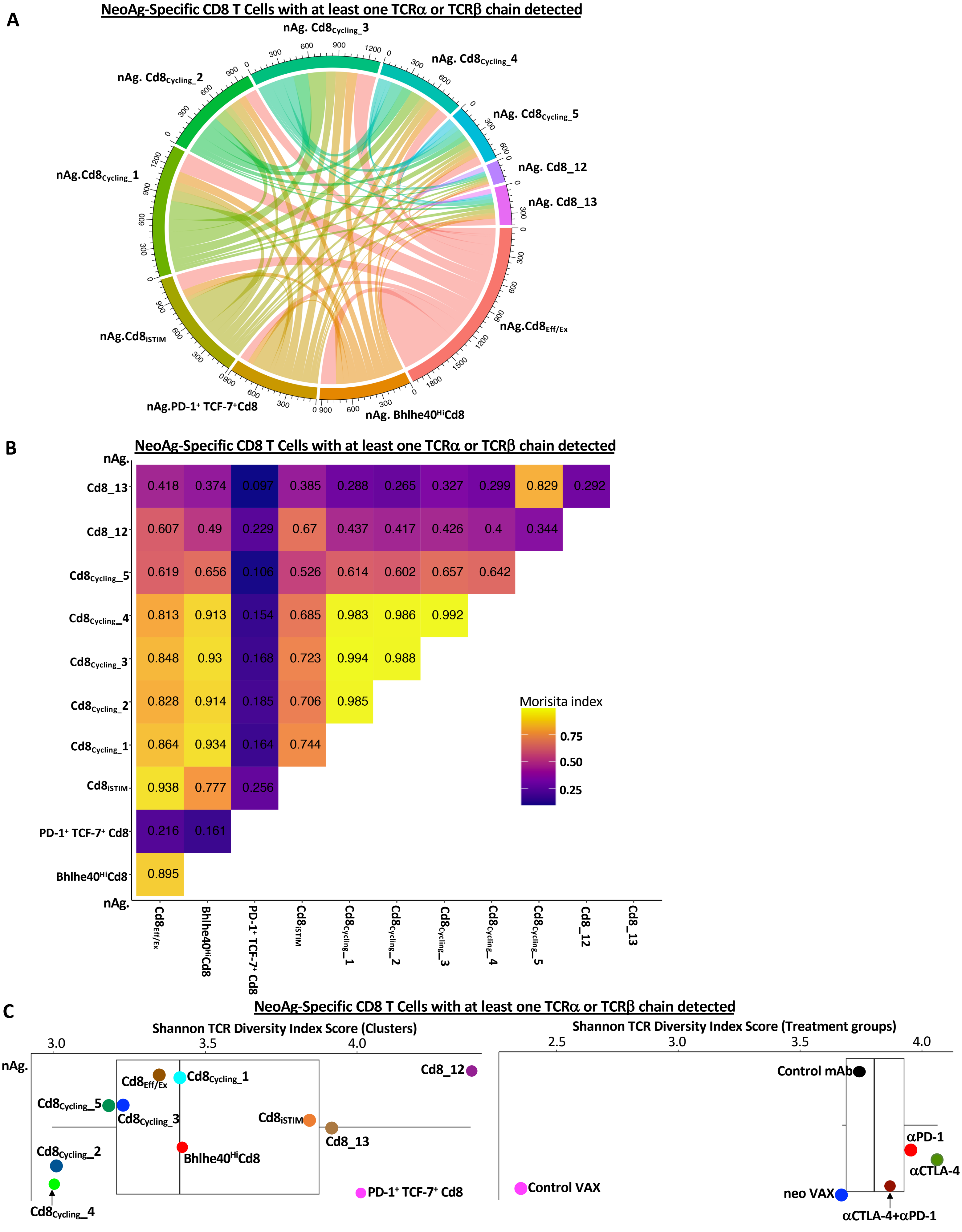
A
NeoAg-Specific CD8 T Cells with at least one TCRα or TCRβ chain detected
nAg. Cd8Cycling_3
nAg. Cd8Cycling_4
nAg. Cd8Cycling_5
nAg. Cd8_12
nAg. Cd8_13
nAg.Cd8Eff/Ex
nAg. Bhlhe40HiCd8
nAg.PD-1+ TCF-7+Cd8
nAg.Cd8iSTIM
nAg.Cd8Cycling_1
nAg. Cd8Cycling_2
B
NeoAg-Specific CD8 T Cells with at least one TCRα or TCRβ chain detected
nAg.
Cd8_13: 0.418 0.374 0.097 0.385 0.288 0.265 0.327 0.299 0.829 0.292
Cd8_12: 0.607 0.49 0.229 0.67 0.437 0.417 0.426 0.4 0.344
Cd8Cycling_5: 0.619 0.656 0.106 0.526 0.614 0.602 0.657 0.642
Cd8Cycling_4: 0.813 0.913 0.154 0.685 0.983 0.986 0.992
Cd8Cycling_3: 0.848 0.93 0.168 0.723 0.994 0.988
Cd8Cycling_2: 0.828 0.914 0.185 0.706 0.985
Cd8Cycling_1: 0.864 0.934 0.164 0.744
Cd8iSTIM: 0.938 0.777 0.256
PD-1+ TCF-7+ Cd8: 0.216 0.161
Bhlhe40HiCd8: 0.895
nAg.
Cd8Eff/Ex
Bhlhe40HiCd8
PD-1+ TCF-7+ Cd8
Cd8iSTIM
Cd8Cycling_1
Cd8Cycling_2
Cd8Cycling_3
Cd8Cycling_4
Cd8Cycling_5
Cd8_12
Cd8_13
Morisita index
0.75
0.50
0.25
C
NeoAg-Specific CD8 T Cells with at least one TCRα or TCRβ chain detected
Shannon TCR Diversity Index Score (Clusters)
3.0
3.5
4.0
nAg.
Cd8Eff/Ex
Cd8Cycling_1
Cd8_12
Cd8Cycling_5
Cd8Cycling_3
Cd8iSTIM
Cd8_13
Cd8Cycling_2
Bhlhe40HiCd8
PD-1+ TCF-7+ Cd8
Cd8Cycling_4
Shannon TCR Diversity Index Score (Treatment groups)
2.5
3.0
3.5
4.0
Control mAb
αPD-1
αCTLA-4
Control VAX
neo VAX
αCTLA-4+αPD-1

**Figure S10. scTCRseq analysis of mLama4 NeoAg-specific CD8 T cells with at least one TCR alpha or beta or both chains. Related to Figure 4.**

(A) Chord diagram displaying overlapping TCR clonotypes (TCR$\alpha$/TCR$\beta$/TCR$\alpha\beta$) of mLama4 NeoAg-specific CD8 T cells by cluster. (B) Morisita index values depicting overlapping TCR clonotypes (TCR$\alpha$/TCR$\beta$/TCR$\alpha\beta$) of mLama4 NeoAg-specific CD8 T cells by cluster. (C) Shannon TCR (TCR$\alpha$/TCR$\beta$/TCR$\alpha\beta$) diversity index of by clusters and treatment groups.

### Figure S11

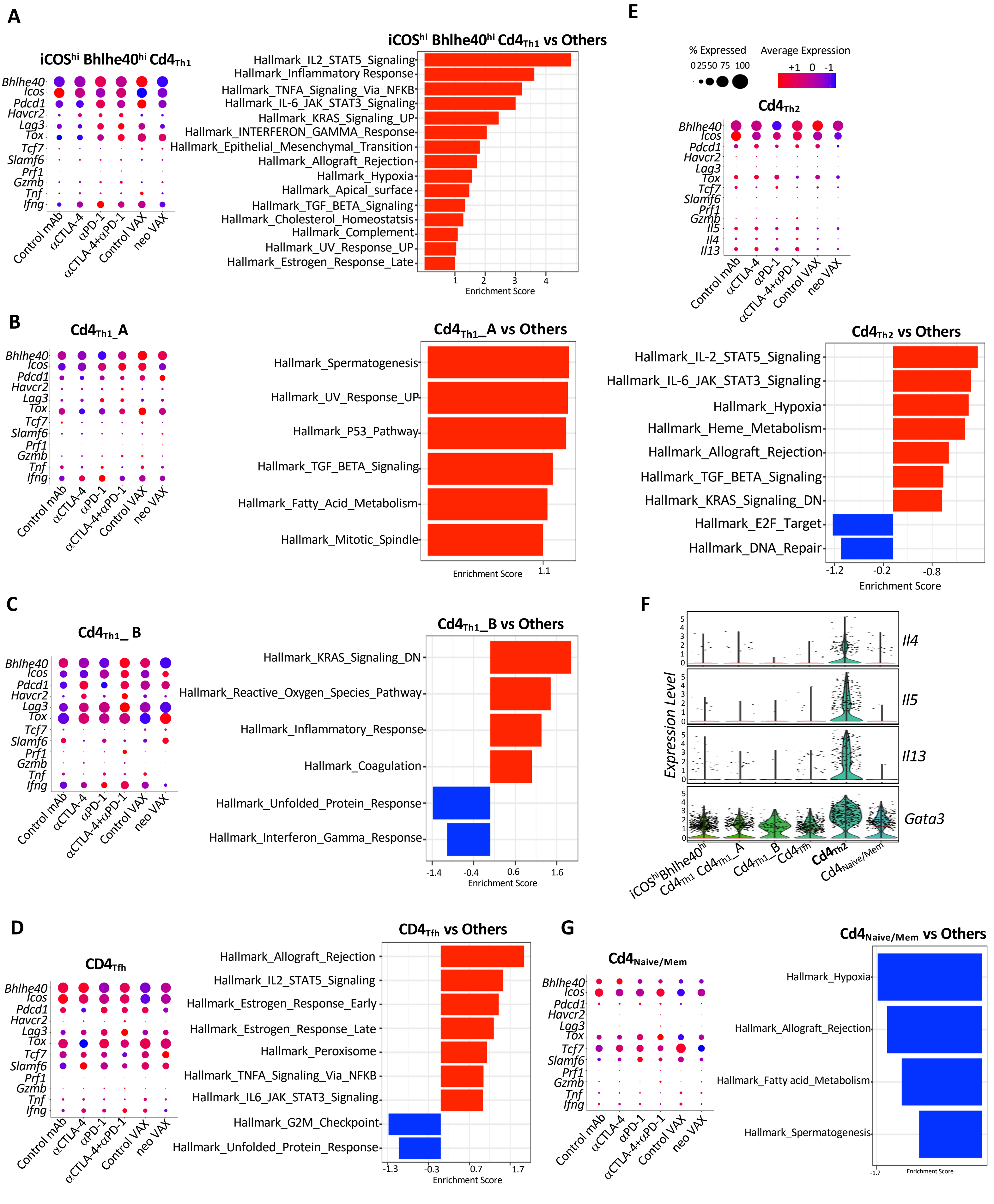

A
iCOS^hi Bhlhe40^hi Cd4Th1
Bhlhe40
Icos
Pdcd1
Havcr2
Lag3
Tox
Tcf7
Slamf6
Prf1
Gzmb
Tnf
Ifng
Control mAb
αCTLA-4
αPD-1
αCTLA-4+αPD-1
Control VAX
neo VAX
iCOS^hi Bhlhe40^hi Cd4Th1 vs Others
Hallmark_IL2_STAT5_Signaling
Hallmark_Inflammatory Response
Hallmark_TNFA_Signaling_Via_NFKB
Hallmark_IL-6_JAK_STAT3_Signaling
Hallmark_KRAS_Signaling_UP
Hallmark_INTERFERON_GAMMA_Response
Hallmark_Epithelial_Mesenchymal_Transition
Hallmark_Allograft_Rejection
Hallmark_Hypoxia
Hallmark_Apical_surface
Hallmark_TGF_BETA_Signaling
Hallmark_Cholesterol_Homeostatsis
Hallmark_Complement
Hallmark_UV_Response_UP
Hallmark_Estrogen_Response_Late
1 2 3 4
Enrichment Score
E
% Expressed
0 25 50 75 100
Average Expression
+1 0 -1
Cd4Th2
Bhlhe40
Icos
Pdcd1
Havcr2
Lag3
Tox
Tcf7
Slamf6
Prf1
Gzmb
Il5
Il4
Il13
B
Cd4Th1_A
Cd4Th1_A vs Others
Hallmark_Spermatogenesis
Hallmark_UV_Response_UP
Hallmark_P53_Pathway
Hallmark_TGF_BETA_Signaling
Hallmark_Fatty_Acid_Metabolism
Hallmark_Mitotic_Spindle
Enrichment Score
1.1
Cd4Th2 vs Others
Hallmark_IL-2_STAT5_Signaling
Hallmark_IL-6_JAK_STAT3_Signaling
Hallmark_Hypoxia
Hallmark_Heme_Metabolism
Hallmark_Allograft_Rejection
Hallmark_TGF_BETA_Signaling
Hallmark_KRAS_Signaling_DN
Hallmark_E2F_Target
Hallmark_DNA_Repair
-1.2 -0.2 -0.8
Enrichment Score
C
Cd4Th1_ B
Cd4Th1_B vs Others
Hallmark_KRAS_Signaling_DN
Hallmark_Reactive_Oxygen_Species_Pathway
Hallmark_Inflammatory_Response
Hallmark_Coagulation
Hallmark_Unfolded_Protein_Response
Hallmark_Interferon_Gamma_Response
-1.4 -0.4 0.6 1.6
Enrichment Score
F
Expression Level
Il4
Il5
Il13
Gata3
iCOS^hi Bhlhe40^hi Cd4Th1
Cd4Th1_A
Cd4Th1_B
Cd4Tfh
Cd4Th2
Cd4Naive/Mem
D
CD4Tfh
CD4Tfh vs Others
Hallmark_Allograft_Rejection
Hallmark_IL2_STAT5_Signaling
Hallmark_Estrogen_Response_Early
Hallmark_Estrogen_Response_Late
Hallmark_Peroxisome
Hallmark_TNFA_Signaling_Via_NFKB
Hallmark_IL6_JAK_STAT3_Signaling
Hallmark_G2M_Checkpoint
Hallmark_Unfolded_Protein_Response
-1.3 -0.3 0.7 1.7
Enrichment Score
G
Cd4Naive/Mem
Cd4Naive/Mem vs Others
Hallmark_Hypoxia
Hallmark_Allograft_Rejection
Hallmark_Fatty acid_Metabolism
Hallmark_Spermatogenesis
-1.7
Enrichment Score

**Figure S11. Select transcript alterations and cluster-specific enriched gene sets in conventional CD4 T cells (see also Figure 2A). Related to Figure 5.**

(A-E and G) scRNAseq dot plots depicting expression level/percent of cells expressing select transcripts and GSEA displaying significantly enriched gene sets within each CD4 T cell cluster by treatment condition (see also Figure 2D). (F) Violin plots denoting expression level of select genes per CD4 T cell.

**Figure S12**

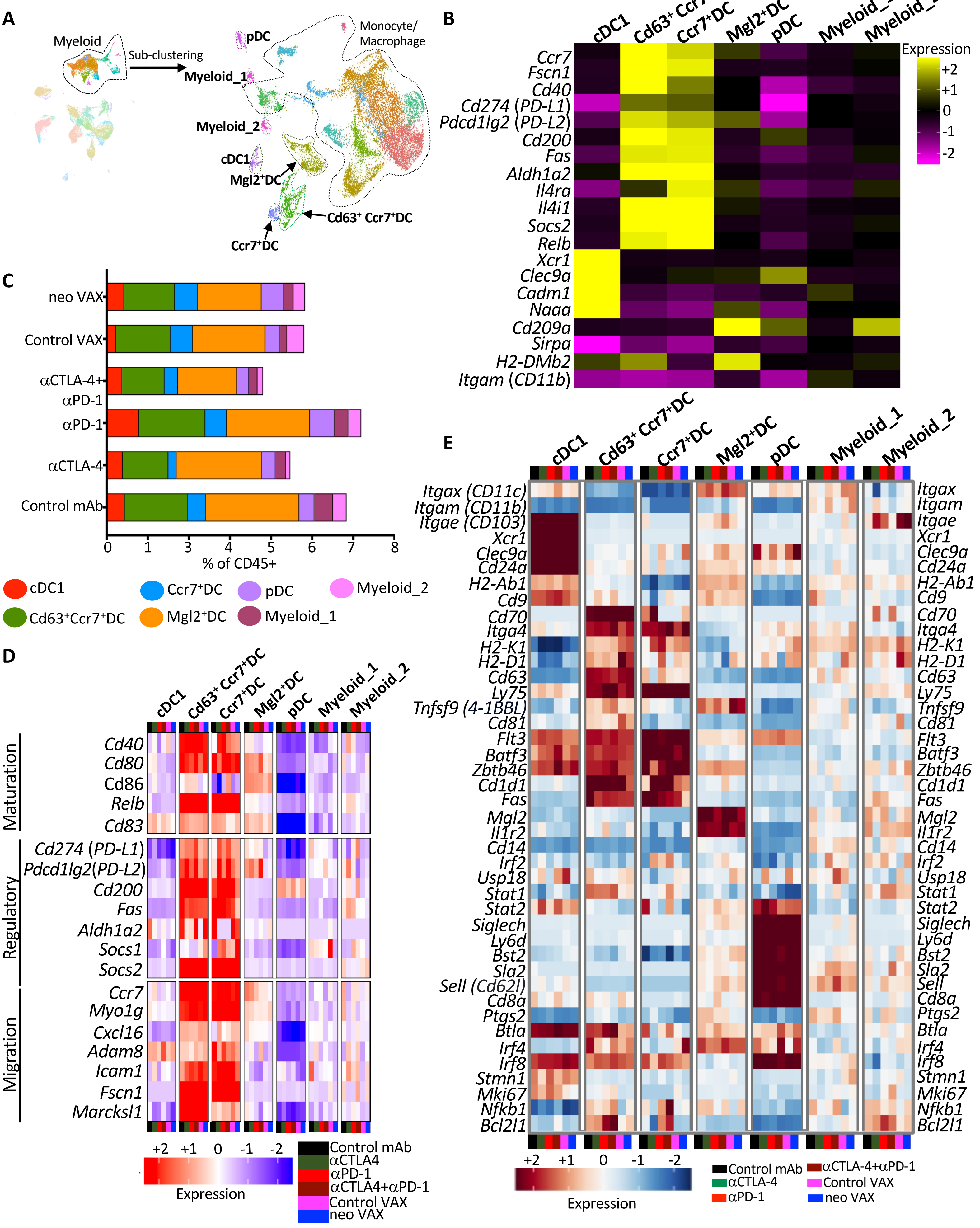
A
Myeloid
Sub-clustering
pDC
Monocyte/
Macrophage
Myeloid_1
Myeloid_2
cDC1
Mgl2+DC
Cd63+ Ccr7+DC
Ccr7+DC
B
cDC1
Cd63+ Ccr7+DC
Ccr7+DC
Mgl2+DC
pDC
Myeloid_1
Myeloid_2
Expression
+2
+1
0
-1
-2
Ccr7
Fscn1
Cd40
Cd274 (PD-L1)
Pdcd1lg2 (PD-L2)
Cd200
Fas
Aldh1a2
Il4ra
Il4i1
Socs2
Relb
Xcr1
Clec9a
Cadm1
Naaa
Cd209a
Sirpa
H2-DMb2
Itgam (CD11b)
C
neo VAX
Control VAX
αCTLA-4+αPD-1
αPD-1
αCTLA-4
Control mAb
0
1
2
3
4
5
6
7
8
% of CD45+
cDC1
Cd63+Ccr7+DC
Ccr7+DC
Mgl2+DC
pDC
Myeloid_1
Myeloid_2
D
cDC1
Cd63+ Ccr7+DC
Ccr7+DC
Mgl2+DC
pDC
Myeloid_1
Myeloid_2
Maturation
Cd40
Cd80
Cd86
Relb
Cd83
Regulatory
Cd274 (PD-L1)
Pdcd1lg2(PD-L2)
Cd200
Fas
Aldh1a2
Socs1
Socs2
Migration
Ccr7
Myo1g
Cxcl16
Adam8
Icam1
Fscn1
Marcksl1
+2
+1
0
-1
-2
Expression
Control mAb
αCTLA4
αPD-1
αCTLA4+αPD-1
Control VAX
neo VAX
E
cDC1
Cd63+ Ccr7+DC
Ccr7+DC
Mgl2+DC
pDC
Myeloid_1
Myeloid_2
Itgax (CD11c)
Itgam (CD11b)
Itgae (CD103)
Xcr1
Clec9a
Cd24a
H2-Ab1
Cd9
Cd70
Itga4
H2-K1
H2-D1
Cd63
Ly75
Tnfsf9 (4-1BBL)
Cd81
Flt3
Batf3
Zbtb46
Cd1d1
Fas
Mgl2
Il1r2
Cd14
Irf2
Usp18
Stat1
Stat2
Siglech
Ly6d
Bst2
Sla2
Sell (Cd62l)
Cd8a
Ptgs2
Btla
Irf4
Irf8
Stmn1
Mki67
Nfkb1
Bcl2l1
+2
+1
0
-1
-2
Expression
Control mAb
αCTLA-4
αPD-1
αCTLA-4+αPD-1
Control VAX
neo VAX

**Figure S12. scRNAseq analysis of dendritic cell (DC) clusters from Y1.7LI tumor bearing mice treated with NeoAg vaccines or ICT.**

(A) UMAP displaying myeloid cell sub-clustering and DC annotations (See also Figures 2A and 6A). (B) Heat map displaying normalized expression of select genes in each DC cluster. (C) Graphs depicting frequency of DCs in each cluster by condition and treatment represented as percent of live CD45$^+$ cells. (D) Heat map displaying normalized expression of select genes[61] in each DC cluster by treatment condition. (E) Heat map displaying normalized expression of select genes in each DC cluster by treatment condition.

**Figure S13**

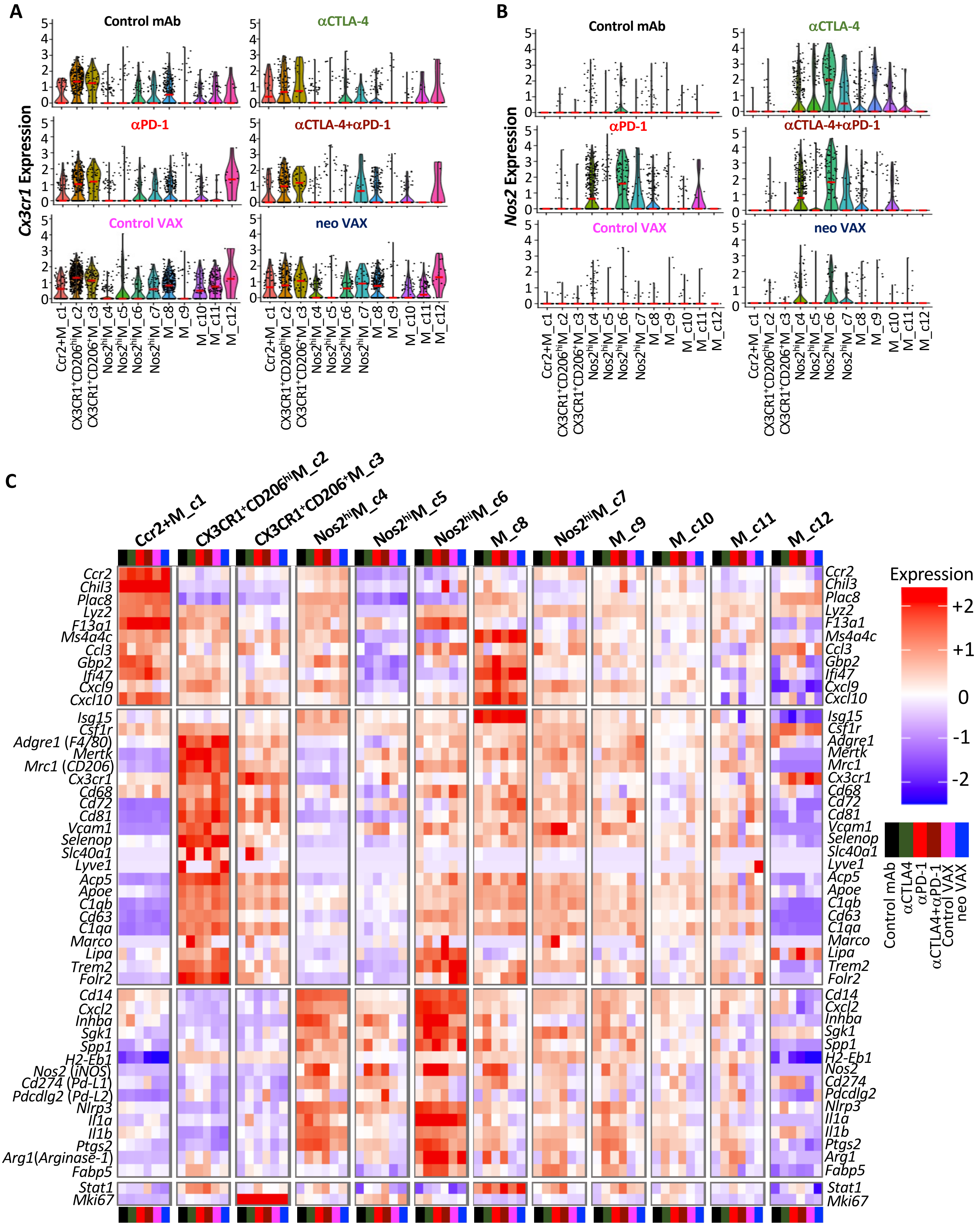
A
Cx3cr1 Expression
Control mAb
αCTLA-4
αPD-1
αCTLA-4+αPD-1
Control VAX
neo VAX
Ccr2+M_c1
CX3CR1+CD206hiM_c2
CX3CR1+CD206+M_c3
Nos2hiM_c4
Nos2hiM_c5
Nos2hiM_c6
Nos2hiM_c7
M_c8
M_c9
M_c10
M_c11
M_c12
B
Nos2 Expression
C
Expression
+2
+1
0
-1
-2
Control mAb
αCTLA4
αPD-1
αCTLA4+αPD-1
Control VAX
neo VAX
Ccr2
Chil3
Plac8
Lyz2
F13a1
Ms4a4c
Ccl3
Gbp2
Ifi47
Cxcl9
Cxcl10
Isg15
Csf1r
Adgre1 (F4/80)
Mertk
Mrc1 (CD206)
Cx3cr1
Cd68
Cd72
Cd81
Vcam1
Selenop
Slc40a1
Lyve1
Acp5
Apoe
C1qb
Cd63
C1qa
Marco
Lipa
Trem2
Folr2
Cd14
Cxcl2
Inhba
Sgk1
Spp1
H2-Eb1
Nos2 (iNOS)
Cd274 (Pd-L1)
Pdcdlg2 (Pd-L2)
Nlrp3
Il1a
Il1b
Ptgs2
Arg1(Arginase-1)
Fabp5
Stat1
Mki67

**Figure S13. scRNAseq analysis of macrophage clusters from Y1.7LI tumor bearing mice treated with NeoAg vaccines or ICT. Related to Figure 6.**
(A) Violin plots denoting expression level of *Cx3cr1* transcript per cell in each monocyte/macrophage cluster by treatment condition. (B) Violin plots denoting expression level of *Nos2* (iNOS) transcript per cell in each monocyte/macrophage cluster by treatment condition. (C) Heat map displaying normalized expression of select genes in each monocyte/macrophage cluster by treatment condition.

### Figure S14

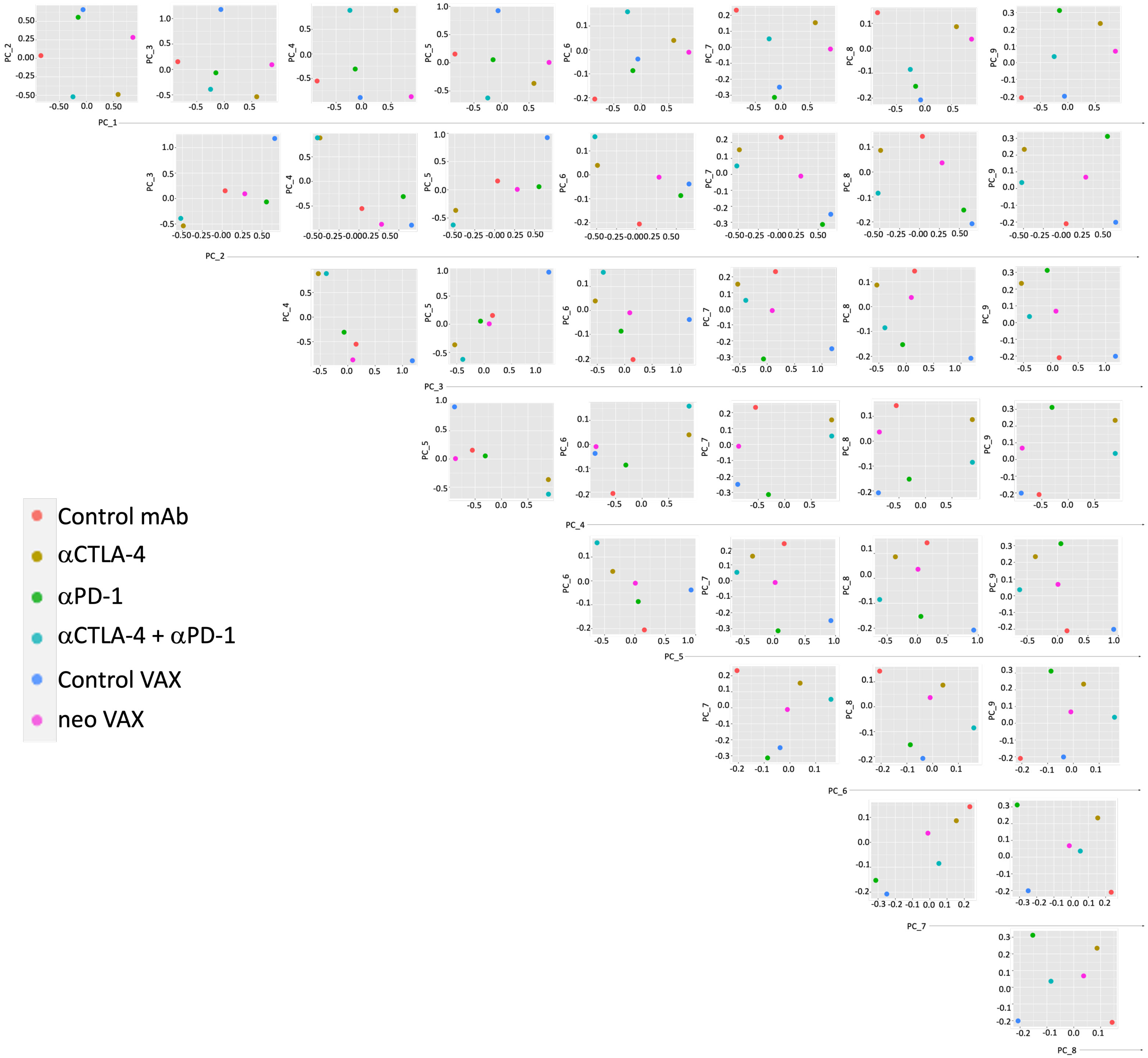
Control mAb
αCTLA-4
αPD-1
αCTLA-4 + αPD-1
Control VAX
neo VAX
PC_1
PC_2
PC_3
PC_4
PC_5
PC_6
PC_7
PC_8
PC_9

**Figure S14. Principal component analysis of subclustered T Cells/ILCs.** Each dot represents individual sample from different treatment conditions (see also Figures 2A and 2D). **Related to Figure 2.**

### Figure S15

#### Myeloid Subsets

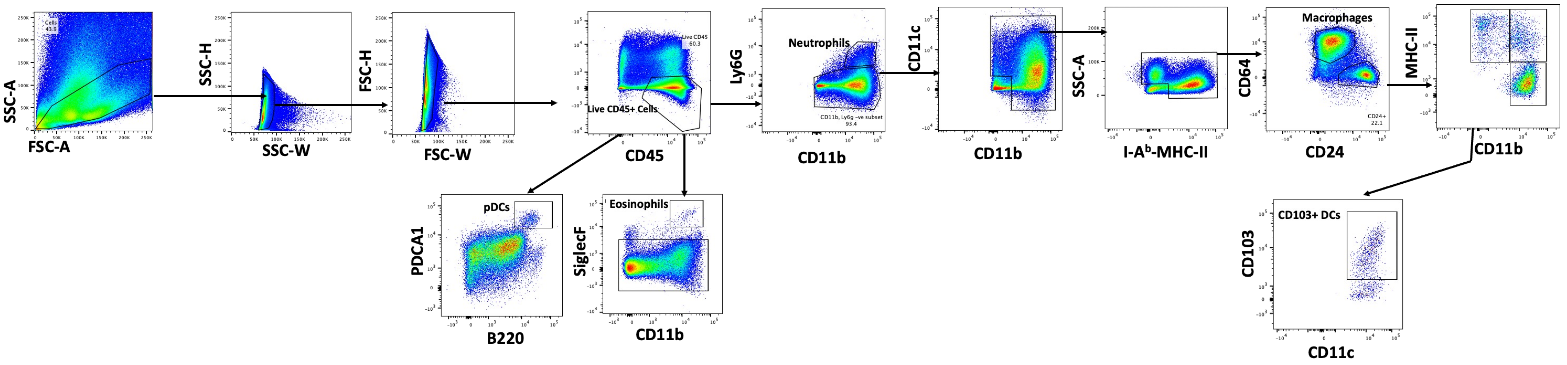


#### Lymphoid Subsets

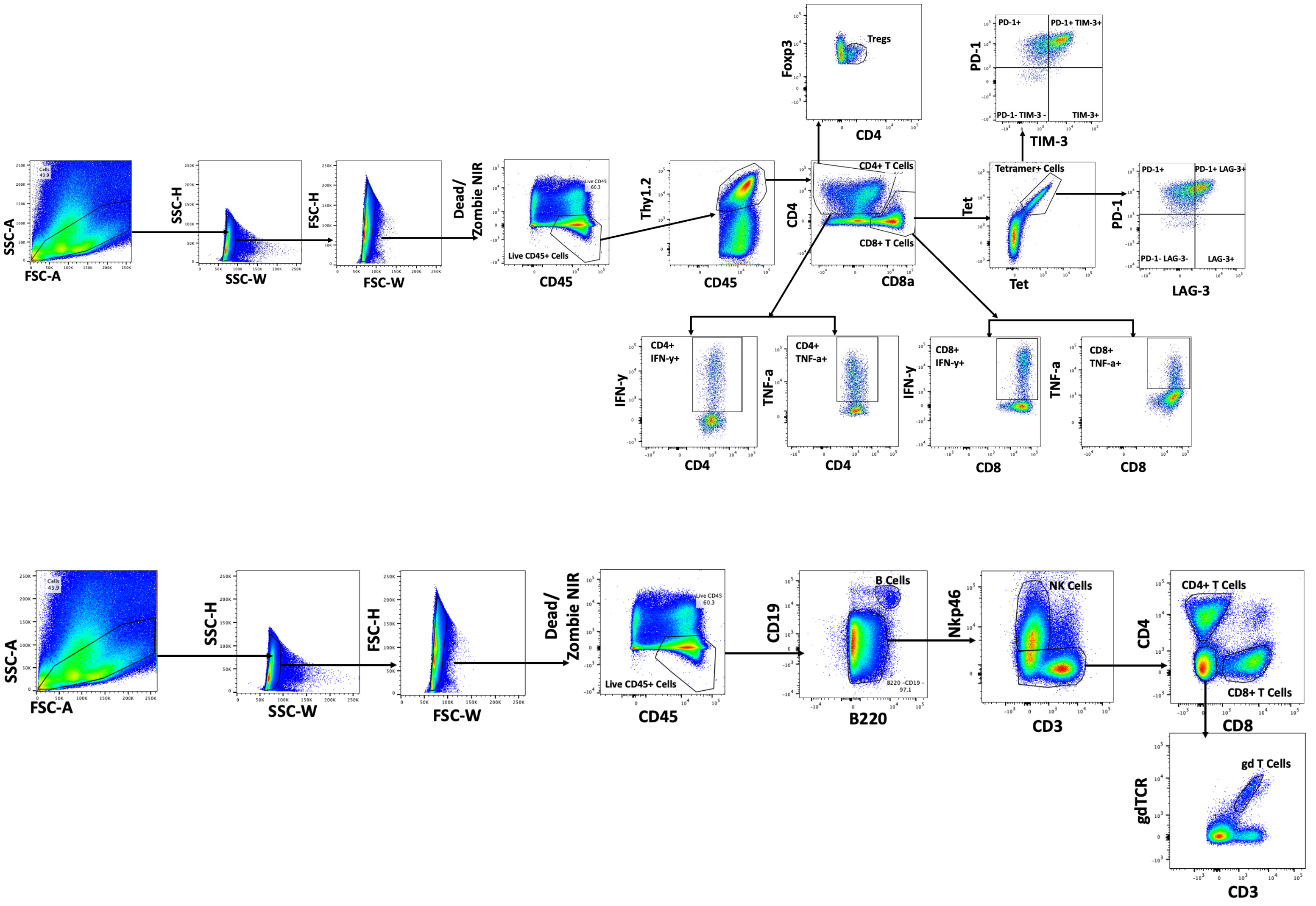

**Figure S15. Gating strategy for identifying intratumoral immune cells.** Flow cytometry dot plots and gating of intratumoral myeloid and lymphoid populations.
