## Supplementary material for "Neoantigen Cancer Vaccines and Different Immune Checkpoint Therapies Each Utilize Both Converging and Distinct Mechanisms that in Combination Enable Synergistic Therapeutic Efficacy": Key Sources Table

| REAGENT or RESOURCE | SOURCE | IDENTIFIER |
|---|---|---|
| ***In vivo* Antibodies** | | |
| Anti-CTLA-4 (clone 9D9) | Leinco | Cat# C2856, RRID: AB_2829611 |
| Anti-CD279 (PD-1) (clone RMP1-14) | Leinco | Cat# RMP1-14, RRID: AB_2749820 |
| Anti-mouse CD8 (clone YTS 169) | Leinco | Cat# C2442, RRID: AB_2829540 |
| Anti-mouse CD4 (Clone GK1.5) | Leinco | Cat# C2838, RRID: AB_2829596 |
| Mouse IgG2a Isotype | Leinco | Cat# P381, RRID: AB_2831654 |
| **Flow Antibodies and reagents** | | |
| Anti-CD16/32 (clone 2.4G2) | BD Biosciences | Cat#553141, RRID: AB_394656 |
| Anti-mouse CD45 BV605 (clone 30-F11) (1:800 dilution) | BioLegend | Cat#103140, RRID: AB_2562342 |
| Anti-mouse CD90.2/Thy1.2-PE-Cy7 (clone 30-H12) (1:200 dilution) | BioLegend | Cat#105326, RRID: AB_2201290 |
| Anti-mouse CD8a-BV786 (clone 53–6.7) (1:200 dilution) | BD Bioscience | Cat#563332, RRID: AB_2721167 |
| Anti-mouse CD4-BV711(clone RM4–5) (1:200 dilution) | BioLegend | Cat#100550, RRID: AB_2562099 |
| Anti-mouse CD19-BV650 (clone 1D3) (1:200 dilution) | BD Bioscience | Cat#563235, RRID: AB_2738085 |
| Anti-mouse CD20-BV421 (clone SA275A11) (1:200 dilution) | BioLegend | Cat#150405, RRID: AB_2566540 |
| Anti-mouse CD45R/B220-BUV395 (clone RA3-6B2) (1:200 dilution) | BD Bioscience | Cat# 563793, RRID: AB_2738427 |
| Anti-mouse Nkp46/CD335-FITC (clone 29A1.4) (1:300 dilution) | BioLegend | Cat# 560756, RRID: AB_1727465 |
| Anti-mouse γδ TCR-PE-Cy7 (clone GL3) (1:300 dilution) | BioLegend | Cat# 118124, RRID: AB_11204423 |
| Anti-mouse PD-1-BV421 (clone 29F.1A12) (1:200 dilution) | BioLegend | Cat# 135218, RRID: AB_2561447 |
| Anti-mouse TIM-3 (clone RMT3-23) (1:200 dilution) | BioLegend | Cat# 119727, RRID: AB_2716208 |
| Anti-mouse LAG-3-PerCP-Cy5.5 (clone C9B7W) (1:200 dilution) | BioLegend | Cat# 125212, RRID: AB_2561517 |
| Anti-mouse CD3e-APC (clone 145–2C11) (1:200 dilution) | BioLegend | Cat# 100312, RRID: AB_312677 |
| Anti-mouse CD64-BV421 (clone X54–5/7.1) (1:200 dilution) | BioLegend | Cat# 139309, RRID: AB_2562694 |
| Anti-mouse Ly6G-Alexa Fluor 700 (clone 1A8) (1:400 dilution) | BD Biosciences | Cat# 127622, RRID: AB_10643269 |
| Anti-mouse CX3CR1-FITC (clone SA011F11) (1:1,000 dilution) | BioLegend | Cat# 149020, RRID: AB_2565703 |

| | | |
|---|---|---|
| Anti-mouse I-A/I-E-BV650 (clone M5/114.15.2) (1:3,000 dilution) | BD Bioscience | Cat# 563415, RRID: AB_2738192 |
| Anti-mouse CD103-BV421 (clone 2E7) (1:200 dilution) | BioLegend | Cat# 121422, RRID: AB_2562901 |
| Anti-mouse CD24-BV711 (clone M1/69) (1:500 dilution) | BD Bioscience | Cat# 563450, RRID: AB_2738213 |
| Anti-mouse CD11c-BV786 (clone HL3) (1:400 dilution) | BD Biosciences | Cat# 563735, RRID: AB_2738394 |
| Anti-mouse CD11b-APC (clone M1/70) (1:400 dilution) | BioLegend | Cat# 101212, RRID: AB_312795 |
| Anti-mouse F4/80-BUV395 (clone T45–2342) (1:400 dilution) | BD Biosciences | Cat# 565614, RRID: AB_2739304 |
| Anti-mouse CD64-APC (clone X54–5/7.1) (1:400 dilution) | BioLegend | Cat# 139306, RRID: AB_11219391 |
| Anti-mouse CD117-FITC (clone ACK2) (1:100 dilution) | BioLegend | Cat# 135115, RRID: AB_2561633 |
| Anti-mouse CD11b- PerCP-Cy5.5 (clone M1/70) (1:200 dilution) | BioLegend | Cat# 561114, RRID: AB_394002 |
| Anti-mouse PDCA- 1/BST-2 BV650 (clone 927) (1:200 dilution) | BD Biosciences | Cat# 747605, RRID: AB_2744173 |
| Anti-mouse CD172a APC (clone P84) (1:200 dilution) | BioLegend | Cat# 144014, RRID: AB_2564061 |
| Anti-mouse CD274/PDL1-PE (clone MIH5) (1:200 dilution) | BD Biosciences | Cat# 558091, RRID: AB_397018 |
| Anti-mouse FcεRI-PE-Cy7 (clone MAR-1) (1:200 dilution) | BioLegend | Cat# 134326, RRID: AB_2572064 |
| Anti-mouse Mrc1 (CD206)-PE-Cy7 (clone C068C2) (1:400 dilution) | BioLegend | Cat# 141720, RRID: AB_2562248 |
| Anti-mouse FOXP3-FITC (clone FJK-16s) | eBioscience™ | Cat# 11-5773-82, RRID: AB_465243 |
| Anti-mouse IFNγ-APC (Clone XMG1.2) (1:200 dilution) | BD Biosciences | Cat# 554413, RRID: AB_398551 |
| Anti-mouse TNF -PE-Cy7 (Clone MP6-XT22) (1:200 dilution) | BD Biosciences | Cat# 561062, RRID: AB_398553 |
| Anti-mouse Granzyme B-PE (Clone NGZB) (1:200 dilution) | eBioscience™ | Cat# 12-8898-82, RRID: AB_10870787 |
| Anti-mouse iNOS/Nos2 PE (clone CXNFT) (1:200 dilution) | eBioscience™ | Cat# 12-5920-82, RRID: AB_2572642 |
| Zombie Fixable Viability™ Sampler Kit | BioLegend | Cat# 423105 |
| **Chemicals, Peptides, and Gene blocks** | | |
| RPMI-1640 | HyClone | Cat# SH30096.02 |
| Trypsin | Genclone | Cat# 25200056 |

| | | |
|---|---|---|
| Defined fetal bovine serum | HyClone | Cat# SH30070.03HI |
| HBSS | Hyclone | Cat# SH30588.02 |
| PBS | Gibco | Cat# 20012027 |
| Sodium Bicarbonate | Gibco | Cat# 25080094 |
| Sodium Pyruvate | Gibco | Cat# 11360070 |
| L-Glutamine | Gibco | Cat# A2916801 |
| ACK lysis buffer | Gibco | Cat# A1049201 |
| Phorbol-12-myristate-13-acetate (PMA) | MilliporeSigma | Cat# 500582 |
| Ionomycin | Fisher | Cat# BP2527-1 |
| Fugene | Promega | Cat# E2311 |
| pMSCV-IRES GFP | addgene | Cat# 20672 |
| Collagenase Type IA | Sigma-Aldrich | Cat# C9891 |
| Poly(I:C) HMW VacciGrade™ | InvivoGen | Cat# vac-pic |
| Gibson Assembly® Cloning Kit | NEB | Cat# E5510S |
| Mutant Lama4 peptide, sequence VGFNFRTL | Peptide 2.0 | Custom order |
| Mutant Alg8 peptide, sequence ITYTWTRL | Peptide 2.0 | Custom order |
| Mutant Lama4 SLP, sequence QKISFFDGFEVGFNFRTLQPNGLLFYYT | Peptide 2.0 | Custom order |
| Mutant Adpgk SLP, sequence HLELASMTNMELMSSIVHQ | Peptide 2.0 | Custom order |
| Mutant Rpl18 SLP, sequence KAGGKILTFD***R***LALESPK | Peptide 2.0 | Custom order |
| Mutant Dpagt1 SLP, sequence EAGQSLVISASIIVFNL***L***ELEGDYR | Peptide 2.0 | Custom order |
| OVA-I$_{257–264}$ peptide, sequence SIINFEKL | Peptide 2.0 | Custom order |
| Mutant Itgb1 SLP, sequence DDCWFYFTYSVNGY***N***EAIVHVVETPDCP | Peptide 2.0 | Custom order |
| OVA-II$_{323–339}$, sequence ISQAVHAAHAEINEAGR | Peptide 2.0 | Custom order |
| mAlg8-P2A-mItgb1 | IDT | Custom order |

| | | |
|---|---|---|
| CTTCTCTAGGCGCCGGAATTCAGCCACCATGGCAGTG GGCATCACATACACCTGGACCAGGCTGTATGCTTCAGT GTTGACTGGCTCCCTTGTCGGCAGCGGCGCCACAAAC TTCTCTCTGCTAAAGCAAGCAGGTGATGTTGAAGAAAA CCCCGGGCCTGATGACTGCTGGTTCTATTTCACCTATTC AGTGAATGGCTACAATGAAGCTATCGTGCATGTTGTGG AGACTCCAGACTGTCCTTAATACGTAGCTAGCGGATCCCA | | |
| mLama4-P2A-mItgb1 CTTCTCTAGGCGCCGGAATTCAGCCACCATGCAGAAAATA TCTTTCTTTGATGGCTTTGAAGTAGGCTTCAATTTCCGAAC ATTACAGCCAAATGGGTTACTATTCTACTACACAGGCAGCG GCGCCACAAACTTCTCTCTGCTAAAGCAAGCAGGTGATGT TGAAGAAAACCCCGGGCCTGATGACTGCTGGTTCTATTTCA CCTATTCAGTGAATGGCTACAATGAAGCTATCGTGCATGTTG TGGAGACTCCAGACTGTCCTTAATACGTAGCTAGCGGATCCC A | IDT | Custom order |
| **Critical Commercial Assays** | | |
| Fixation/Permeabilization Solution Kit | BD Biosciences | Cat# 555028 |
| Foxp3 / Transcription Factor Staining Buffer Set | eBioscience | Cat# 00-5523-00 |
| Chromium Next GEM Single-cell 5′ Reagent Kit v2 | 10x Genomics | Cat# 100263 |
| Chromium Next GEM Single Cell 5' v2 (Dual Index) | 10x Genomics | Cat# CG000330 |
| Qubit HS dsDNA Assay | ThermoFisher | Cat# Q32851 |
| HS DNA Bioanalyzer | Agilent | |
| **TotalSeq Antibodies** | | |
| CD45 and H-2 MHC class Totalseq™-C0301 anti-mouse Hashtag 1, sequence | BioLegend | Cat# 155861, RRID: AB_2800693 |
| CD45 and H-2 MHC class Totalseq™-C0302 anti-mouse Hashtag 2 Antibody | BioLegend | Cat# 155863, RRID: AB_2800694 |
| CD45 and H-2 MHC class Totalseq™-C0303 anti-mouse Hashtag 3 Antibody | BioLegend | Cat# 155865, RRID: AB_2800695 |
| **Software and algorithms** | | |
| Flow jo_v10.8.1 | | |
| GraphPad Prism version 10 | | |
| Cell Ranger v.7.1.0 | https://support.10xgenomics.com/ single-cell-gene-expression/software | |

| | |
|---|---|
| Seurat R package v.4.3.0.1 | https://satijalab.org/seurat/ |
| scRepertoire v.2.0.0 | https://www.borch.dev/uploads/screpertoire/ |
| ImmGen | https://www.immgen.org |
| ggplot2 | https://ggplot2.tidy verse.org/index.html |
| **Other** | |
| Fortessa X-20 | BD Biosciences |
| LSR Fortessa | BD Biosciences |
| BD FACSAria II | BD Biosciences |
